# Endothelial Knockdown of the Tumor Suppressor, WWOX, Increases Inflammation in Ventilator-Induced Lung Injury

**DOI:** 10.1101/2023.06.29.547087

**Authors:** Zhenguo Zeng, Eltyeb Abdelwahid, Weiguo Chen, Christian Ascoli, Trinh Pham, Jeffrey R. Jacobson, Steven M. Dudek, Viswanathan Natarajan, C. Marcelo Aldaz, Roberto F. Machado, Sunit Singla

## Abstract

**Background:** Chronic cigarette smoke exposure downregulates lung expression of WWOX, an ARDS relevant tumor suppressor. Prior work has revealed a barrier protective function of WWOX during infectious models of ARDS. Proteomic analysis of *WWOX*-silenced lung endothelial cells suggest involvement of WWOX in protection against mechanical stretch-induced inflammation.

**Methods:** Protein lysates from *WWOX*-silenced endothelial cells (ECs) were analyzed using tandem mass tag mass spectrometry (TMT-MS) to determine the differential expression status of the proteome compared to wild type ECs. *WWOX*-silenced ECs as well as those isolated from endothelial *Wwox* knockout (EC *Wwox* KO) mice were subjected to cyclic stretch (18% elongation, 0.5 Hz, 4 hours). Cellular lysates and media supernatant were harvested for assays of cellular signaling, protein expression, and cytokine release. Dual silencing of *WWOX* and zyxin was achieved to determine the role of zyxin upregulation in IL-8 production following mechanical stretch and during *WWOX* knockdown. Control and EC *Wwox* KO mice were subjected to high tidal volume ventilation (VILI, 40ml/kg, 65 breath/min, 4hours). Bronchoalveolar lavage fluid and mouse lung tissue were harvested for cellular signaling, cytokine secretion, and histologic assays.

**Results:** TMT-MS revealed upregulation of zyxin expression during WWOX knockdown which predicted a heightened inflammatory response to mechanical stretch. *WWOX*-silenced ECs and ECs isolated from EC *Wwox* mice displayed significantly increased cyclic stretch-mediated secretion of various cytokines (IL-6, KC/IL-8, IL-1β, and MCP-1) relative to controls. This was associated with increased ERK and JNK phosphorylation but decreased p38 MAPK phosphorylation. EC *Wwox* KO mice subjected to VILI sustained a greater degree of injury than corresponding controls. Silencing of zyxin during *WWOX* knockdown abrogated stretch-induced increases in IL-8 secretion.

**Conclusion:** Loss of WWOX function in ECs is associated with a heightened inflammatory response during mechanical stretch that is associated with increased MAPK phosphorylation and appears to be dependent on upregulation of zyxin.

## Introduction

The WW domain-containing oxidoreductase gene, WWOX, consists of a 1.2 kb exonal product that spans 1.1 Mb of the human genome on chromosome 16q23, making it one of the most active common chromosomal fragile sites in humans [1]. As such, genotoxic environmental exposures can cause homozygous deletions, loss of heterozygosity, and translocation breakpoints at the WWOX locus [2]. This includes exposure to cigarette smoke, and thus Wwox function in the lung is of interest from the standpoint of tobacco-related disease pathogenesis. Previous observations in humans indeed suggest significant downregulation of gene and protein expression of Wwox in the lung following chronic cigarette smoke exposure [3].

The implications of WWOX knockdown for lung disease pathogenesis have been studied in two previous papers. Intratracheal instillation of Wwox-targeting siRNA in mice was observed to result in further potentiation of neutrophilic inflammation in a lipopolysaccharide (LPS)-induced murine model of lung injury [4]. Endothelial cell specific Wwox knockout (EC-Wwox KO) mice stimulated by intratracheal LPS or methicillin-resistant Staphylococcus aureus (MRSA) exhibited significantly greater level of vascular leak and histologic lung injury [3].

These observations are of particular interest in the context of observed associations between prior cigarette smoke exposure and decreased resilience against acute respiratory distress syndrome (ARDS) during conducive conditions. ARDS is a critical illness that occurs following events such as acute aspiration, pneumonia (bacterial or viral), sepsis, blunt force trauma, and non-lung protective mechanical ventilation often required to support severe respiratory failure patients (including ARDS) [5-7]. An increased progression to clinically detectable ARDS has been observed in cigarette smokers versus nonsmokers following blunt force trauma, cardiac surgery, transfusion of blood products, or non-pulmonary sepsis [8-12]. Cigarette smoke worsens human [8, 12, 13] and murine forms of ARDS induced by pulmonary sepsis [14] where it has been observed to exacerbate vascular leak.

The mechanistic underpinning for the observed Wwox-deficient lung phenotype is unknown. In the current study, tandem mass tag-mass spectrometry (TMT-MS) was leveraged to determine the most differentially expressed proteins in human endothelial cells during WWOX knockdown relative to wild type controls. 2051 distinct proteins were identified and quantitated at a 1% false discovery rate. Among the most significant differentially expressed proteins, two were identified which have previously been biologically connected to the inflammatory and endothelial barrier-susceptible phenotypes observed in prior work. These were the mechano-transducer protein, zyxin, which was upregulated during WWOX knockdown, and the mitochondrial superoxide dismutase, SOD2, which was downregulated. Zyxin is a known modulator of inflammatory responses in endothelial cells following mechanical stretch [15]. Along with known interactions between WWOX and the MAP kinases [4, 16], which are also involved in transducing gene transcriptional responses to stretch, these observations led to an examination of the potential involvement of Wwox in VILI-associated inflammation.

## Materials and Methods

### Reagents

FITC-Dextran, RBC Lysis Buffer, DMEM, bovine serum albumin, DNase, Dispase II, and RIPA Buffer were purchased from Sigma-Aldrich. BCA Protein Assay Kit, Shandon Kwik-Diff kit, siPORT Amine transfection reagent, Lipofectamine 2000, High Capacity cDNA Reverse Transcription kit, Taqman Gene Expression Master Mix, and Taqman Gene Expression Assays for GAPDH and WWOX were from Thermo Fisher. Mini-PROTEAN® TGX™ Precast Gels were from BioRad. QIAShredder and RNeasy Mini kits were from Qiagen. Anti-WWOX antibody was from abcam (Cambridge, MA). Human lung microvascular endothelial cells (ECs), and endothelial cell growth media were purchased from Lonza (Basel, Switzerland). Heparin was purchased from Hospira (Lake Forest, IL). Collagenase was from Worthington (Lakewood, NJ). ELISA kits for mouse cytokines (KC, MCP-1, IL-6, IL-1β), anti-mouse CD16/32, anti-mouse CD31-PE/Cy7, and anti-mouse CD45-Alexa700 antibodies were from BioLegend (San Diego, CA) and R&D systems (Minneapolis, MN). siGENOME WWOX-targeting and scrambled control siRNA was from Dharmacon/GE, and the zyxin siRNA was from Santa Cruz Biotechnology (Santa Cruz, CA, USA). pCMV entry vector containing the ORF corresponding to (Myc-DDK-tagged)-human WW domain-containing oxidoreductase (Wwox), transcript variant 1, as well as a (Myc-DDK-tagged)-empty vector were purchased from OriGene. Anti-zyxin antibody was from Thermo Fisher Scientific, Inc., (Waltham, MA, USA).

### Cell culture

Human pulmonary artery endothelial cells (ECs) obtained from Lonza were grown in cell culture using endothelial growth media containing supplemental growth factors and 2% fetal bovine serum according to the manufacturer’s instructions. Cells underwent a maximum of three passages for use in experiments.

For separate studies, ECs were isolated from the lungs of 8 weeks old C57BL/6-Wwoxflox/flox and tamoxifen-treated Cdh5-CreERT2/Wwox^flox/flox^ mice as in Kawasaki et al [17]. Following ketamine/xylazine administration, the thoracic cavity was surgically opened. 20 ml of sterile PBS containing 10U/ml heparin was instilled into the pulmonary vasculature via injection through the right ventricle until the lungs were blood-free. Mouse lungs were then harvested, minced, and digested in an enzyme complex as previously described. Washed cell pellets were resuspended and serially incubated with anti-mouse CD16/32 (to block Fc receptors), anti-mouse CD31-PE/Cy7, and anti-mouse CD45-Alexa700 antibodies. After washing with PBS/BSA, cells were sorted by flow cytometry (MoFlo Astrios Cell Sorter, Beckman Coulter, Indianapolis, IN) to isolate CD31+CD45-ECs which were then placed in EC growth media on gelatin coated plates.

### Mechanical Cyclic Stretch of Endothelial Cells

ECs were plated onto six-well Flexcell plates which were coated with type I collagen (FlexCell international, Hillsborough, NC) and grown to 75-80% prior to transfection of siRNAs as described below. 3 daysafter transfection of siRNA, mechanical stretch was performed on the FlexCell Strain Unit (FX-3000, FlexCell International). The device cyclic stretch cells by a vacuum system. The stretch condition was 18% elongation at frequency of 30 cycles per minute (0.5Hz) for 4 hours. After finishing cyclic stretch, the medium was harvested for ELISA assay and the cell lysates were harvested with cell lysate buffer for western blot [16].

### Animals

All experiments and animal care procedures were approved by the University of Illinois at Chicago Animal Care and Use Committee. C57BL/6-Wwox^flox/flox^ mice were donated generously by Dr. C. Marcelo Aldaz, MD/PhD of MD Anderson Center at the University of Texas. C57BL/6-Cdh5-CreERT2 mice were purchased from Jackson Laboratories. The two strains were crossed to produce Cdh5-CreERT2/Wwox^flox/flox^ mice as in Ludes-Meyers et al[18]. These are a tamoxifen-inducible, endothelial cell-specific Wwox knockout mouse strain. They were housed in cages in a temperature-controlled room with a 12 hrs dark/light cycle, and with free access to food and water.

### Murine High Tidal Volume Ventilation

6 week old male Cdh5-CreERT2/WWOX^flox/flox^ mice and control animals received intraperitoneal injections of tamoxifen (75mg/kg) for 5 consecutive days. 7 days later they were subjected to high tidal volume mechanical ventilation (VT=40ml/kg, 65 breaths/min) for 4 hours to induce VILI as we have previously described. Mice were sacrificed and bronchoalveolar lavage (BAL) were harvested. BAL fluid was assessed for total cells counts, protein concentration, and cytokine levels. Meanwhile, the mouse lungs were flushed and harvested for histological analysis [16].

### Bronchoalveolar lavage fluid (BALF) collection and analysis

Mice were anesthetized with ketamine/xylazine. The trachea was exposed surgically, and a small incision was made on the anterior surface. An 18-gauge blunt-end cannula was inserted into this opening and secured with a suture tied around the trachea. 1 ml of sterile saline was infused through this cannula and slowly withdrawn. This BALF was centrifuged to pellet cells and remove supernatant. RBCs were removed from the cell pellet using RBC lysis buffer according to the manufacturer’s instructions. The remaining cells (leukocytes) were pelleted, resuspended in a fixed volume of saline, and a small aliquot was used to measure cell concentration with an automated cell counter. The remaining cell suspension was used to make a cytospin prep on a glass slide for staining with the Shandon Kwik-Diff kit. A manual differential cell count was performed on 10 hpfs to determine relative percentages of neutrophils in the leukocyte population. Protein concentration was measured using the BCA Protein Assay kit, and the corresponding ELISA kits were used to measure concentrations of KC, MIP-2, MCP-1, IL-6, IL-1β, TNF-α.

### Mouse Lung Endothelial cells for RT-PCR, Western blotting, and Histology

Following BALF collection, the right ventricle was infused with sterile saline until the lungs were free of intravascular blood. The right middle lobe was excised and snap frozen in liquid nitrogen for RT-PCR, and Western blotting. The remaining lobes were excised and placed in 10% formalin for 24 hours followed by transfer to 70% ethanol for storage until paraffin embedding and sectioning for H&E staining.

For RT-PCR, lungs were thawed, lysed, homogenized, and RNA was isolated using the QIAShredder and RNeasy Mini kits per manufacturer’s instructions. cDNA was produced for each sample using the high-capacity cDNA Reverse Transcription kit from ThermoFisher Scientific. Real-time PCR was performed using the Taqman Gene Expression Master Mix and Assays per manufacturer’s instructions. Relative quantification between controls and experimental samples of Wwox gene expression normalized for GAPDH was determined using the ΔΔC_t_ method.

For Western blotting, snap frozen lungs were thawed and homogenized in RIPA buffer. Homogenized samples were centrifuged to pellet debris and isolate protein lysates. Protein concentration of lysates was measured using the BCA Protein Assay kit, and samples were diluted in RIPA buffer until all had the same protein concentration. After dilution with SDS buffer, samples were loaded into 4-20% Mini-PROTEAN® TGX™ Precast Gels and subjected to SDS-PAGE. Gels were transferred to nitrocellulose membranes which were then subjected to Western blotting using antibodies targeting proteins of interest in accordance with the manufacturer’s instructions. Densitometric analysis was performed using ImageJ software.

### Human or Mouse Endothelial Immunochemistry Assay

Human endothelial cells were grown to 75% confluence on glass slides in 12 well-plates, the cells were silenced with 100nM of human scrambled control or *WWOX-*targeting siRNA for 3 days. The cells were fixed with paraformaldehyde followed by 0.1% triton incubation. Cells were then incubated with 1% BSA for 30min followed by overnight incubation with the antibodies indicated. On the next day, the cells were incubated with secondary antibodies for 1 hr. After washing with PBS, the slides were mounted with DAPI mount solution. For mouse endothelial cells,ECWX-flox or ECWX-KO endothelial cells were seeded on the slides in 12 wells overnight. On the next day the cells underwent the same procedure as for human endothelial cells but stained with the Wwox antibody or VE-cadherin antibodies indicated. After slides dried, fluorescent images were taken under Olympus BX51 Fluorescence Microscope [19].

### In vitro silencing of Wwox and Zyxin in human pulmonary endothelial cells

In vitro silencing of Wwox in human ECs (Lonza) was achieved using transfection of siRNA as described in Wolfson, et al[20]. WWOX-targeting (CCAAGGACGGCUGGGUUUA) and/or Zyxin -targeting (GUGUUACAAGUGUGAGGACTT) versus scrambled control siRNA (UAGCGACUAAACACAUCAA) was complexed with siPORT Amine transfection reagent. Serum free media was used to bring the concentration of siRNA to 100 nM. In experiments in which dual silencing of Wwox and Zyxin were achieved, 45 nM concentrations of each siRNA were used together. Transfection of ECs was then performed as described previously [20]. 72 hours after transfection, ECs were analyzed by Western blotting for WWOX and zyxin expression.

### Proteomic Analysis

TMT-MS was used to obtain relative quantitation of protein expression between control and WWOX-silenced human ECs as described previously [21-23]. Briefly samples were digested, and 50 micrograms of peptides from each condition were labelled with TMT tags, combined, fractionated, and desalted with C18 spin columns. The labelled peptides were subjected to MS analysis and run in triplicate. Data were analyzed against a human database using Scaffold 5. In total, 2051 proteins, including 20 decoys, were identified. Differential protein expression between groups was assessed comparing mean normalized intensity values utilizing Scaffold’s unpaired t-test and considered significant upon correction for multiple testing at a Benjamini-Hochberg false discovery rate <1%. Log_2_ fold change was defined as the difference of log_2_ normalized intensity values between *WWOX-*silenced human ECs and controls.

### Statistical analysis

A Mann-Whitney U test was performed to evaluate comparisons between groups using GraphPad Prism software, where statistical significance was defined as p<0.05.

## Results

### TMT-MS Proteomics and Validation Studies Identified Zyxin as a Significant Differentially Expressed Protein in WWOX-silenced ECs compared to wild-type Controls

TMT-MS was used to profile the proteome of control versus *WWOX*-silenced ECs. Silencing was confirmed by Western Blotting and immunostaining (Figures 1A and 1B). Results of the proteomic analysis are shown in Figure 1C and Supplemental Table 1. Of the most significant differentially expressed proteins were the mechanotransducer, zyxin, and the mitochondrial superoxide dismutase, SOD2. Validation studies for zyxin were performed using RT-PCR and Western Blotting which reproduced the TMT-MS result at both the mRNA and protein levels (Figures 1D, 1E).

**Figure 1.**
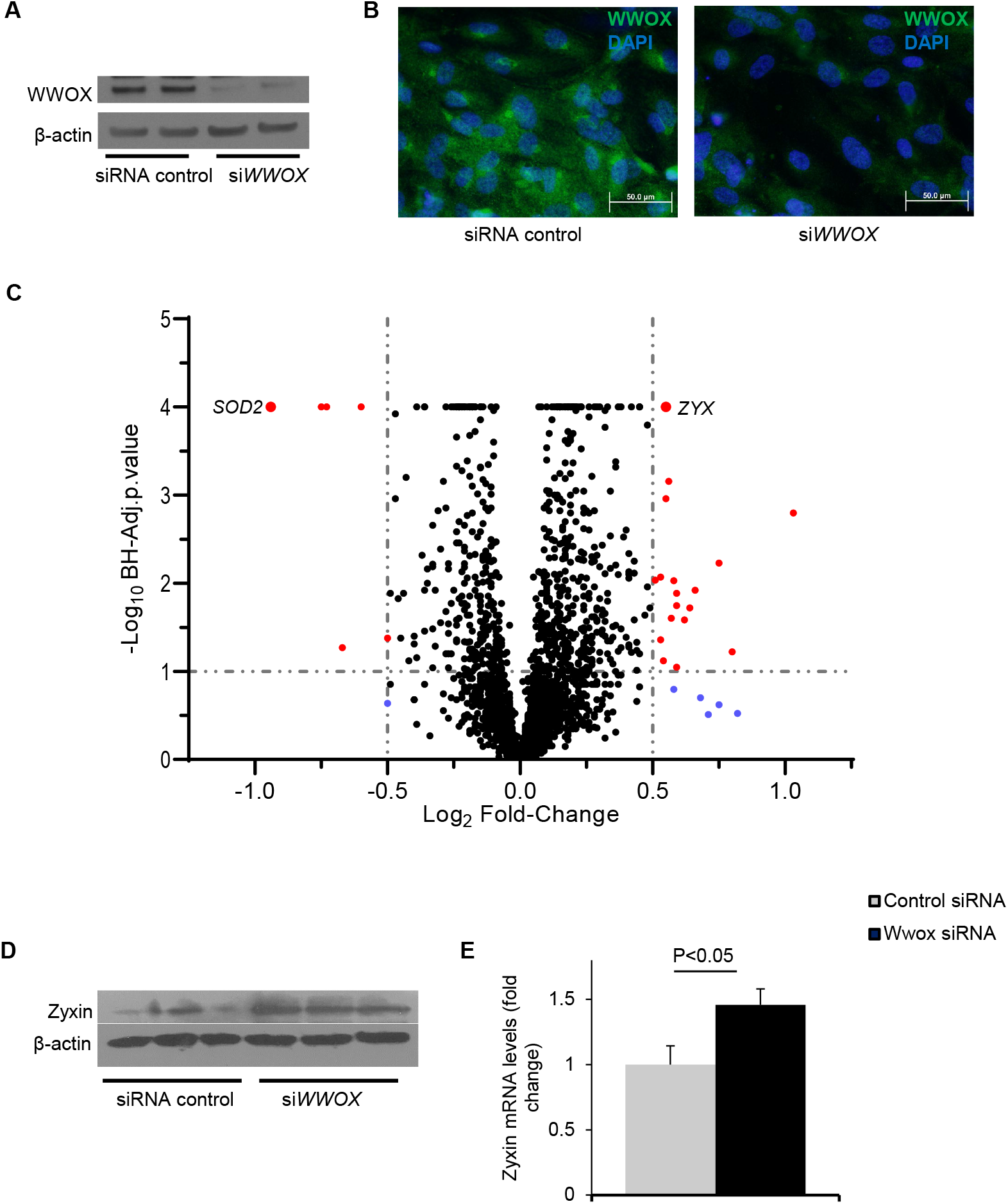
*WWOX* Knockdown is Associated with Upregulation of the Mechanosensitive Transducer, Zyxin. **A**. ECs were transfected with either scrambled, control or *WWOX*-silencing siRNA. Western blotting illustrates the degree of *WWOX* knockdown achieved. **B**. Immunostaining for WWOX confirms knockdown of the predominantly cytosolic protein in siRNA-transfected ECs. **C**. TMT-MS differential protein expression between *WWOX*-silenced and control ECs is depicted for uniquely identified proteins on the volcano plot and is detailed in Supplemental Table 1. Horizontal dotted line indicates the t-test’s significance threshold (Benjamini-Hochberg adjusted p-value < 1%). Vertical dotted lines indicate log_2_ fold change of -0.5 or 0.5. SOD2 (*superoxide dismutase 2*) and ZYX (*zyxin*) were among the most significant differentially expressed proteins and are labelled for reference. **D**. Zyxin upregulation during *WWOX* knockdown was confirmed by Western blotting.**E**. qPCR for zyxin revealed that the upregulation occurs at the mRNA level (n=6, mean +/- SEM, p<0.05)

### Loss of WWOX promotes EC Cytokine Production in response to Cyclic Stretch

18% elongation cyclic stretch (CS) of *WWOX*-silenced endothelial cells was used to mimic high tidal volume ventilation in vitro. After 4 hours of cyclic stretch (18% elongation, 0.5Hz), cytokine assays done on EC culture media showed that silencing of *WWOX* significantly increased CS-mediated production of IL-6 (239.5± 21.8 vs 113.77±5.0 pg/mL, n=6, P<0.05, si*WWOX* vs siRNA control) and IL-8 (128.42± 3.11 vs 76.10±3.5 pg/mL, n=6, P<0.05, si*WWOX* vs siRNA control) (Figure 2).

**Figure 2.**
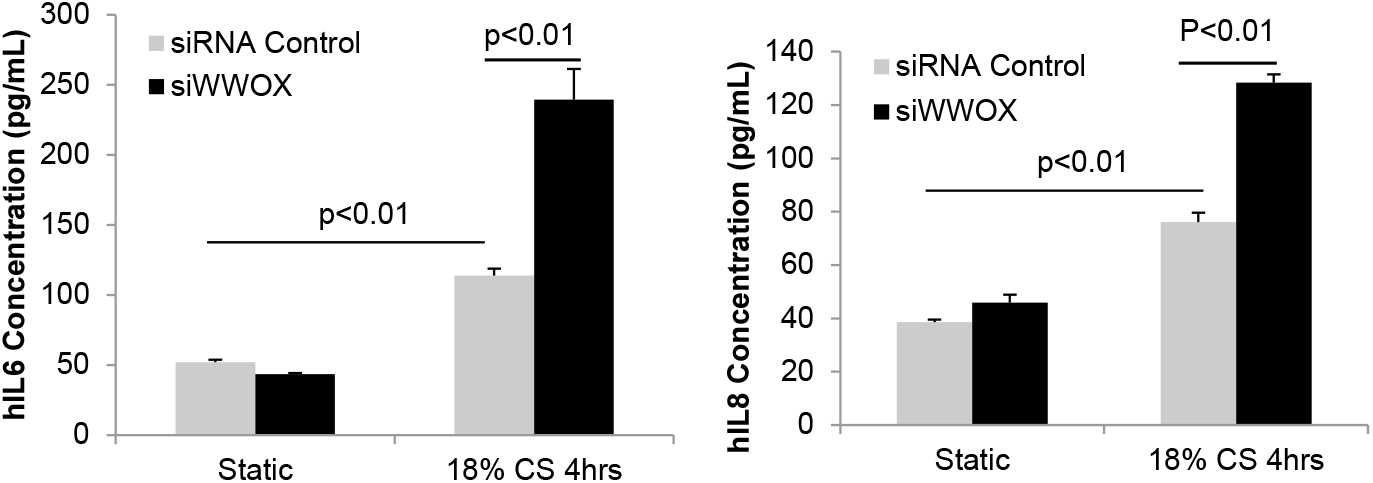
Silencing of *WWOX* enhanced endothelial inflammatory response to cyclic stretch. Human ECs were seeded on a Flexcell plate and silenced with siRNA control or siRNA of *WWOX* for 3 days. After cyclic stretch, the media were harvested for ELISA. IL-6 and IL-8 production increased significantly in control ECs subjected to stretch compared to static controls (n=6, mean +/-SD, p<0.01). Compared to the stretched ECs transfected with siRNA control, IL-6 and IL-8 production from stretched *WWOX*-silenced ECs was significantly increased (n=6, mean +/- SD, p<0.01).

MAPK signaling is a well known effector pathway of mechanosensitive gene expression changes in ECs [24]. Therefore, ERK and JNK phosphorylation status were examined in *WWOX*-silenced endothelial cells. As shown in Figure 3, CS induced time-dependent ERK phosphorylation in control siRNA transfected endothelial cells with peak phosphorylation occurring at 5 minutes. However, silencing of *WWOX* induced an increase in basal and CS-mediated ERK phosphorylation compared to control siRNA transfected endothelial cells (Figure 3 upper panel). Similarly, silencing of *WWOX* also induced increase of basal and CS-mediated JNK phosphorylation compared to control siRNA transfected endothelial cells (Figure 3 second panel). Interestingly, silencing of *WWOX* significantly decreased basal and CS-mediated p38 MAPK phosphorylation compared to controls (Figure 3 third panel).

**Figure 3.**
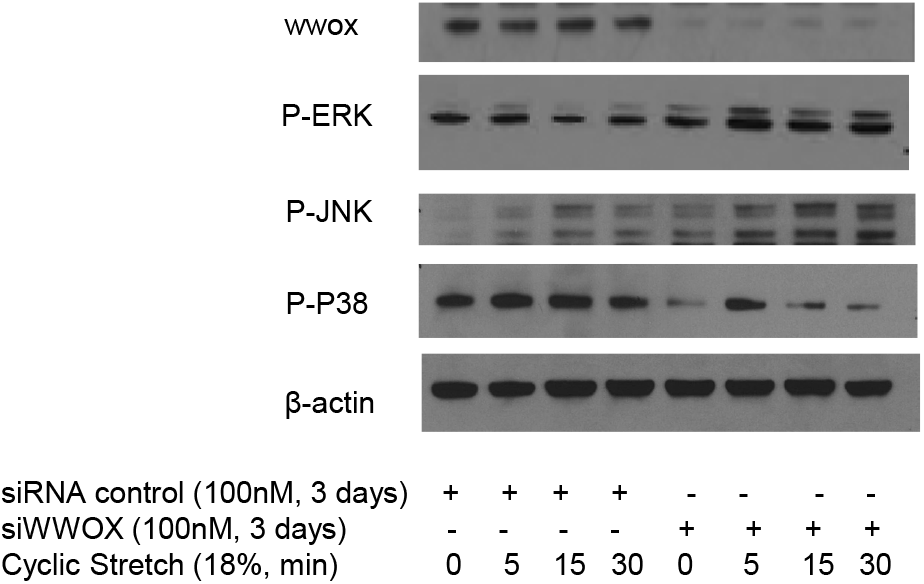
Silencing of *WWOX* enhanced MAPK activity in endothelial inflammatory response to cyclic stretch. Human ECs were seeded on a Flexcell plate and silenced with siRNA control or siRNA of *WWOX* for 3 days. After cyclic stretch, cellular lysates were harvested for Western blotting. 18% elongation 0.5 Hz cyclic stretch induced ERK and JNK and p38 phosphorylation in both control and *WWOX*-silenced ECs. Silencing of *WWOX* induced significantly higher ERK and JNK phosphorylation and lower p38 phosphorylation during both basal and cyclic stretch conditions.

### EC *Wwox* KO mice exhibit more injury during VILI

To further investigate the role of WWOX in VILI, *Wwox* flox/flox (control) mice and EC *Wwox* KO mice underwent high tidal volume ventilation for 4 hours (40ml/kg, 65 breath/min). BAL protein and cell counts assay indicated that EC *Wwox* KO mice had significantly increased BAL protein concentration (0.87± 0.21 vs 0.55±0.095 mg/ml, n=5, P<0.05, ECWX -KO vs ECWX-Flox) and cell infiltration (1.62 ± 0.21×10^5^/mL vs 0.93±0.20×10^5^/mL, n=5, P<0.05, ECWX -KO vs ECWX-Flox) (Figure 4A). H&E staining of lung sections showed VILI-induced cell infiltration in both ECWX-Flox and ECWX -KO lung interstitium. Consistent with BAL cell counts, VILI induced significantly higher cell infiltration in ECWX -KO lung compared to ECWX-Flox (Figure 4B).

**Figure 4.**
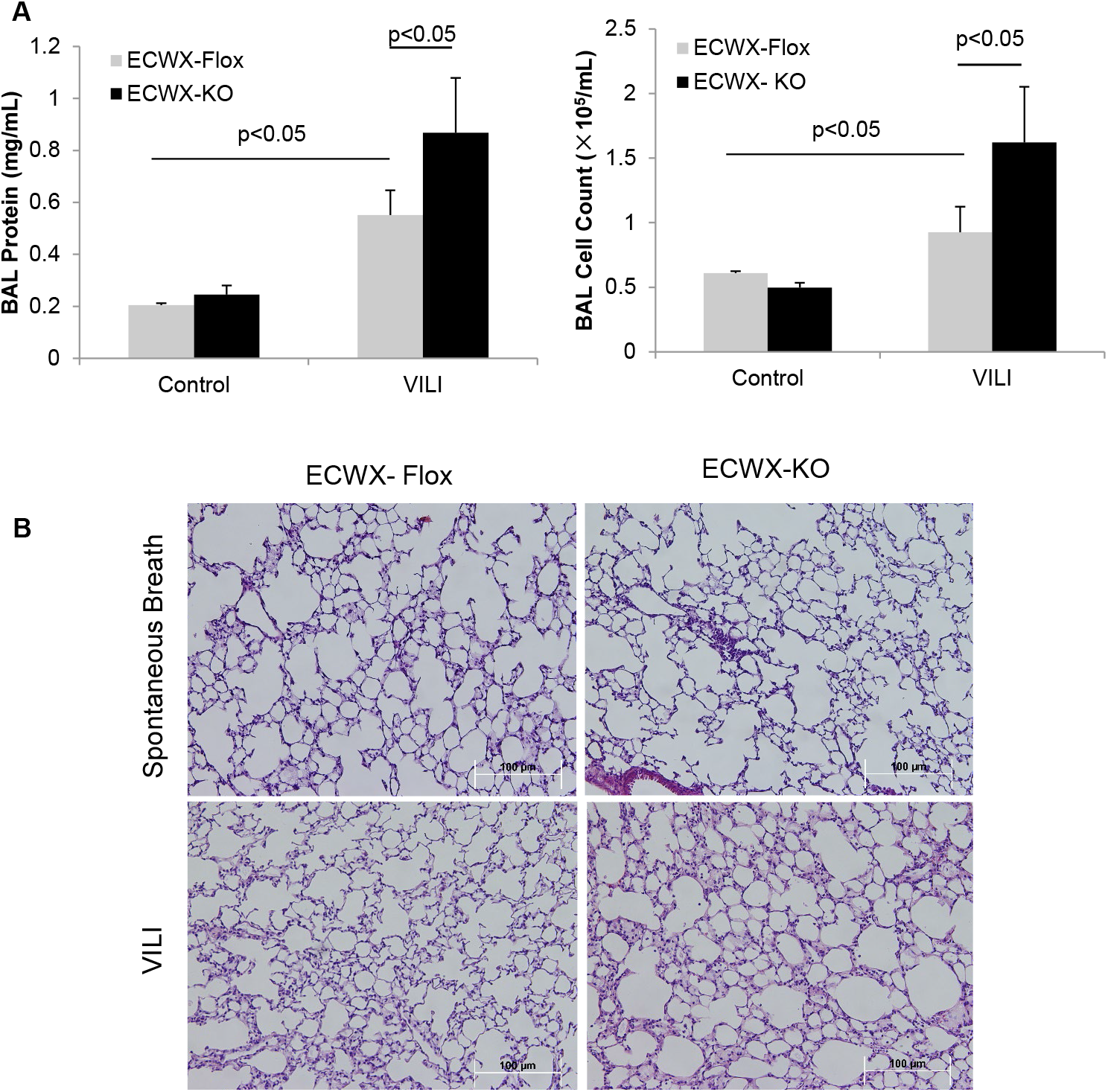
Endothelial-specific *Wwox* KO mice exhibit greater vascular leak during high tidal volume ventilation. EC *Wwox* KO and control mice were subjected to high tidal volume mechanical ventilation as described in materials and methods. **A**. VILI induced significantly higher BAL protein concentrations and cell counts in control mice (n=5, mean +/-SD, p<0.05). Significantly higher protein concentration and cell counts were present in BAL fluid from EC *Wwox* KO mice subjected to VILI compared to corresponding controls (n=5, mean +/- SD, p<0.05). **B**. H&E stain of sections of lung tissue showed increased VILI-induced interstitial and alveolar cellular infiltration in EC *Wwox* KO mice lung compared to controls.

Increased BAL protein and alveolar cellular infiltration in ECWX-KO vs. ECWX-Flox mice after VILI correlated with increased BAL IL-6, KC, IL-1β and MCP-1 production in WXEC-KO mice groups (Figure 5).

**Figure 5.**
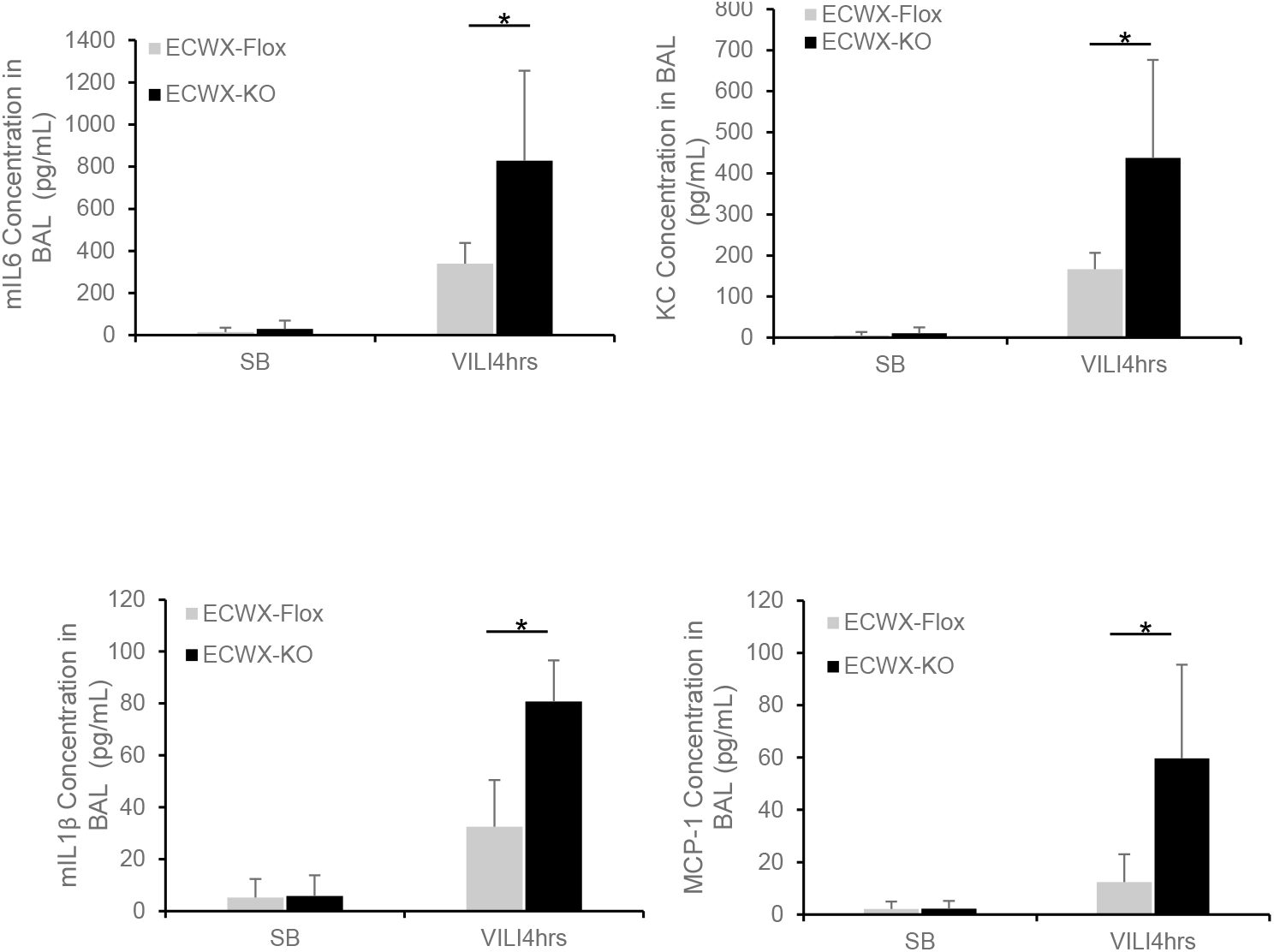
Endothelial-specific *Wwox* KO mice exhibit greater inflammatory response to high tidal volume ventilation. BAL cytokines in VILI-exposed EC *Wwox* KO and control mice were measured by ELISA. These showed that VILI induced significantly higher IL6, KC, IL-1β and MCP-1 generation in EC *Wwox* KO mice (*n=5, mean +/- SD, p<0.05).

### ECs Isolated from EC *Wwox* KO Mice Produce more Cytokines following Cyclic Stretch

Endothelial cells were isolated from ECWX Flox (control) and ECWX-KO mice as described previously [3]. Flow cytometry for CD31+CD45-cells was performed, and these endothelial cells were then cultured on gel coated plates.

Cyclic stretch-mediated cytokine generation from isolated WXEC-flox and WXEC-KO endothelial cells was measured. As shown in Figure 6, 4 hours of 18% elongation cyclic stretch stimulated production of IL-6, KC, IL-1β and MCP-1 in WXEC-Flox endothelial cells medium compared to static control. Elevations of all these cytokines were greater in WXEC-KO ECs undergoing stretch when compared to CS-mediated elevations in WXEC-Flox cells: IL-6 (16.43±11.59 pg/ml vs 44.17±13.47 pg/ml, p<0.05); KC (16.08±3.76 pg/ml vs 65.67±17.84 pg/ml, p<0.05); IL-1β (13.43±17.14 pg/ml vs 38.28±8.97 pg/ml, p<0.05) and MCP-1 (41.12±29.82 pg/ml vs 90.02±10.46 pg/ml, p<0.05) (Figure 6).

**Figure 6.**
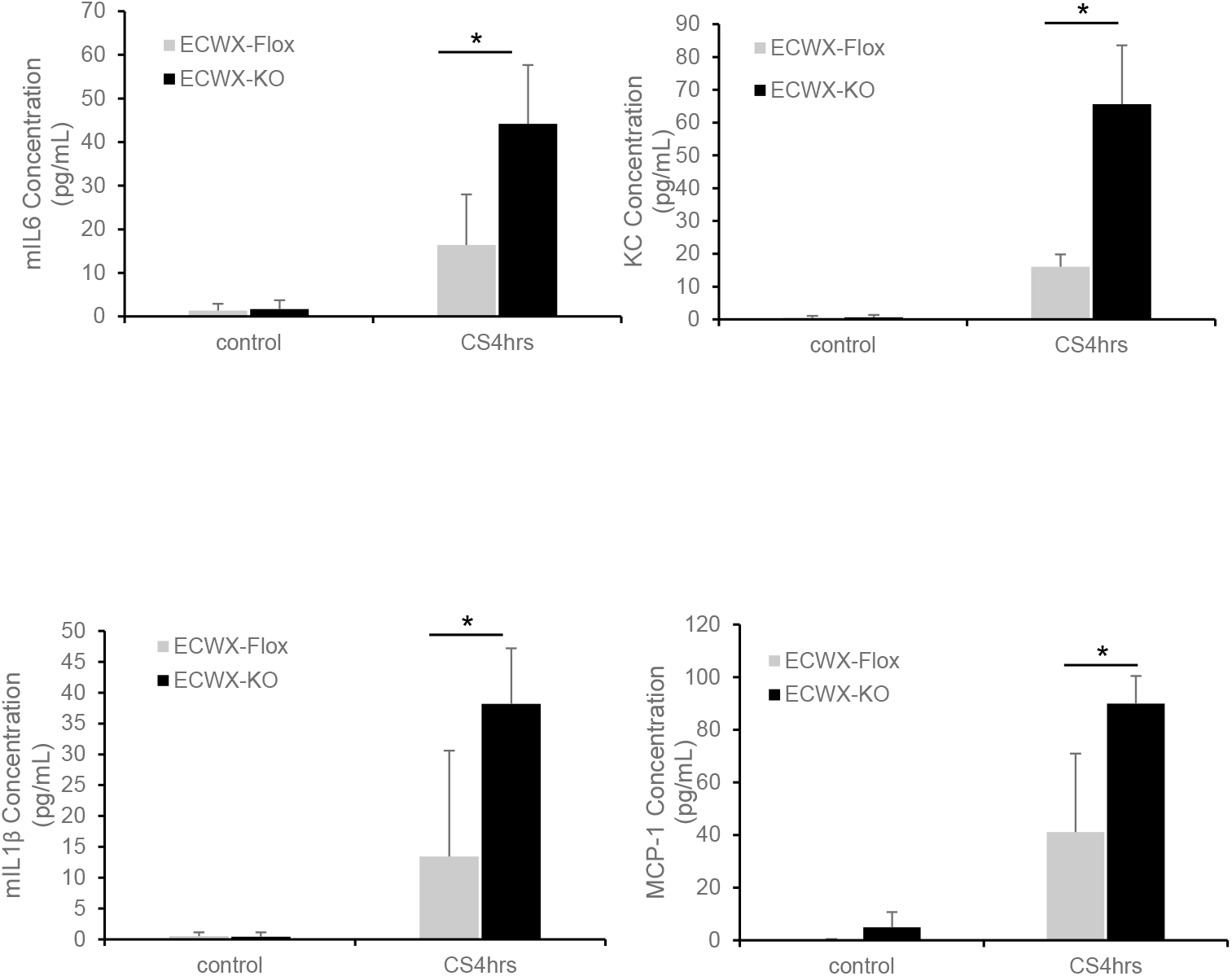
Endothelial cells derived from EC *Wwox* KO mice exhibit greater cyclic stretch-induced cytokine generation. ECs were isolated from EC *Wwox* KO mice and controls. These were grown on FlexCell plates subjected to cyclic stretch. After 4 hours, the medium was harvested for ELISA-based measurement of cytokines. ECs from KO mice exhibited significantly higher IL6, KC, IL-1β and MCP-1 generation following cyclic stretch compared to controls (*n=5, mean +/- SD, p<0.05).

### Silencing of Zyxin during WWOX Knockdown in ECs Mitigates Stretch-Induced Increases in IL-8 Production

The mechanotransducer, zyxin, has previously been shown to be an enhancer for IL-8 transcription during cyclic stretch of ECs [15]. To determine the relative contribution of zyxin upregulation to the increases in stretch-induced increases in IL-8 production during WWOX knockdown, co-silencing of WWOX and zyxin in human ECs was utilized. Silencing of zyxin during WWOX knockdown resulted in a 35% reduction in CS-induced IL-8 secretion compared to ECs in which WWOX was solely silenced (Figure 7).

**Figure 7.**
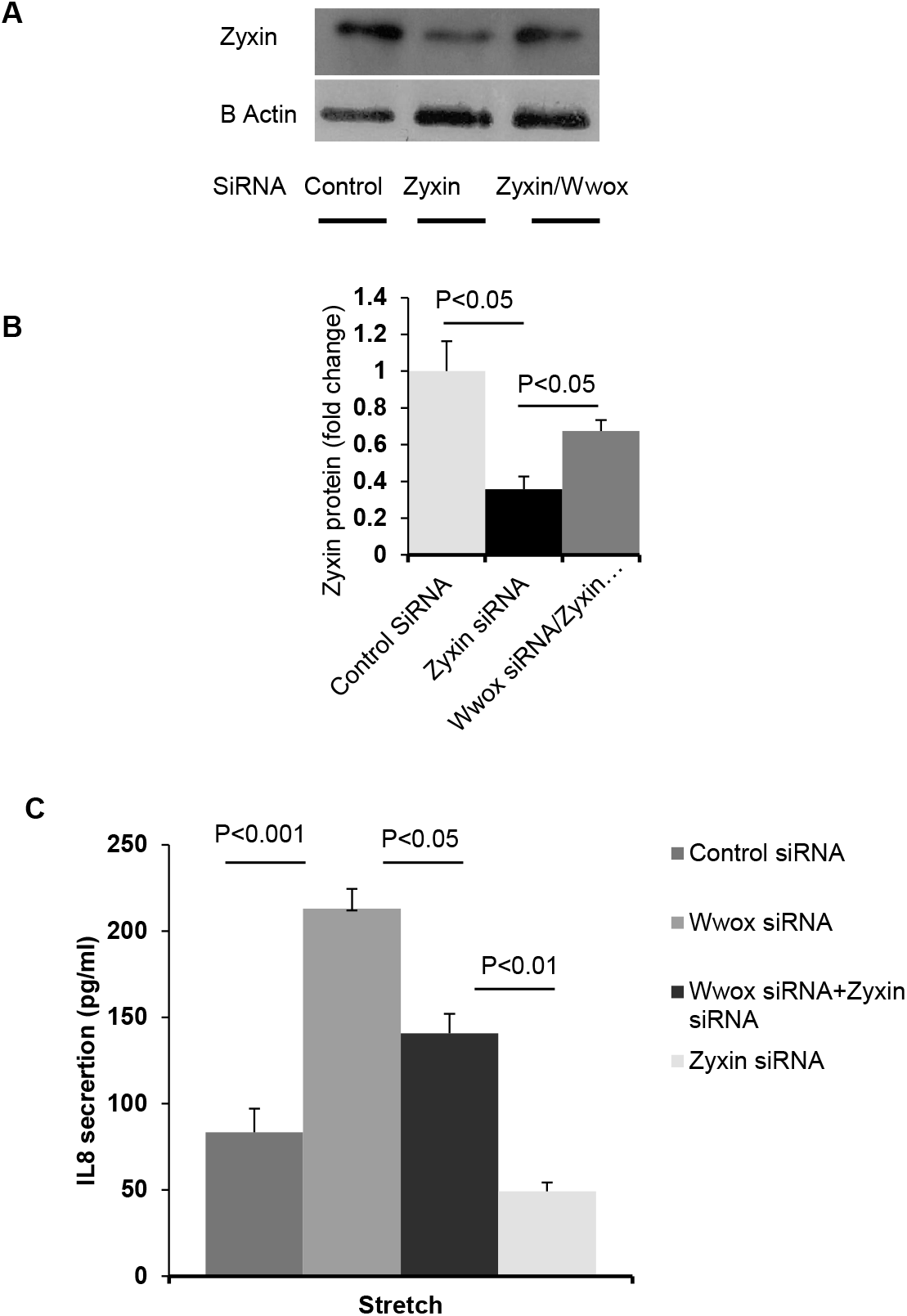
Silencing of Zyxin during *WWOX* knockdown decreases IL-8 production following cyclic stretch. **A**. Human ECs were treated with control, *WWOX*-targeting, and zyxin-targeting siRNA. Dual-silencing of *WWOX* and zyxin was achieved with treatment involving both siRNAs. A representative Western blot of lysates is shown. **B**. Densitometric analysis of several blots from these experiments is reported in the bar graph (n =4, mean +/- SEM, p<0.05). **C**. Following cyclic stretch, culture medium was harvested for measurement of IL-8 generation. The greater amount of IL-8 generation exhibited by *WWOX*-silenced ECs compared to controls following cyclic stretch appears to be abrogated by zyxin knockdown. (n =3, mean +/- SEM, p p<0.05).

### Silencing of WWOX Increases Nuclear Localization of Zyxin

An assessment of zyxin nuclear localization during WWOX knockdown, necessary for zyxin’s IL-8 enhancing function, was performed. As shown in Figure 8, silencing of WWOX was associated with significantly increased nuclear localization of zyxin.

**Figure 8.**
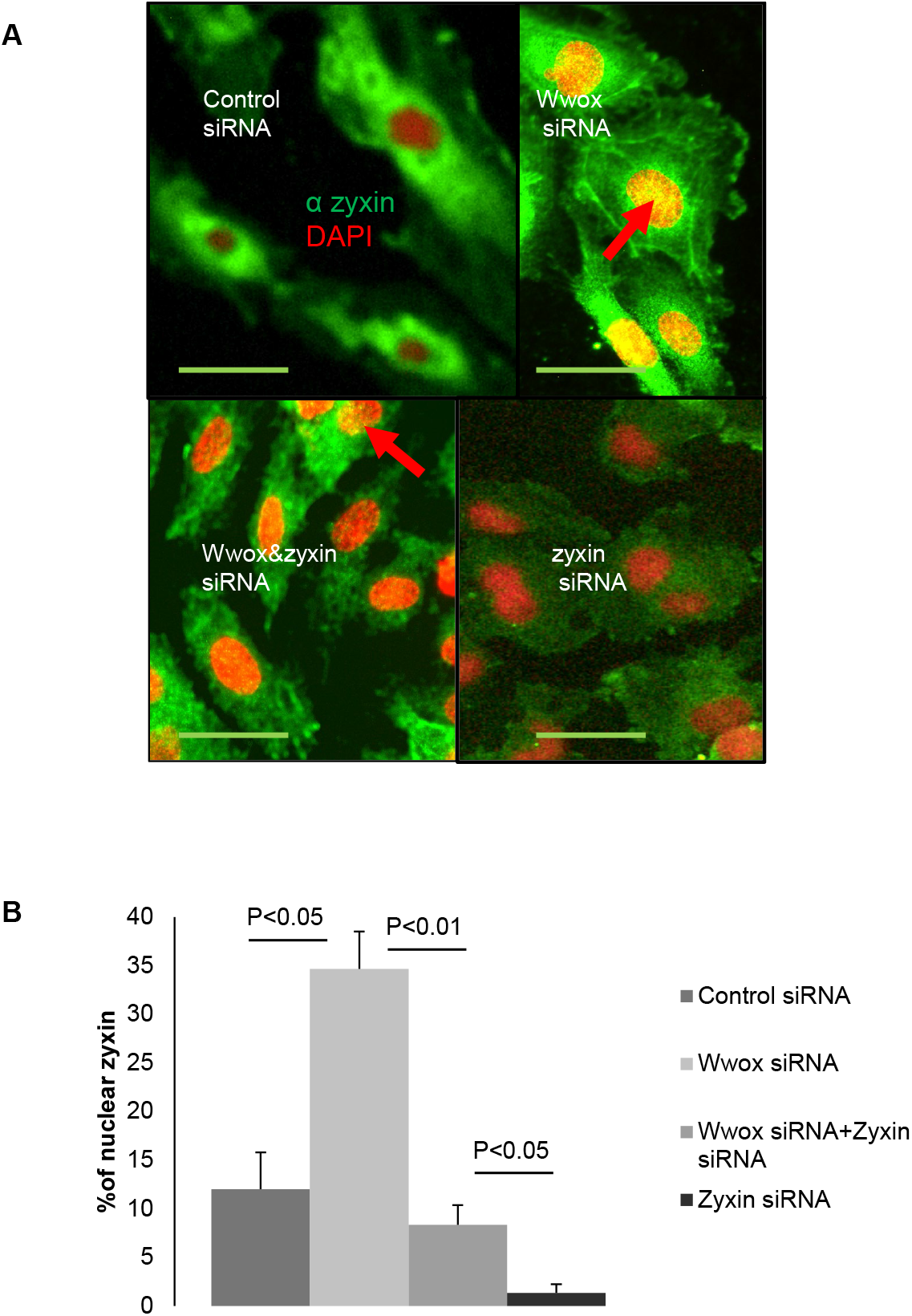
Increased zyxin nuclear localization upon *WWOX* knockdown. ECs were treated with control siRNA, *WWOX* siRNA, *WWOX* and zyxin siRNA, and zyxin siRNA and labeled with anti-zyxin antibody as well as with the nuclear dye DAPI. The percentages of cells with zyxin nuclear staining in each condition were counted. Scale bar=20 μm. (*n=3, mean +/- SEM, p<0.05). Arrow indicates nuclear localization of zyxin.

## Discussion

In this study, observations of the WWOX-deficient lung phenotype, which is associated with tobacco smoke exposure, have been extended to high tidal volume ventilation-mediated acute lung injury. Silencing of the *WWOX* gene in ECs results in a pro-inflammatory, barrier sensitive phenotype characterized, in part, by the upregulation of the mechanosensitive IL-8 enhancer, zyxin. Furthermore, the transduction of signals in ECs following mechanical stress occurs through focal adhesion complexes via mitogen-activated kinases (MAPK). These signaling pathways appear to be potentiated in WWOX-deficient ECs with increases in both CS-mediated ERK and JNK phosphorylation. EC *Wwox* KO mice subjected to a VILI model exhibited, relative to VILI-exposed wild type mice, increased BAL protein, accumulation of alveolar cellular infiltrates, and pro-inflammatory cytokines (IL6, KC, IL-1β and MCP-1). Isolated endothelial cells from EC-*Wwox* KO mice subjected to CS resulted in significantly higher production of cytokines when compared to stretched mouse EC-*Wwox* flox endothelial cells.

In pursuing the mechanisms underlying the WWOX-deficient endothelial cell phenotype, TMT-MS for analysis of differential protein expression between WWOX-silenced ECs and control cells was performed. Two significant differences in protein expression were identified. The mitochondrial superoxide dismutase, SOD2 was markedly downregulated in WWOX-silenced ECs. SOD2 converts mitochondrial superoxide to hydrogen peroxide [25]. In human ECs, loss of SOD2 such as that which occurs in sickle cell disease, is associated with decreased mitochondrial potential without impairment in respiration/ATP production and increased EC layer permeability [25]. Lipopolysaccharide (LPS) or bacterial endotoxin, an ARDS relevant agonist, is known to increase EC barrier permeability, at least in part, by increase in mitochondrial ROS and subsequent VE-Cadherin phosphorylation [26]. Therefore, the loss of SOD2 during WWOX knockdown is currently being studied in the context of LPS- and other bacterial infection-related stimuli in ECs.

The other significant protein expression profile finding in WWOX-silenced ECs is upregulation of the mechanotransducer, zyxin. Zyxin is primarily located in focal adhesion plaques but can shuttle to actin stress fibers and the nucleus in response to mechanical stimulation [27]. Nuclear translocation of zyxin has been shown to be associated with gene expression changes, including upregulation of IL-8 during mechanical stretch of ECs [15, 27]. In the current study, it appears that zyxin upregulation during WWOX-silencing plays a role in the exaggerated IL-8 response of WWOX-deficient ECs during mechanical stretch and VILI. The connection between zyxin and WWOX remains unknown but is the subject of ongoing study. One possibility is via the known direct inhibitory interaction between SMAD3 and WWOX [28], and SMAD-3 dependent upregulation of zyxin [29]. The relationship between increased MAPK signaling and zyxin upregulation during mechanical stretch of WWOX-deficient ECs is also undergoing dissection in ongoing studies.

The pathological change of VILI includes overdistention of alveoli from high tidal volume and repetitive opening and closing of alveoli, with resultant inflammatory response leading to disruption of the endothelial barrier and interstitial and alveolar infiltration of monocytes and neutrophils [30]. Why some patients sustain VILI and recover quickly and others do not may be, in part, related to chronic gene expression changes associated with past exposures such as cigarette smoke. WWOX downregulation appears to be one of these changes with a now-defined impact on some of the molecular mechanisms of stretch-induced inflammatory response in ECs. Further studies will focus on the clinical relevance of these findings related to WWOX versus other ARDS-relevant gene expression changes associated with chronic tobacco product exposure.

## Acknowledgments

This work was supported by Natural National Science Foundation of China (81870011, 81460015 and 8116023) to ZZ; R01-HL-127342, R01-HL-133951, 2R01 HL111656-06 to RFM; 1K08 HL140222-01A1 to SS.

