## Supplemental Table 1 for "Endothelial Knockdown of the Tumor Suppressor, WWOX, Increases Inflammation in Ventilator-Induced Lung Injury"

| Row | Protein | UniProtKB/Swiss-Prot | BH.adj.P.val | Neg.Log.10.BH.adj.P.val | Log2.FoldChange (Treatment - Reference) | Log2.Normalized.Intensity (Controls) | Log2.Normalized.Intensity (WWOX-silenced human ECs) |
|---|---|---|---|---|---|---|---|
| 1 | AAMDC | Q9H7C9 | 0.77 | 0.113509275 | -0.03 | 16.94 | 16.91 |
| 2 | AARS | P49588 | 0.044 | 1.356547324 | -0.07 | 16.69 | 16.62 |
| 3 | AASDHPPT | Q9NRN7 | 0.13 | 0.886056648 | 0.01 | 14.18 | 14.2 |
| 4 | ABCC1 | P33527 | 0.099 | 1.004364805 | 0.18 | 16.2 | 16.37 |
| 5 | ABCD3 | P28288 | 0.11 | 0.958607315 | -0.21 | 15.71 | 15.5 |
| 6 | ABCE1 | P61221 | 0.46 | 0.337242168 | 0.03 | 16.68 | 16.71 |
| 7 | ABCF1 | Q8NE71 | 0.58 | 0.236572006 | -0.06 | 14.71 | 14.66 |
| 8 | ABCF2 | Q9UG63 | 0.026 | 1.585026652 | 0.15 | 15.95 | 16.09 |
| 9 | ABHD10 | Q9NUJ1 | 0.22 | 0.657577319 | 0.23 | 14.21 | 14.44 |
| 10 | ABI1 | Q8IZP0 | 0.24 | 0.619788758 | -0.11 | 14.27 | 14.16 |
| 11 | ABI3 | Q9P2A4 | NA | NA | 0.08 | 15.43 | 15.51 |
| 12 | ABR | Q12979 | 0.51 | 0.292429824 | 0.05 | 14.88 | 14.93 |
| 13 | ABRACL | Q9P1F3 | NA | NA | 0.74 | 14.07 | 14.8 |
| 14 | ACAA1 | P09110 | 0.56 | 0.251811973 | -0.08 | 15.13 | 15.05 |
| 15 | ACAA2 | P42765 | 0.49 | 0.30980392 | -0.04 | 15.79 | 15.75 |
| 16 | ACACA | Q13085 | 0.019 | 1.721246399 | 0.64 | 13.74 | 14.39 |
| 17 | ACAD9 | Q9H845 | 0.84 | 0.075720714 | 0.02 | 14.22 | 14.24 |
| 18 | ACADM | P11310 | 0.0041 | 2.387216143 | -0.3 | 16.74 | 16.44 |
| 19 | ACADS | P16219 | NA | NA | 0.04 | 14.12 | 14.15 |
| 20 | ACADVL | P49748 | 0.87 | 0.060480747 | 0 | 16.81 | 16.81 |
| 21 | ACAT1 | P24752 | 0.42 | 0.37675071 | 0.04 | 16.6 | 16.64 |
| 22 | ACAT2 | Q9BWD1 | 0.014 | 1.853871964 | -0.12 | 15.21 | 15.08 |
| 23 | ACBD3 | Q9H3P7 | 1 | 0 | 0.02 | 13.72 | 13.74 |
| 24 | ACIN1 | Q9UKV3 | 0.00041 | 3.387216143 | -0.2 | 16.57 | 16.37 |
| 25 | ACLY | P53396 | 0.79 | 0.102372909 | 0 | 16.36 | 16.35 |
| 26 | ACO1 | P21399 | 0.16 | 0.795880017 | -0.1 | 15.37 | 15.27 |
| 27 | ACO2 | Q99798 | 0.54 | 0.26760624 | 0.02 | 16.06 | 16.08 |
| 28 | ACOT13 | Q9NPJ3 | 0.0048 | 2.318758763 | 0.29 | 17.12 | 17.41 |
| 29 | ACOT2 | P49753 (+1) | 0.81 | 0.091514981 | 0.03 | 15.92 | 15.95 |
| 30 | ACOT7 | O00154 | 0.056 | 1.251811973 | 0.17 | 16.15 | 16.31 |
| 31 | ACOT9 | Q9Y305 | 0.02 | 1.698970004 | 0.24 | 15.77 | 16.01 |
| 32 | ACP1 | P24666 | 0.44 | 0.356547324 | -0.09 | 15.52 | 15.44 |
| 33 | ACSL3 | O95573 | 0.037 | 1.431798276 | 0.16 | 15.28 | 15.44 |
| 34 | ACSL4 | O60488 | 0.2 | 0.698970004 | 0.17 | 15 | 15.17 |
| 35 | ACTB | P60709 | 0.83 | 0.080921908 | -0.03 | 15.94 | 15.91 |
| 36 | ACTC1 | P68032 | 0.22 | 0.657577319 | -0.16 | 18.12 | 17.96 |
| 37 | ACTG1 | P63261 | 0.55 | 0.259637311 | 0.12 | 14.87 | 14.99 |
| 38 | ACTN1 | P12814 | 0.0001 | 4 | 0.15 | 16.29 | 16.45 |
| 39 | ACTN4 | O43707 | 0.76 | 0.119186408 | 0 | 16.62 | 16.61 |
| 40 | ACTR1A | P61163 | 0.13 | 0.886056648 | -0.11 | 16.71 | 16.6 |
| 41 | ACTR2 | P61160 | 0.33 | 0.48148606 | 0.03 | 17.5 | 17.53 |
| 42 | ACTR3 | P61158 | 0.0098 | 2.008773924 | 0.11 | 16.78 | 16.89 |
| 43 | ADAM17 | P78536 | 0.26 | 0.585026652 | -0.08 | 16.89 | 16.81 |
| 44 | ADAM9 | Q13443 | 0.29 | 0.537602002 | 0.06 | 15.47 | 15.53 |
| 45 | ADAR | P55265 | 0.1 | 1 | 0.15 | 16.8 | 16.95 |
| 46 | ADD1 | P35611 | 0.28 | 0.552841969 | 0.15 | 16.41 | 16.56 |
| 47 | ADGRL4 | Q9HBW9 | 0.48 | 0.318758763 | 0.05 | 17.29 | 17.33 |

| | | | | | | | |
|---|---|---|---|---|---|---|---|
| 48 | ADH5 | P11766 | 0.11 | 0.958607315 | -0.13 | 18.01 | 17.88 |
| 49 | ADK | P55263 | 0.39 | 0.408935393 | -0.07 | 15.14 | 15.07 |
| 50 | ADPGK | Q9BRR6 | 0.005 | 2.301029996 | 0.27 | 16.01 | 16.28 |
| 51 | ADRM1 | Q16186 | 0.089 | 1.050609993 | -0.17 | 15.24 | 15.07 |
| 52 | ADSL | P30566 | 0.72 | 0.142667504 | -0.03 | 16.44 | 16.41 |
| 53 | ADSS | P30520 | 0.21 | 0.677780705 | 0.04 | 16.1 | 16.14 |
| 54 | AFDN | P55196 | 0.0016 | 2.795880017 | 0.19 | 15.6 | 15.79 |
| 55 | AFG3L2 | Q9Y4W6 | 0.076 | 1.119186408 | 0.54 | 14.16 | 14.69 |
| 56 | AGFG1 | P52594 | 0.14 | 0.853871964 | 0.11 | 17.73 | 17.84 |
| 57 | AGK | Q53H12 | 0.16 | 0.795880017 | 0.12 | 15.94 | 16.06 |
| 58 | AGPS | O00116 | 0.8 | 0.096910013 | 0.03 | 15.11 | 15.14 |
| 59 | AHCY | P23526 | 0.0011 | 2.958607315 | 0.14 | 17.56 | 17.7 |
| 60 | AHNAK | Q09666 | 0.0001 | 4 | -0.18 | 17.45 | 17.27 |
| 61 | AHNAK2 | Q8IVF2 | 0.64 | 0.193820026 | -0.03 | 17.69 | 17.67 |
| 62 | AHSA1 | O95433 | 0.31 | 0.508638306 | -0.1 | 16.63 | 16.53 |
| 63 | AIFM1 | O95831 | 0.7 | 0.15490196 | 0.01 | 16.15 | 16.17 |
| 64 | AIMP1 | Q12904 | 0.41 | 0.387216143 | -0.05 | 17.39 | 17.34 |
| 65 | AIMP2 | Q13155 | 0.17 | 0.769551079 | 0.15 | 16.21 | 16.36 |
| 66 | AK1 | P00568 | 0.41 | 0.387216143 | 0.02 | 16.36 | 16.39 |
| 67 | AK2 | P54819 | 0.072 | 1.142667504 | -0.15 | 16.93 | 16.78 |
| 68 | AK3 | Q9UIJ7 | 0.027 | 1.568636236 | 0.13 | 16.49 | 16.62 |
| 69 | AKAP12 | Q02952 | 0.0002 | 3.698970004 | 0.11 | 16.39 | 16.51 |
| 70 | AKAP2 | Q9Y2D5 | 0.65 | 0.187086643 | -0.02 | 15.57 | 15.56 |
| 71 | AKAP8 | O43823 | 0.044 | 1.356547324 | 0.53 | 15.39 | 15.92 |
| 72 | AKR1A1 | P14550 | 0.018 | 1.744727495 | -0.15 | 15.06 | 14.91 |
| 73 | AKR1B1 | P15121 | 0.92 | 0.036212173 | 0 | 18.38 | 18.38 |
| 74 | AKR1C3 | P42330 | 0.27 | 0.568636236 | 0.1 | 17.28 | 17.39 |
| 75 | AKR7A2 | O43488 | 0.74 | 0.13076828 | 0.07 | 14.98 | 15.06 |
| 76 | ALB | P02768 | 0.14 | 0.853871964 | 0.32 | 17.87 | 18.19 |
| 77 | ALDH16A1 | Q8IZ83 | NA | NA | -0.09 | 14.94 | 14.85 |
| 78 | ALDH18A1 | P54886 | 0.11 | 0.958607315 | 0.22 | 15.57 | 15.79 |
| 79 | ALDH1A1 | P00352 | 0.0001 | 4 | -0.26 | 17.07 | 16.81 |
| 80 | ALDH1B1 | P30837 | 0.11 | 0.958607315 | 0.28 | 14.3 | 14.59 |
| 81 | ALDH2 | P05091 | 0.0001 | 4 | 0.23 | 16.34 | 16.57 |
| 82 | ALDH3A2 | P51648 | 0.029 | 1.537602002 | -0.27 | 16.62 | 16.35 |
| 83 | ALDH4A1 | P30038 | 0.064 | 1.193820026 | 0.12 | 17 | 17.11 |
| 84 | ALDH6A1 | Q02252 | 0.0035 | 2.455931956 | 0.16 | 16.22 | 16.39 |
| 85 | ALDH7A1 | P49419 | 0.00026 | 3.585026652 | 0.18 | 16.26 | 16.44 |
| 86 | ALDH9A1 | P49189 | 0.0044 | 2.356547324 | 0.23 | 16.58 | 16.81 |
| 87 | ALDOA | P04075 | 0.64 | 0.193820026 | -0.02 | 17.51 | 17.48 |
| 88 | ALYREF | Q86V81 | 0.25 | 0.602059991 | -0.07 | 16.73 | 16.66 |
| 89 | ANKRD17 | O75179 | 0.015 | 1.823908741 | 0.11 | 16.51 | 16.62 |
| 90 | ANO6 | Q4KMQ2 | 0.74 | 0.13076828 | 0.09 | 15.73 | 15.82 |
| 91 | ANP32A | P39687 | 0.063 | 1.200659451 | -0.28 | 18.07 | 17.79 |
| 92 | ANP32B | Q92688 | 0.34 | 0.468521083 | -0.1 | 17.56 | 17.46 |
| 93 | ANP32E | Q9BTT0 | 0.13 | 0.886056648 | 0.07 | 14.44 | 14.51 |
| 94 | ANPEP | P15144 | 0.00014 | 3.853871964 | 0.12 | 16.02 | 16.14 |
| 95 | ANXA1 | P04083 | 0.0001 | 4 | -0.23 | 17.62 | 17.39 |
| 96 | ANXA11 | P50995 | 0.00019 | 3.721246399 | -0.17 | 17.04 | 16.88 |
| 97 | ANXA2 | P07355 | 0.0001 | 4 | -0.17 | 17.66 | 17.49 |

| | | | | | | | |
|---|---|---|---|---|---|---|---|
| 98 | ANXA3 | P12429 | 0.5 | 0.301029996 | -0.05 | 16.33 | 16.28 |
| 99 | ANXA4 | P09525 | 0.025 | 1.602059991 | -0.13 | 16.16 | 16.02 |
| 100 | ANXA5 | P08758 | 0.0078 | 2.107905397 | -0.14 | 16.18 | 16.04 |
| 101 | ANXA6 | P08133 | 0.16 | 0.795880017 | 0.03 | 16.74 | 16.77 |
| 102 | ANXA7 | P20073 | 0.014 | 1.853871964 | -0.19 | 17.59 | 17.4 |
| 103 | AP1B1 | Q10567 | 0.022 | 1.657577319 | 0.2 | 15.27 | 15.47 |
| 104 | AP1G1 | O43747 | 0.21 | 0.677780705 | 0.09 | 15.26 | 15.35 |
| 105 | AP2A1 | O95782 | 0.041 | 1.387216143 | 0.13 | 16.44 | 16.56 |
| 106 | AP2A2 | O94973 | 0.021 | 1.677780705 | 0.14 | 15.6 | 15.74 |
| 107 | AP2B1 | P63010 | 0.12 | 0.920818754 | 0.07 | 15.03 | 15.1 |
| 108 | AP2M1 | Q96CW1 | 0.011 | 1.958607315 | 0.24 | 17.67 | 17.91 |
| 109 | AP2S1 | P53680 | 0.52 | 0.283996656 | 0.11 | 17.52 | 17.63 |
| 110 | AP3B1 | O00203 | 0.064 | 1.193820026 | 0.14 | 16.36 | 16.5 |
| 111 | AP3D1 | O14617 | 0.046 | 1.337242168 | 0.11 | 16.61 | 16.72 |
| 112 | AP3M1 | Q9Y2T2 | 0.0062 | 2.207608311 | 0.36 | 15.83 | 16.19 |
| 113 | APEX1 | P27695 | 0.81 | 0.091514981 | 0.01 | 15.5 | 15.51 |
| 114 | API5 | Q9BZZ5 | 0.075 | 1.124938737 | 0.06 | 15.35 | 15.41 |
| 115 | APMAP | Q9HDC9 | 0.012 | 1.920818754 | 0.1 | 17 | 17.1 |
| 116 | APOL2 | Q9BQE5 | 0.14 | 0.853871964 | 0.16 | 15.7 | 15.85 |
| 117 | APOOL | Q6UXV4 | 0.056 | 1.251811973 | 0.25 | 14.08 | 14.33 |
| 118 | APP | P05067 | 0.34 | 0.468521083 | -0.14 | 17.09 | 16.95 |
| 119 | APRT | P07741 | 0.15 | 0.823908741 | 0.05 | 15.54 | 15.59 |
| 120 | ARAP3 | Q8WWN8 | 0.0088 | 2.055517328 | 0.2 | 14.83 | 15.03 |
| 121 | ARCN1 | P48444 | 0.016 | 1.795880017 | 0.1 | 16.53 | 16.63 |
| 122 | ARF3 | P61204 | 0.42 | 0.37675071 | 0.08 | 16.92 | 17 |
| 123 | ARF4 | P18085 | 0.13 | 0.886056648 | -0.13 | 16.35 | 16.22 |
| 124 | ARF5 | P84085 | 0.095 | 1.022276395 | 0.16 | 15.9 | 16.05 |
| 125 | ARF6 | P62330 | 0.00042 | 3.37675071 | 0.36 | 15.79 | 16.15 |
| 126 | ARFGEF1 | Q9Y6D6 | 0.003 | 2.522878745 | 0.19 | 15.93 | 16.13 |
| 127 | ARHGAP1 | Q07960 | 0.67 | 0.173925197 | -0.03 | 13.74 | 13.71 |
| 128 | ARHGAP17 | Q68EM7 | 0.27 | 0.568636236 | 0.08 | 15.4 | 15.48 |
| 129 | ARHGAP18 | Q8N392 | 0.73 | 0.13667714 | 0.03 | 15.49 | 15.52 |
| 130 | ARHGAP31 | Q2M1Z3 | 0.44 | 0.356547324 | 0.29 | 14.74 | 15.03 |
| 131 | ARHGDIA | P52565 | 0.0075 | 2.124938737 | -0.13 | 17.85 | 17.72 |
| 132 | ARHGDIB | P52566 | 0.66 | 0.180456064 | -0.02 | 16.92 | 16.91 |
| 133 | ARHGEF7 | Q14155 | 0.6 | 0.22184875 | 0.07 | 15.53 | 15.6 |
| 134 | ARL1 | P40616 | 0.017 | 1.769551079 | -0.26 | 16.23 | 15.97 |
| 135 | ARL2 | P36404 | 0.56 | 0.251811973 | 0.11 | 15.99 | 16.1 |
| 136 | ARL3 | P36405 | 0.69 | 0.161150909 | 0.08 | 14.5 | 14.57 |
| 137 | ARL6IP5 | O75915 | 0.0051 | 2.292429824 | 0.13 | 16.14 | 16.27 |
| 138 | ARL8A | Q96BM9 | 0.35 | 0.455931956 | 0.09 | 14.97 | 15.05 |
| 139 | ARMC6 | Q6NXE6 | 0.14 | 0.853871964 | 0.24 | 15.94 | 16.18 |
| 140 | ARMT1 | Q9H993 | 0.22 | 0.657577319 | 0.22 | 16.36 | 16.58 |
| 141 | ARPC1A | Q92747 | 0.43 | 0.366531544 | 0.05 | 18.96 | 19.01 |
| 142 | ARPC1B | O15143 | 0.11 | 0.958607315 | 0.14 | 18.37 | 18.51 |
| 143 | ARPC2 | O15144 | 0.12 | 0.920818754 | 0.05 | 17.04 | 17.09 |
| 144 | ARPC3 | O15145 | 0.034 | 1.468521083 | 0.31 | 15.63 | 15.94 |
| 145 | ARPC4 | P59998 | 0.28 | 0.552841969 | 0.09 | 17.74 | 17.83 |
| 146 | ARPC5 | O15511 | 0.15 | 0.823908741 | 0.14 | 16.72 | 16.85 |
| 147 | ARPC5L | Q9BPX5 | 0.33 | 0.48148606 | 0.09 | 16.78 | 16.86 |

| | | | | | | | | |
|---|---|---|---|---|---|---|---|---|
| 148 | ARRB1 | P49407 | 0.21 | 0.677780705 | -0.12 | 15.67 | 15.55 | |
| 149 | ARSB | P15848 | 0.06 | 1.22184875 | 0.17 | 16.11 | 16.28 | |
| 150 | ASAH1 | Q13510 | 0.2 | 0.698970004 | 0.2 | 14.9 | 15.1 | |
| 151 | ASAP2 | O43150 | 0.33 | 0.48148606 | 0.16 | 17.06 | 17.23 | |
| 152 | ASCC3 | Q8N3C0 | 0.33 | 0.48148606 | 0.08 | 16.58 | 16.66 | |
| 153 | ASMTL | O95671 | 0.058 | 1.236572006 | 0.23 | 17.58 | 17.81 | |
| 154 | ASNA1 | O43681 | 0.2 | 0.698970004 | 0.15 | 15.8 | 15.95 | |
| 155 | ASPH | Q12797 | 0.089 | 1.050609993 | -0.1 | 15.6 | 15.5 | |
| 156 | ASPM | Q8IZT6 | 0.003 | 2.522878745 | 0.39 | 14.76 | 15.15 | |
| 157 | ASRGL1 | Q7L266 | 0.54 | 0.26760624 | -0.12 | 15.77 | 15.66 | |
| 158 | ATAD3A | Q9NVI7 | 0.58 | 0.236572006 | -0.09 | 13.82 | 13.73 | |
| 159 | ATAD3B | Q5T9A4 | 0.22 | 0.657577319 | 0.1 | 16.98 | 17.08 | |
| 160 | ATIC | P31939 | 0.97 | 0.013228266 | 0 | 15.49 | 15.49 | |
| 161 | ATL3 | Q6DD88 | 0.18 | 0.744727495 | 0.06 | 16.61 | 16.67 | |
| 162 | ATOX1 | O00244 | 0.76 | 0.119186408 | -0.06 | 16.57 | 16.51 | |
| 163 | ATP13A1 | Q9HD20 | 0.023 | 1.638272164 | 0.31 | 16.42 | 16.73 | |
| 164 | ATP1A1 | P05023 | 0.48 | 0.318758763 | 0.02 | 16.47 | 16.49 | |
| 165 | ATP1B3 | P54709 | 0.32 | 0.494850022 | 0.15 | 15.75 | 15.89 | |
| 166 | ATP2A2 | P16615 | 0.028 | 1.552841969 | 0.13 | 16.07 | 16.2 | |
| 167 | ATP2B4 | P23634 | 0.094 | 1.026872146 | 0.13 | 15.49 | 15.62 | |
| 168 | ATP5F1A | P25705 | 0.0026 | 2.585026652 | 0.11 | 16.79 | 16.9 | |
| 169 | ATP5F1B | P06576 | 0.0001 | 4 | 0.14 | 16.22 | 16.36 | |
| 170 | ATP5F1C | P36542 | 0.28 | 0.552841969 | 0.08 | 15.49 | 15.57 | |
| 171 | ATP5F1D | P30049 | 0.42 | 0.37675071 | 0.09 | 17.27 | 17.36 | |
| 172 | ATP5MF | P56134 | 0.31 | 0.508638306 | 0.09 | 16.79 | 16.89 | |
| 173 | ATP5MG | O75964 | 0.0055 | 2.259637311 | 0.24 | 16.99 | 17.23 | |
| 174 | ATP5MPL | P56378 | 0.98 | 0.008773924 | 0.01 | 19.65 | 19.66 | |
| 175 | ATP5PB | P24539 | 0.11 | 0.958607315 | 0.25 | 17.83 | 18.08 | |
| 176 | ATP5PD | O75947 | 0.24 | 0.619788758 | -0.07 | 16.18 | 16.1 | |
| 177 | ATP5PO | P48047 | 0.0001 | 4 | 0.38 | 15.19 | 15.57 | |
| 178 | ATP6V1A | P38606 | 0.017 | 1.769551079 | -0.21 | 16.06 | 15.85 | |
| 179 | ATP6V1B2 | | P21281 | 0.13 | 0.886056648 | -0.09 | 16 | 15.91 |
| 180 | ATP6V1E1 | | P36543 | 0.021 | 1.677780705 | -0.28 | 15.63 | 15.35 |
| 181 | ATP6V1H | Q9UI12 | 0.93 | 0.031517051 | 0.02 | 15.2 | 15.21 | |
| 182 | ATXN10 | Q9UBB4 | 0.031 | 1.508638306 | 0.12 | 15.81 | 15.93 | |
| 183 | ATXN2L | Q8WWM7 | 0.084 | 1.075720714 | 0.18 | 15.73 | 15.91 | |
| 184 | B2M | P61769 | 0.00063 | 3.200659451 | -0.43 | 18.49 | 18.06 | |
| 185 | BAG1 | Q99933 | 0.85 | 0.070581074 | 0.08 | 14.59 | 14.67 | |
| 186 | BAG2 | O95816 | 0.79 | 0.102372909 | 0.01 | 16.84 | 16.86 | |
| 187 | BAG3 | O95817 | 0.042 | 1.37675071 | -0.5 | 16.63 | 16.13 | |
| 188 | BAG6 | P46379 | 0.25 | 0.602059991 | 0.12 | 15.03 | 15.15 | |
| 189 | BASP1 | P80723 | 0.0009 | 3.045757491 | 0.24 | 17.01 | 17.25 | |
| 190 | BAX | Q07812 | 0.091 | 1.040958608 | -0.33 | 18.15 | 17.81 | |
| 191 | BCAP31 | P51572 | 0.33 | 0.48148606 | 0.04 | 18.85 | 18.89 | |
| 192 | BCAR1 | P56945 | 0.32 | 0.494850022 | 0.11 | 14.69 | 14.8 | |
| 193 | BCAT1 | P54687 | 0.099 | 1.004364805 | 0.1 | 16.43 | 16.53 | |
| 194 | BCHE | P06276 | 0.86 | 0.065501549 | 0.06 | 14.72 | 14.77 | |
| 195 | BCKDHA | P12694 | 0.22 | 0.657577319 | 0.18 | 15.35 | 15.53 | |
| 196 | BCL2L13 | Q9BXK5 | 0.19 | 0.721246399 | -0.18 | 14.56 | 14.38 | |
| 197 | BCLAF1 | Q9NYF8 | 0.4 | 0.397940009 | -0.09 | 16.45 | 16.36 | |

| | | | | | | | |
|---|---|---|---|---|---|---|---|
| 198 | BCS1L | Q9Y276 | 0.23 | 0.638272164 | 0.12 | 15.55 | 15.67 |
| 199 | BLMH | Q13867 | 0.94 | 0.026872146 | 0.05 | 17.33 | 17.38 |
| 200 | BLVRA | P53004 | 0.015 | 1.823908741 | -0.23 | 14.86 | 14.63 |
| 201 | BMP2K | Q9NSY1 | 0.32 | 0.494850022 | 0.16 | 15.26 | 15.41 |
| 202 | BMS1 | Q14692 | 0.11 | 0.958607315 | 0.37 | 14.61 | 14.99 |
| 203 | BMX | P51813 | 0.31 | 0.508638306 | -0.2 | 14.9 | 14.7 |
| 204 | BRIX1 | Q8TDN6 | 0.53 | 0.27572413 | 0.09 | 15.31 | 15.4 |
| 205 | BSG | P35613 | 0.66 | 0.180456064 | -0.04 | 17.78 | 17.75 |
| 206 | BTF3 | P20290 | 0.21 | 0.677780705 | -0.1 | 15.56 | 15.45 |
| 207 | BTF3L4 | Q96K17 | 0.18 | 0.744727495 | -0.29 | 15.12 | 14.82 |
| 208 | BUB3 | O43684 | 0.43 | 0.366531544 | 0.07 | 17.61 | 17.68 |
| 209 | BZW1 | Q7L1Q6 | 0.93 | 0.031517051 | -0.01 | 16.96 | 16.95 |
| 210 | BZW2 | Q9Y6E2 | 0.7 | 0.15490196 | 0.12 | 14.57 | 14.69 |
| 211 | C12orf10 | Q9HB07 | 0.46 | 0.337242168 | 0.26 | 16.19 | 16.45 |
| 212 | C1QBP | Q07021 | 0.13 | 0.886056648 | -0.07 | 15.99 | 15.92 |
| 213 | C20orf27 | Q9GZN8 | 0.35 | 0.455931956 | 0.23 | 15.15 | 15.38 |
| 214 | CACNA2D1 | P54289 | 0.0069 | 2.161150909 | -0.35 | 14.26 | 13.91 |
| 215 | CACYBP | Q9HB71 | 0.43 | 0.366531544 | -0.09 | 17.12 | 17.03 |
| 216 | CAD | P27708 | 0.0001 | 4 | 0.2 | 15.49 | 15.69 |
| 217 | CALD1 | Q05682 | 0.021 | 1.677780705 | 0.08 | 17.19 | 17.26 |
| 218 | CALM1 | P0DP23 (+2) | 0.0055 | 2.259637311 | -0.12 | 17.56 | 17.44 |
| 219 | CALR | P27797 | 0.0001 | 4 | -0.6 | 16.91 | 16.31 |
| 220 | CALU | O43852 | 0.002 | 2.698970004 | -0.23 | 16.7 | 16.47 |
| 221 | CAMK2D | Q13557 | 0.29 | 0.537602002 | 0.1 | 15.61 | 15.71 |
| 222 | CAND1 | Q86VP6 | 0.0001 | 4 | 0.26 | 16.07 | 16.32 |
| 223 | CANX | P27824 | 0.16 | 0.795880017 | -0.07 | 17.44 | 17.37 |
| 224 | CAP1 | Q01518 | 0.016 | 1.795880017 | 0.1 | 16.96 | 17.05 |
| 225 | CAPG | P40121 | 0.0009 | 3.045757491 | 0.34 | 17.89 | 18.23 |
| 226 | CAPN1 | P07384 | 0.035 | 1.455931956 | 0.24 | 15.01 | 15.25 |
| 227 | CAPN2 | P17655 | 0.015 | 1.823908741 | 0.1 | 16.07 | 16.17 |
| 228 | CAPNS1 | P04632 | 0.0011 | 2.958607315 | 0.15 | 15.04 | 15.19 |
| 229 | CAPRIN1 | Q14444 | 0.12 | 0.920818754 | -0.19 | 17.33 | 17.14 |
| 230 | CAPZA1 | P52907 | 0.011 | 1.958607315 | 0.1 | 16.34 | 16.44 |
| 231 | CAPZA2 | P47755 | 0.4 | 0.397940009 | -0.05 | 16.09 | 16.05 |
| 232 | CAPZB | P47756 | 0.26 | 0.585026652 | 0.07 | 17.08 | 17.15 |
| 233 | CARHSP1 | Q9Y2V2 | 0.54 | 0.26760624 | 0.06 | 15.53 | 15.59 |
| 234 | CARM1 | Q86X55 | 0.03 | 1.522878745 | 0.36 | 17.2 | 17.57 |
| 235 | CARS | P49589 | 0.93 | 0.031517051 | 0 | 15.69 | 15.69 |
| 236 | CASP3 | P42574 | 0.48 | 0.318758763 | 0.09 | 15.35 | 15.43 |
| 237 | CASP7 | P55210 | 0.024 | 1.619788758 | 0.27 | 16.35 | 16.62 |
| 238 | CASP8 | Q14790 | 0.99 | 0.004364805 | 0.01 | 15.48 | 15.49 |
| 239 | CAST | P20810 | 0.017 | 1.769551079 | -0.1 | 15.48 | 15.38 |
| 240 | CAT | P04040 | 0.32 | 0.494850022 | -0.05 | 16.21 | 16.16 |
| 241 | CAV1 | Q03135 | 0.0001 | 4 | -0.23 | 18.69 | 18.46 |
| 242 | CAVIN1 | Q6NZI2 | 0.0001 | 4 | -0.39 | 17.02 | 16.63 |
| 243 | CAVIN2 | O95810 | 0.0001 | 4 | -0.36 | 16.5 | 16.13 |
| 244 | CBFB | Q13951 | 0.73 | 0.13667714 | -0.04 | 15.35 | 15.31 |
| 245 | CBR1 | P16152 | 0.89 | 0.050609993 | 0 | 16.98 | 16.97 |
| 246 | CBX1 | P83916 | 0.023 | 1.638272164 | -0.2 | 14.48 | 14.28 |
| 247 | CBX3 | Q13185 | 0.0001 | 4 | -0.18 | 15.76 | 15.59 |

| | | | | | | | |
|---|---|---|---|---|---|---|---|
| 248 | CCAR1 | Q8IX12 | 0.025 | 1.602059991 | 0.18 | 17.22 | 17.4 |
| 249 | CCAR2 | Q8N163 | 0.89 | 0.050609993 | 0.01 | 15.72 | 15.73 |
| 250 | CCDC47 | Q96A33 | 0.15 | 0.823908741 | 0.13 | 16.19 | 16.32 |
| 251 | CCDC50 | Q8IVM0 | 0.79 | 0.102372909 | 0.05 | 15.24 | 15.29 |
| 252 | CCDC6 | Q16204 | 0.55 | 0.259637311 | 0.07 | 13.17 | 13.24 |
| 253 | CCDC7 | Q96M83 | NA | NA | 0.17 | 13.48 | 13.65 |
| 254 | CCS | O14618 | 0.88 | 0.055517328 | -0.02 | 15.42 | 15.4 |
| 255 | CCT2 | P78371 | 0.024 | 1.619788758 | 0.09 | 16.25 | 16.34 |
| 256 | CCT3 | P49368 | 0.0082 | 2.086186148 | 0.09 | 16.73 | 16.82 |
| 257 | CCT4 | P50991 | 0.0001 | 4 | 0.2 | 16.33 | 16.52 |
| 258 | CCT5 | P48643 | 0.00029 | 3.537602002 | 0.1 | 16.62 | 16.73 |
| 259 | CCT6A | P40227 | 0.28 | 0.552841969 | 0.04 | 16.48 | 16.52 |
| 260 | CCT7 | Q99832 | 0.0001 | 4 | 0.19 | 16.22 | 16.41 |
| 261 | CCT8 | P50990 | 0.0004 | 3.397940009 | 0.1 | 17.51 | 17.61 |
| 262 | CD2AP | Q9Y5K6 | 0.51 | 0.292429824 | -0.03 | 17.38 | 17.35 |
| 263 | CD44 | P16070 | 0.0001 | 4 | 0.45 | 16.65 | 17.1 |
| 264 | CD47 | Q08722 | 0.19 | 0.721246399 | -0.11 | 17.58 | 17.47 |
| 265 | CD55 | P08174 | 0.019 | 1.721246399 | 0.49 | 17.27 | 17.76 |
| 266 | CD59 | P13987 | 0.0074 | 2.13076828 | -0.22 | 19.46 | 19.24 |
| 267 | CD9 | P21926 | 0.037 | 1.431798276 | -0.27 | 18.07 | 17.8 |
| 268 | CD93 | Q9NPY3 | 0.53 | 0.27572413 | 0.05 | 16.16 | 16.2 |
| 269 | CDC37 | Q16543 | 0.23 | 0.638272164 | 0.05 | 16.38 | 16.42 |
| 270 | CDC42 | P60953 | 0.087 | 1.060480747 | 0.1 | 16.54 | 16.64 |
| 271 | CDC42BPB | Q9Y5S2 | 0.76 | 0.119186408 | 0.03 | 17.28 | 17.31 |
| 272 | CDC5L | Q99459 | 0.38 | 0.420216403 | 0.1 | 16.23 | 16.33 |
| 273 | CDK17 | Q00537 | 0.92 | 0.036212173 | 0.01 | 16.73 | 16.74 |
| 274 | CDS2 | O95674 | 0.46 | 0.337242168 | 0.11 | 15.52 | 15.64 |
| 275 | CDV3 | Q9UKY7 | 0.096 | 1.017728767 | -0.08 | 15.92 | 15.84 |
| 276 | CEBPZ | Q03701 | 0.4 | 0.397940009 | -0.39 | 15.65 | 15.26 |
| 277 | CELF1 | Q92879 | 0.39 | 0.408935393 | 0.07 | 18.29 | 18.36 |
| 278 | CEP170 | Q5SW79 | 0.41 | 0.387216143 | 0.14 | 16.78 | 16.92 |
| 279 | CFL1 | P23528 | 0.032 | 1.494850022 | -0.14 | 17.32 | 17.18 |
| 280 | CFL2 | Q9Y281 | 0.27 | 0.568636236 | -0.07 | 17.15 | 17.08 |
| 281 | CHAMP1 | Q96JM3 | 0.58 | 0.236572006 | 0.06 | 14.72 | 14.77 |
| 282 | CHCHD3 | Q9NX63 | 0.018 | 1.744727495 | -0.11 | 17.38 | 17.26 |
| 283 | CHD4 | Q14839 | 0.24 | 0.619788758 | 0.25 | 13.81 | 14.06 |
| 284 | CHD6 | Q8TD26 | 0.72 | 0.142667504 | 0.06 | 14.12 | 14.17 |
| 285 | CHID1 | Q9BWS9 | 0.56 | 0.251811973 | 0.04 | 15.14 | 15.18 |
| 286 | CHTOP | Q9Y3Y2 | 0.047 | 1.327902142 | -0.2 | 17.89 | 17.7 |
| 287 | CIAPIN1 | Q6FI81 | 0.5 | 0.301029996 | -0.16 | 17.77 | 17.61 |
| 288 | CIRBP | Q14011 | 0.012 | 1.920818754 | -0.36 | 16.5 | 16.14 |
| 289 | CISD1 | Q9NZ45 | 0.026 | 1.585026652 | -0.13 | 19.3 | 19.17 |
| 290 | CKAP4 | Q07065 | 0.69 | 0.161150909 | -0.02 | 17.65 | 17.63 |
| 291 | CKAP5 | Q14008 | 0.088 | 1.055517328 | 0.07 | 16.19 | 16.26 |
| 292 | CLASP1 | Q7Z460 | 0.14 | 0.853871964 | 0.28 | 15.21 | 15.48 |
| 293 | CLEC14A | Q86T13 | 0.046 | 1.337242168 | -0.21 | 15.7 | 15.49 |
| 294 | CLIC1 | O00299 | 0.096 | 1.017728767 | 0.11 | 16.55 | 16.65 |
| 295 | CLIC4 | Q9Y696 | 0.0001 | 4 | 0.45 | 15.69 | 16.14 |
| 296 | CLINT1 | Q14677 | 0.89 | 0.050609993 | -0.05 | 18.25 | 18.19 |
| 297 | CLIP1 | P30622 | 0.94 | 0.026872146 | -0.01 | 16.81 | 16.8 |

| | | | | | | | |
|---|---|---|---|---|---|---|---|
| 298 | CLPTM1 | O96005 | 0.31 | 0.508638306 | 0.14 | 16.57 | 16.71 |
| 299 | CLPTM1L | Q96KA5 | 0.0052 | 2.283996656 | 0.17 | 15.81 | 15.98 |
| 300 | CLPX | O76031 | 0.17 | 0.769551079 | 0.1 | 16.61 | 16.71 |
| 301 | CLTA | P09496 | 0.0035 | 2.455931956 | -0.24 | 19.25 | 19.01 |
| 302 | CLTB | P09497 | 0.02 | 1.698970004 | 0.29 | 18.2 | 18.49 |
| 303 | CLTC | Q00610 | 0.0085 | 2.070581074 | 0.05 | 16.21 | 16.27 |
| 304 | CMPK1 | P30085 | 0.0053 | 2.27572413 | 0.22 | 15.2 | 15.42 |
| 305 | CNDP2 | Q96KP4 | 0.25 | 0.602059991 | -0.04 | 15.78 | 15.74 |
| 306 | CNN2 | Q99439 | 0.0079 | 2.102372909 | 0.15 | 17.51 | 17.66 |
| 307 | CNN3 | Q15417 | 0.31 | 0.508638306 | 0.03 | 16.88 | 16.91 |
| 308 | CNOT1 | A5YKK6 | 0.14 | 0.853871964 | 0.34 | 14.29 | 14.64 |
| 309 | CNP | P09543 | 0.51 | 0.292429824 | -0.04 | 16.18 | 16.13 |
| 310 | CNPY2 | Q9Y2B0 | 0.62 | 0.207608311 | -0.04 | 14.95 | 14.9 |
| 311 | CNPY3 | Q9BT09 | 0.15 | 0.823908741 | -0.14 | 16.19 | 16.05 |
| 312 | CNRIP1 | Q96F85 | 0.033 | 1.48148606 | 0.23 | 15.01 | 15.25 |
| 313 | COL18A1 | P39060 | 0.045 | 1.346787486 | 0.34 | 15.46 | 15.8 |
| 314 | COL4A1 | P02462 | 0.086 | 1.065501549 | 0.19 | 15.72 | 15.91 |
| 315 | COL4A2 | P08572 | 0.025 | 1.602059991 | 0.57 | 16.86 | 17.43 |
| 316 | COLEC12 | Q5KU26 | 0.14 | 0.853871964 | 0.27 | 17.01 | 17.28 |
| 317 | COLGALT1 | Q8NBJ5 | 0.43 | 0.366531544 | -0.05 | 17.13 | 17.08 |
| 318 | COMT | P21964 | 0.054 | 1.26760624 | 0.1 | 14.99 | 15.09 |
| 319 | COPA | P53621 | 0.089 | 1.050609993 | 0.05 | 15.66 | 15.72 |
| 320 | COPB1 | P53618 | 0.24 | 0.619788758 | -0.04 | 16.32 | 16.28 |
| 321 | COPB2 | P35606 | 0.59 | 0.229147988 | 0.02 | 16.56 | 16.59 |
| 322 | COPE | O14579 | 0.43 | 0.366531544 | -0.06 | 14.65 | 14.59 |
| 323 | COPG1 | Q9Y678 | 0.71 | 0.148741651 | -0.02 | 15.85 | 15.83 |
| 324 | COPG2 | Q9UBF2 | 0.92 | 0.036212173 | 0 | 16.52 | 16.52 |
| 325 | COPS2 | P61201 | 0.8 | 0.096910013 | -0.02 | 15.23 | 15.21 |
| 326 | COPS3 | Q9UNS2 | 0.13 | 0.886056648 | 0.09 | 17.14 | 17.24 |
| 327 | COPS4 | Q9BT78 | 0.19 | 0.721246399 | 0.09 | 16.3 | 16.39 |
| 328 | COPS5 | Q92905 | 0.0088 | 2.055517328 | 0.41 | 15.12 | 15.54 |
| 329 | COPS6 | Q7L5N1 | 0.27 | 0.568636236 | 0.08 | 16.18 | 16.25 |
| 330 | COPS7B | Q9H9Q2 | 0.63 | 0.200659451 | -0.04 | 13.91 | 13.87 |
| 331 | COPS8 | Q99627 | 0.44 | 0.356547324 | -0.05 | 16.46 | 16.41 |
| 332 | COPZ1 | P61923 | 0.3 | 0.522878745 | -0.05 | 16.59 | 16.54 |
| 333 | CORO1B | Q9BR76 | 0.48 | 0.318758763 | 0.07 | 14.73 | 14.8 |
| 334 | CORO1C | Q9ULV4 | 0.00089 | 3.050609993 | 0.21 | 16.16 | 16.37 |
| 335 | CORO7 | P57737 | 0.39 | 0.408935393 | 0.09 | 14.64 | 14.73 |
| 336 | COTL1 | Q14019 | 0.035 | 1.455931956 | -0.36 | 17.4 | 17.03 |
| 337 | COX4I1 | P13073 | 0.32 | 0.494850022 | 0.04 | 17.71 | 17.75 |
| 338 | COX5A | P20674 | 0.52 | 0.283996656 | -0.04 | 17 | 16.97 |
| 339 | COX5B | P10606 | 0.75 | 0.124938737 | 0.01 | 17.89 | 17.9 |
| 340 | CPNE1 | Q99829 | 0.7 | 0.15490196 | 0.02 | 16.49 | 16.51 |
| 341 | CPNE3 | O75131 | 0.21 | 0.677780705 | -0.12 | 15.24 | 15.12 |
| 342 | CPOX | P36551 | 0.32 | 0.494850022 | 0.15 | 16.67 | 16.82 |
| 343 | CPPED1 | Q9BRF8 | 0.11 | 0.958607315 | -0.15 | 16.04 | 15.89 |
| 344 | CPSF6 | Q16630 | 0.32 | 0.494850022 | -0.11 | 15.99 | 15.88 |
| 345 | CPSF7 | Q8N684 | 0.53 | 0.27572413 | 0.05 | 15.51 | 15.56 |
| 346 | CPT1A | P50416 | 0.28 | 0.552841969 | 0.26 | 14.2 | 14.46 |
| 347 | CREBBP | Q92793 | 0.82 | 0.086186148 | -0.02 | 13.14 | 13.13 |

| | | | | | | | |
|---|---|---|---|---|---|---|---|
| 348 | CRELD1 | Q96HD1 | 0.072 | 1.142667504 | -0.19 | 17.04 | 16.85 |
| 349 | CRELD2 | Q6UXH1 | 0.93 | 0.031517051 | 0.01 | 15.05 | 15.06 |
| 350 | CRIP2 | P52943 | 0.00067 | 3.173925197 | -0.14 | 17.6 | 17.47 |
| 351 | CRK | P46108 | 0.26 | 0.585026652 | 0.08 | 15.3 | 15.38 |
| 352 | CRKL | P46109 | 0.057 | 1.244125144 | 0.24 | 16.45 | 16.69 |
| 353 | CROT | Q9UKG9 | 0.0021 | 2.677780705 | -0.14 | 14.1 | 13.96 |
| 354 | CRTAP | O75718 | 0.063 | 1.200659451 | 0.45 | 14.59 | 15.04 |
| 355 | CRYZ | Q08257 | 0.97 | 0.013228266 | -0.01 | 16.06 | 16.06 |
| 356 | CS | O75390 | 0.0003 | 3.522878745 | 0.23 | 15.31 | 15.54 |
| 357 | CSDE1 | O75534 | 0.45 | 0.346787486 | 0.19 | 16.64 | 16.84 |
| 358 | CSE1L | P55060 | 0.084 | 1.075720714 | 0.07 | 15.8 | 15.87 |
| 359 | CSNK2A1 | P68400 | 0.062 | 1.207608311 | -0.14 | 16.87 | 16.73 |
| 360 | CSNK2B | P67870 | 0.55 | 0.259637311 | 0.13 | 15.21 | 15.34 |
| 361 | CSRP1 | P21291 | 0.0058 | 2.236572006 | 0.27 | 17.87 | 18.14 |
| 362 | CSTB | P04080 | 0.013 | 1.886056648 | 0.33 | 17.24 | 17.57 |
| 363 | CSTF2T | Q9H0L4 | 0.048 | 1.318758763 | 0.18 | 16.15 | 16.33 |
| 364 | CTNNA1 | P35221 | 0.018 | 1.744727495 | 0.12 | 15.36 | 15.48 |
| 365 | CTNNA2 | P26232 | NA | NA | -0.05 | 15.75 | 15.69 |
| 366 | CTNNB1 | P35222 | 0.0047 | 2.327902142 | 0.17 | 15.46 | 15.63 |
| 367 | CTNNBL1 | Q8WYA6 | NA | NA | 0.37 | 15.65 | 16.02 |
| 368 | CTNND1 | O60716 | 0.64 | 0.193820026 | 0.02 | 15.63 | 15.65 |
| 369 | CTPS1 | P17812 | 0.39 | 0.408935393 | 0.07 | 16.22 | 16.29 |
| 370 | CTSA | P10619 | 0.76 | 0.119186408 | -0.03 | 13.65 | 13.62 |
| 371 | CTSB | P07858 | 0.018 | 1.744727495 | 0.07 | 17.29 | 17.36 |
| 372 | CTSC | P53634 | 0.31 | 0.508638306 | 0.17 | 18.21 | 18.38 |
| 373 | CTSD | P07339 | 0.53 | 0.27572413 | -0.03 | 16.11 | 16.08 |
| 374 | CTSL | P07711 | 0.015 | 1.823908741 | -0.46 | 19.29 | 18.83 |
| 375 | CTSZ | Q9UBR2 | 0.019 | 1.721246399 | 0.18 | 19.07 | 19.25 |
| 376 | CTTN | Q14247 | 0.00081 | 3.091514981 | -0.11 | 17.78 | 17.67 |
| 377 | CTTNBP2NL | Q9P2B4 | 0.78 | 0.107905397 | -0.02 | 16.87 | 16.86 |
| 378 | CUL1 | Q13616 | 0.47 | 0.327902142 | -0.08 | 15.61 | 15.53 |
| 379 | CUL2 | Q13617 | 0.0056 | 2.251811973 | 0.28 | 15.75 | 16.03 |
| 380 | CUL3 | Q13618 | 0.29 | 0.537602002 | 0.35 | 12.27 | 12.62 |
| 381 | CUL4A | Q13619 | 0.3 | 0.522878745 | 0.82 | 13.97 | 14.79 |
| 382 | CUL4B | Q13620 | 0.049 | 1.30980392 | -0.09 | 16.79 | 16.7 |
| 383 | CUL5 | Q93034 | 0.88 | 0.055517328 | 0.01 | 14.3 | 14.3 |
| 384 | CUTA | O60888 | 0.39 | 0.408935393 | 0.1 | 14.44 | 14.54 |
| 385 | CYB5B | O43169 | 0.35 | 0.455931956 | 0.03 | 15.56 | 15.59 |
| 386 | CYB5R3 | P00387 | 0.29 | 0.537602002 | 0.1 | 17.01 | 17.11 |
| 387 | CYCS | P99999 | 0.24 | 0.619788758 | 0.14 | 17.33 | 17.47 |
| 388 | CYFIP1 | Q7L576 | 0.2 | 0.698970004 | 0.16 | 16.63 | 16.79 |
| 389 | CYP20A1 | Q6UW02 | 0.069 | 1.161150909 | 0.18 | 14.91 | 15.1 |
| 390 | DAB2 | P98082 | 0.63 | 0.200659451 | 0.02 | 16.26 | 16.28 |
| 391 | DARS | P14868 | 0.87 | 0.060480747 | -0.02 | 17.3 | 17.29 |
| 392 | DARS2 | Q6PI48 | 0.033 | 1.48148606 | -0.13 | 15.49 | 15.37 |
| 393 | DAZAP1 | Q96EP5 | 0.034 | 1.468521083 | 0.12 | 15.33 | 15.45 |
| 394 | DBN1 | Q16643 | 0.00096 | 3.017728767 | 0.11 | 16.1 | 16.21 |
| 395 | DBNL | Q9UJU6 | 0.03 | 1.522878745 | -0.19 | 15.47 | 15.28 |
| 396 | DCAF7 | P61962 | 0.17 | 0.769551079 | 0.11 | 16.34 | 16.45 |
| 397 | DCPS | Q96C86 | 0.98 | 0.008773924 | 0.01 | 16.09 | 16.1 |

| | | | | | | | |
|---|---|---|---|---|---|---|---|
| 398 | DCTN1 | Q14203 | 0.041 | 1.387216143 | -0.08 | 15.34 | 15.26 |
| 399 | DCTN2 | Q13561 | 0.068 | 1.167491087 | -0.17 | 15.84 | 15.67 |
| 400 | DCTN4 | Q9UJW0 | 0.89 | 0.050609993 | 0.03 | 14.69 | 14.72 |
| 401 | DCXR | Q7Z4W1 | 0.61 | 0.214670165 | 0.02 | 15.7 | 15.72 |
| 402 | DDAH1 | O94760 | 0.0057 | 2.244125144 | -0.11 | 17.35 | 17.24 |
| 403 | DDAH2 | O95865 | 0.86 | 0.065501549 | 0 | 17.25 | 17.25 |
| 404 | DDB1 | Q16531 | 0.59 | 0.229147988 | -0.04 | 15.93 | 15.88 |
| 405 | DDI2 | Q5TDH0 | 0.3 | 0.522878745 | -0.12 | 15.34 | 15.21 |
| 406 | DDOST | P39656 | 0.027 | 1.568636236 | 0.14 | 16.33 | 16.48 |
| 407 | DDRGK1 | Q96HY6 | 0.76 | 0.119186408 | 0.04 | 14.16 | 14.19 |
| 408 | DDT | P30046 | 0.33 | 0.48148606 | 0.03 | 15.59 | 15.63 |
| 409 | DDX1 | Q92499 | 0.023 | 1.638272164 | 0.07 | 16.35 | 16.42 |
| 410 | DDX17 | Q92841 | 0.0082 | 2.086186148 | 0.09 | 16.94 | 17.03 |
| 411 | DDX18 | Q9NVP1 | 0.27 | 0.568636236 | 0.3 | 13.6 | 13.9 |
| 412 | DDX19B | Q9UMR2 | 0.58 | 0.236572006 | -0.02 | 17.03 | 17.01 |
| 413 | DDX21 | Q9NR30 | 0.00013 | 3.886056648 | 0.26 | 16.19 | 16.46 |
| 414 | DDX23 | Q9BUQ8 | 0.78 | 0.107905397 | -0.05 | 15.61 | 15.57 |
| 415 | DDX39A | O00148 | 0.083 | 1.080921908 | 0.15 | 16.18 | 16.32 |
| 416 | DDX39B | Q13838 | 0.36 | 0.443697499 | 0.06 | 15.64 | 15.69 |
| 417 | DDX3X | O00571 | 0.0001 | 4 | 0.33 | 16.16 | 16.5 |
| 418 | DDX42 | Q86XP3 | 0.38 | 0.420216403 | 0.06 | 15.63 | 15.68 |
| 419 | DDX46 | Q7L014 | 0.22 | 0.657577319 | 0.12 | 17.19 | 17.31 |
| 420 | DDX5 | P17844 | 0.0042 | 2.37675071 | 0.17 | 17.77 | 17.94 |
| 421 | DDX58 | O95786 | 0.18 | 0.744727495 | -0.12 | 16.58 | 16.46 |
| 422 | DDX6 | P26196 | 0.51 | 0.292429824 | -0.07 | 16.17 | 16.1 |
| 423 | Decoy1 | Q8WZ42-DECOY | 0.61 | 0.214670165 | -0.08 | 21.27 | 21.19 |
| 424 | Decoy10 | Q8WWQ0-DECOY | 0.015 | 1.823908741 | 0.39 | 13.64 | 14.04 |
| 425 | Decoy11 | Q6UB98-DECOY | 0.53 | 0.27572413 | -0.2 | 16.95 | 16.76 |
| 426 | Decoy12 | Q9UI33-DECOY | 0.8 | 0.096910013 | 0.13 | 13.34 | 13.48 |
| 427 | Decoy13 | Q15149-DECOY | 0.53 | 0.27572413 | 0.46 | 14.87 | 15.33 |
| 428 | Decoy14 | Q92616-DECOY | 0.79 | 0.102372909 | -0.18 | 14.46 | 14.28 |
| 429 | Decoy15 | Q7Z628-DECOY | 0.052 | 1.283996656 | 0.46 | 17.15 | 17.61 |
| 430 | Decoy16 | Q9UQ84-DECOY | 0.91 | 0.040958608 | 0.06 | 15.28 | 15.34 |
| 431 | Decoy17 | Q8IWZ3-DECOY | 0.43 | 0.366531544 | 0.17 | 18.11 | 18.27 |
| 432 | Decoy18 | Q9BY77-DECOY | 0.016 | 1.795880017 | 0.16 | 19.89 | 20.05 |
| 433 | Decoy19 | O14686-DECOY | 0.17 | 0.769551079 | -0.39 | 15.61 | 15.22 |
| 434 | Decoy2 | P16401-DECOY | 0.046 | 1.337242168 | -0.35 | 19.42 | 19.07 |
| 435 | Decoy20 | O95714-DECOY | NA | NA | -1.07 | 20.41 | 19.34 |
| 436 | Decoy3 | Q03001-DECOY | 0.028 | 1.552841969 | -0.29 | 17.04 | 16.75 |
| 437 | Decoy4 | Q96G01-DECOY | 0.93 | 0.031517051 | -0.01 | 17.87 | 17.87 |
| 438 | Decoy5 | Q8IZT6-DECOY | 0.18 | 0.744727495 | -0.08 | 18.48 | 18.4 |
| 439 | Decoy6 | P21817-DECOY | NA | NA | 0.72 | 12.76 | 13.48 |
| 440 | Decoy7 | Q3L8U1-DECOY | 0.63 | 0.200659451 | -0.1 | 21.14 | 21.04 |
| 441 | Decoy8 | Q96NL6-DECOY | 0.42 | 0.37675071 | 0.22 | 15.21 | 15.42 |
| 442 | Decoy9 | O15078-DECOY | 0.59 | 0.229147988 | -0.12 | 15.8 | 15.68 |
| 443 | DECR1 | Q16698 | 0.41 | 0.387216143 | 0.05 | 16.75 | 16.8 |
| 444 | DEK | P35659 | 0.74 | 0.13076828 | 0.02 | 16.03 | 16.05 |
| 445 | DENND4C | Q5VZ89 | 0.71 | 0.148741651 | 0.17 | 11.64 | 11.81 |
| 446 | DENR | O43583 | 0.0069 | 2.161150909 | -0.27 | 15.24 | 14.97 |
| 447 | DGLUCY | Q7Z3D6 | 0.015 | 1.823908741 | 0.3 | 15.15 | 15.45 |

| | | | | | | | |
|---|---|---|---|---|---|---|---|
| 448 | DHX15 | O43143 | 0.011 | 1.958607315 | 0.16 | 15.98 | 16.14 |
| 449 | DHX29 | Q7Z478 | 0.17 | 0.769551079 | 0.22 | 13.89 | 14.11 |
| 450 | DHX30 | Q7L2E3 | 0.3 | 0.522878745 | 0.2 | 15.35 | 15.55 |
| 451 | DHX9 | Q08211 | 0.0026 | 2.585026652 | 0.11 | 16.08 | 16.19 |
| 452 | DIAPH1 | O60610 | 0.11 | 0.958607315 | -0.2 | 13.59 | 13.39 |
| 453 | DIS3 | Q9Y2L1 | 0.076 | 1.119186408 | 0.3 | 14.67 | 14.97 |
| 454 | DKC1 | O60832 | 0.012 | 1.920818754 | 0.18 | 15.87 | 16.05 |
| 455 | DLAT | P10515 | 0.058 | 1.236572006 | 0.1 | 16.5 | 16.6 |
| 456 | DLD | P09622 | 0.049 | 1.30980392 | 0.13 | 16.17 | 16.31 |
| 457 | DLST | P36957 | 0.16 | 0.795880017 | 0.12 | 16.94 | 17.06 |
| 458 | DNAJA1 | P31689 | 0.51 | 0.292429824 | 0.06 | 15.93 | 15.99 |
| 459 | DNAJA2 | O60884 | 0.3 | 0.522878745 | -0.16 | 15.6 | 15.44 |
| 460 | DNAJB11 | Q9UBS4 | 0.14 | 0.853871964 | 0.05 | 16.06 | 16.11 |
| 461 | DNAJB4 | Q9UDY4 | 0.0022 | 2.657577319 | -0.21 | 16.64 | 16.43 |
| 462 | DNAJC10 | Q8IXB1 | 0.063 | 1.200659451 | 0.17 | 14.34 | 14.51 |
| 463 | DNAJC13 | O75165 | 0.52 | 0.283996656 | 0.03 | 15.02 | 15.04 |
| 464 | DNAJC3 | Q13217 | 0.0015 | 2.823908741 | -0.12 | 16.55 | 16.42 |
| 465 | DNAJC7 | Q99615 | 0.4 | 0.397940009 | -0.13 | 15.18 | 15.04 |
| 466 | DNM1L | O00429 | 0.14 | 0.853871964 | 0.11 | 15.29 | 15.39 |
| 467 | DNM2 | P50570 | 0.28 | 0.552841969 | 0.05 | 15.56 | 15.6 |
| 468 | DNMBP | Q6XZF7 | 0.33 | 0.48148606 | 0.17 | 13.63 | 13.8 |
| 469 | DNTTIP2 | Q5QJE6 | 0.85 | 0.070581074 | 0.07 | 17.11 | 17.18 |
| 470 | DOCK10 | Q96BY6 | 0.3 | 0.522878745 | 0.33 | 15.16 | 15.49 |
| 471 | DOCK4 | Q8N1I0 | 0.13 | 0.886056648 | 0.12 | 14.92 | 15.04 |
| 472 | DOCK6 | Q96HP0 | 0.7 | 0.15490196 | 0.04 | 14.95 | 14.99 |
| 473 | DOCK9 | Q9BZ29 | 0.33 | 0.48148606 | 0.13 | 15.45 | 15.58 |
| 474 | DPP3 | Q9NY33 | 0.043 | 1.366531544 | -0.15 | 15.67 | 15.52 |
| 475 | DPP7 | Q9UHL4 | 0.31 | 0.508638306 | 0.17 | 15.33 | 15.5 |
| 476 | DPP9 | Q86TI2 | 0.27 | 0.568636236 | 0.1 | 15.47 | 15.56 |
| 477 | DPY19L1 | Q2PZI1 | 0.8 | 0.096910013 | 0.09 | 15.07 | 15.15 |
| 478 | DPYSL2 | Q16555 | 0.035 | 1.455931956 | -0.09 | 16.48 | 16.4 |
| 479 | DPYSL3 | Q14195 | 0.91 | 0.040958608 | 0 | 17.08 | 17.07 |
| 480 | DRG1 | Q9Y295 | 0.56 | 0.251811973 | 0.03 | 15.86 | 15.89 |
| 481 | DRG2 | P55039 | 0.3 | 0.522878745 | 0.14 | 15.67 | 15.81 |
| 482 | DST | Q03001 | 0.43 | 0.366531544 | 0.28 | 13.59 | 13.86 |
| 483 | DSTN | P60981 | 0.38 | 0.420216403 | 0.04 | 16.55 | 16.59 |
| 484 | DTX3L | Q8TDB6 | 0.06 | 1.22184875 | 0.06 | 15.03 | 15.09 |
| 485 | DTYMK | P23919 | 0.064 | 1.193820026 | -0.16 | 16.54 | 16.38 |
| 486 | DUT | P33316 | 0.14 | 0.853871964 | -0.49 | 14.34 | 13.85 |
| 487 | DYNC1H1 | Q14204 | 0.0001 | 4 | 0.08 | 15.78 | 15.86 |
| 488 | DYNC1I2 | Q13409 | 0.75 | 0.124938737 | -0.02 | 16.15 | 16.13 |
| 489 | DYNC1LI1 | Q9Y6G9 | 0.68 | 0.167491087 | 0.06 | 16.62 | 16.68 |
| 490 | DYNC1LI2 | O43237 | 0.43 | 0.366531544 | -0.17 | 16.39 | 16.22 |
| 491 | DYNLL1 | P63167 | 0.043 | 1.366531544 | 0.13 | 17.21 | 17.34 |
| 492 | DYNLRB1 | Q9NP97 | 0.3 | 0.522878745 | -0.09 | 16.6 | 16.52 |
| 493 | DYNLT1 | P63172 | 0.42 | 0.37675071 | 0.09 | 15.39 | 15.48 |
| 494 | DYSF | O75923 | 0.00025 | 3.602059991 | -0.1 | 16.53 | 16.44 |
| 495 | EBNA1BP2 | Q99848 | 0.3 | 0.522878745 | -0.02 | 16.85 | 16.83 |
| 496 | ECE1 | P42892 | 0.13 | 0.886056648 | -0.07 | 15.84 | 15.78 |
| 497 | ECH1 | Q13011 | 0.31 | 0.508638306 | 0.08 | 17.55 | 17.63 |

| | | | | | | | |
|---|---|---|---|---|---|---|---|
| 498 | ECHS1 | P30084 | 0.83 | 0.080921908 | -0.02 | 16.66 | 16.64 |
| 499 | ECI2 | O75521 | 0.069 | 1.161150909 | 0.15 | 16.56 | 16.7 |
| 500 | ECPAS | Q5VYK3 | 0.005 | 2.301029996 | 0.17 | 16.03 | 16.21 |
| 501 | EDC3 | Q96F86 | 0.51 | 0.292429824 | 0.12 | 16.29 | 16.41 |
| 502 | EDC4 | Q6P2E9 | 0.34 | 0.468521083 | 0.16 | 13.95 | 14.11 |
| 503 | EDF1 | O60869 | 0.14 | 0.853871964 | -0.36 | 18.41 | 18.05 |
| 504 | EEA1 | Q15075 | 0.39 | 0.408935393 | -0.04 | 16.81 | 16.77 |
| 505 | EEF1A1 | P68104 | 0.51 | 0.292429824 | -0.03 | 16.69 | 16.66 |
| 506 | EEF1B2 | P24534 | 0.55 | 0.259637311 | -0.01 | 17.4 | 17.39 |
| 507 | EEF1D | P29692 | 0.9 | 0.045757491 | 0 | 16.78 | 16.79 |
| 508 | EEF1E1 | O43324 | 0.95 | 0.022276395 | 0 | 14.12 | 14.11 |
| 509 | EEF1G | P26641 | 0.001 | 3 | 0.14 | 16.61 | 16.76 |
| 510 | EEF2 | P13639 | 0.0074 | 2.13076828 | 0.07 | 16.91 | 16.98 |
| 511 | EFEMP1 | Q12805 | 0.79 | 0.102372909 | -0.06 | 16.03 | 15.96 |
| 512 | EFHD2 | Q96C19 | 0.87 | 0.060480747 | -0.01 | 17.27 | 17.27 |
| 513 | EFL1 | Q7Z2Z2 | 0.65 | 0.187086643 | 0.07 | 14.69 | 14.76 |
| 514 | EFTUD2 | Q15029 | 0.81 | 0.091514981 | 0.01 | 15.98 | 15.99 |
| 515 | EHD1 | Q9H4M9 | 0.51 | 0.292429824 | 0.04 | 16.06 | 16.1 |
| 516 | EHD2 | Q9NZN4 | 0.0001 | 4 | -0.21 | 15.68 | 15.47 |
| 517 | EHD4 | Q9H223 | 0.00021 | 3.677780705 | -0.19 | 17.06 | 16.87 |
| 518 | EIF1 | P41567 | 0.22 | 0.657577319 | -0.14 | 15.49 | 15.35 |
| 519 | EIF1AX | P47813 | 0.039 | 1.408935393 | -0.13 | 18.58 | 18.46 |
| 520 | EIF2A | Q9BY44 | 0.36 | 0.443697499 | 0.04 | 17.08 | 17.13 |
| 521 | EIF2AK2 | P19525 | 0.98 | 0.008773924 | 0 | 15.46 | 15.46 |
| 522 | EIF2B1 | Q14232 | 0.86 | 0.065501549 | 0.01 | 15.93 | 15.94 |
| 523 | EIF2B3 | Q9NR50 | 0.72 | 0.142667504 | -0.03 | 17.7 | 17.67 |
| 524 | EIF2B4 | Q9UI10 | 0.16 | 0.795880017 | -0.12 | 15.22 | 15.1 |
| 525 | EIF2B5 | Q13144 | 0.25 | 0.602059991 | 0.14 | 15 | 15.14 |
| 526 | EIF2S1 | P05198 | 0.2 | 0.698970004 | 0.08 | 16.21 | 16.28 |
| 527 | EIF2S2 | P20042 | 0.22 | 0.657577319 | 0.06 | 16.94 | 17.01 |
| 528 | EIF2S3 | P41091 | 0.0002 | 3.698970004 | 0.2 | 17.19 | 17.38 |
| 529 | EIF3A | Q14152 | 0.9 | 0.045757491 | -0.01 | 15.48 | 15.47 |
| 530 | EIF3B | P55884 | 0.52 | 0.283996656 | -0.03 | 17.2 | 17.18 |
| 531 | EIF3C | Q99613 | 0.54 | 0.26760624 | -0.03 | 16.94 | 16.9 |
| 532 | EIF3D | O15371 | 0.1 | 1 | 0.23 | 14.62 | 14.86 |
| 533 | EIF3E | P60228 | 0.43 | 0.366531544 | 0.11 | 15.37 | 15.47 |
| 534 | EIF3F | O00303 | 0.59 | 0.229147988 | 0.09 | 16.34 | 16.43 |
| 535 | EIF3G | O75821 | 0.028 | 1.552841969 | -0.3 | 17.96 | 17.67 |
| 536 | EIF3H | O15372 | 0.37 | 0.431798276 | -0.13 | 15.25 | 15.11 |
| 537 | EIF3I | Q13347 | 0.73 | 0.13667714 | -0.02 | 18.76 | 18.75 |
| 538 | EIF3J | O75822 | 0.018 | 1.744727495 | -0.2 | 16.52 | 16.32 |
| 539 | EIF3L | Q9Y262 | 0.37 | 0.431798276 | -0.11 | 16 | 15.89 |
| 540 | EIF3M | Q7L2H7 | 0.22 | 0.657577319 | 0.1 | 15.49 | 15.59 |
| 541 | EIF4A1 | P60842 | 0.47 | 0.327902142 | -0.04 | 16.97 | 16.93 |
| 542 | EIF4A2 | Q14240 | 0.57 | 0.244125144 | 0.04 | 16.75 | 16.79 |
| 543 | EIF4A3 | P38919 | 0.11 | 0.958607315 | 0.05 | 16.12 | 16.18 |
| 544 | EIF4B | P23588 | 0.00022 | 3.657577319 | -0.24 | 17.45 | 17.22 |
| 545 | EIF4E | P06730 | 0.98 | 0.008773924 | 0.02 | 16.33 | 16.35 |
| 546 | EIF4G1 | Q04637 | 0.18 | 0.744727495 | 0.05 | 17.08 | 17.12 |
| 547 | EIF4G2 | P78344 | 0.87 | 0.060480747 | 0.01 | 16.24 | 16.25 |

| | | | | | | | |
|---|---|---|---|---|---|---|---|
| 548 | EIF4H | Q15056 | 0.0014 | 2.853871964 | -0.28 | 17.56 | 17.27 |
| 549 | EIF5 | P55010 | 0.91 | 0.040958608 | 0.01 | 15.74 | 15.74 |
| 550 | EIF5A | P63241 | 0.048 | 1.318758763 | -0.07 | 17.34 | 17.26 |
| 551 | EIF5B | O60841 | 0.0001 | 4 | 0.17 | 16.52 | 16.69 |
| 552 | EIF6 | P56537 | 0.23 | 0.638272164 | -0.09 | 17.04 | 16.94 |
| 553 | ELAC2 | Q9BQ52 | 0.56 | 0.251811973 | 0.06 | 13.81 | 13.87 |
| 554 | ELAVL1 | Q15717 | 0.09 | 1.045757491 | 0.25 | 17.47 | 17.73 |
| 555 | ELMO1 | Q92556 | 0.2 | 0.698970004 | 0.23 | 16.06 | 16.3 |
| 556 | ELOB | Q15370 | 0.38 | 0.420216403 | 0.1 | 16.81 | 16.91 |
| 557 | ELOC | Q15369 | 0.9 | 0.045757491 | -0.01 | 17.33 | 17.32 |
| 558 | ELOVL1 | Q9BW60 | 0.14 | 0.853871964 | -0.14 | 17.87 | 17.73 |
| 559 | ELP3 | Q9H9T3 | 0.66 | 0.180456064 | 0.1 | 15.06 | 15.16 |
| 560 | EMC1 | Q8N766 | 0.93 | 0.031517051 | 0 | 15.23 | 15.23 |
| 561 | EMD | P50402 | 0.17 | 0.769551079 | -0.09 | 18.67 | 18.58 |
| 562 | EMG1 | Q92979 | 0.67 | 0.173925197 | 0.06 | 15.2 | 15.26 |
| 563 | EML1 | O00423 | 0.1 | 1 | 0.36 | 16.14 | 16.5 |
| 564 | ENDOD1 | O94919 | 0.0026 | 2.585026652 | -0.18 | 14.48 | 14.3 |
| 565 | ENG | P17813 | 0.71 | 0.148741651 | -0.02 | 17.19 | 17.17 |
| 566 | ENO1 | P06733 | 0.0038 | 2.420216403 | -0.11 | 17.06 | 16.95 |
| 567 | ENOPH1 | Q9UHY7 | 0.12 | 0.920818754 | 0.23 | 15.08 | 15.3 |
| 568 | EOGT | Q5NDL2 | 0.21 | 0.677780705 | 0.31 | 15.6 | 15.91 |
| 569 | EPB41L2 | O43491 | 0.2 | 0.698970004 | -0.05 | 16.64 | 16.59 |
| 570 | EPHA2 | P29317 | 0.7 | 0.15490196 | 0.05 | 15.42 | 15.47 |
| 571 | EPN1 | Q9Y6I3 | 0.2 | 0.698970004 | 0.22 | 15.59 | 15.82 |
| 572 | EPRS | P07814 | 0.8 | 0.096910013 | -0.04 | 16.59 | 16.55 |
| 573 | EPS15 | P42566 | 0.025 | 1.602059991 | -0.23 | 14.94 | 14.7 |
| 574 | EPS15L1 | Q9UBC2 | 0.21 | 0.677780705 | -0.11 | 13.88 | 13.77 |
| 575 | ERAP1 | Q9NZ08 | 0.12 | 0.920818754 | -0.07 | 15.25 | 15.18 |
| 576 | ERC1 | Q8IUD2 | 0.21 | 0.677780705 | -0.4 | 18.12 | 17.72 |
| 577 | ERG | P11308 | 0.61 | 0.214670165 | -0.09 | 16.82 | 16.73 |
| 578 | ERGIC1 | Q969X5 | 0.41 | 0.387216143 | 0.06 | 16.82 | 16.88 |
| 579 | ERGIC2 | Q96RQ1 | 0.71 | 0.148741651 | -0.03 | 16.74 | 16.7 |
| 580 | ERLIN1 | O75477 | 0.017 | 1.769551079 | -0.22 | 16.82 | 16.59 |
| 581 | ERLIN2 | O94905 | 0.13 | 0.886056648 | -0.11 | 15.83 | 15.72 |
| 582 | ERO1A | Q96HE7 | 0.015 | 1.823908741 | 0.21 | 15.89 | 16.11 |
| 583 | ERP29 | P30040 | 0.87 | 0.060480747 | 0.01 | 16.55 | 16.56 |
| 584 | ERP44 | Q9BS26 | 0.58 | 0.236572006 | -0.02 | 16.58 | 16.55 |
| 585 | ESAM | Q96AP7 | 0.41 | 0.387216143 | -0.16 | 17.01 | 16.85 |
| 586 | ESD | P10768 | 0.34 | 0.468521083 | 0.04 | 16.3 | 16.34 |
| 587 | ESM1 | Q9NQ30 | 0.6 | 0.22184875 | 0.18 | 16.85 | 17.03 |
| 588 | ESYT1 | Q9BSJ8 | 1 | 0 | 0 | 15.72 | 15.72 |
| 589 | ESYT2 | A0FGR8 | 0.86 | 0.065501549 | -0.01 | 15.72 | 15.7 |
| 590 | ETF1 | P62495 | 0.00065 | 3.187086643 | 0.17 | 16.06 | 16.23 |
| 591 | ETFA | P13804 | 0.57 | 0.244125144 | 0.01 | 15.5 | 15.51 |
| 592 | ETFB | P38117 | 0.34 | 0.468521083 | -0.04 | 17.31 | 17.27 |
| 593 | ETHE1 | O95571 | 0.12 | 0.920818754 | 0.21 | 15.72 | 15.93 |
| 594 | EWSR1 | Q01844 | 0.11 | 0.958607315 | -0.15 | 16.95 | 16.8 |
| 595 | EXOC1 | Q9NV70 | 0.92 | 0.036212173 | 0.05 | 14.74 | 14.8 |
| 596 | EXOC2 | Q96KP1 | 0.14 | 0.853871964 | 0.17 | 15.61 | 15.79 |
| 597 | EXOC4 | Q96A65 | 0.94 | 0.026872146 | -0.01 | 16.34 | 16.33 |

| | | | | | | | | |
|---|---|---|---|---|---|---|---|---|
| 598 | EXOC5 | O00471 | 0.92 | 0.036212173 | | -0.01 | 14.98 | 14.97 |
| 599 | EXOSC10 | Q01780 | 0.095 | 1.022276395 | | 0.24 | 17.2 | 17.44 |
| 600 | EXOSC7 | Q15024 | 0.97 | 0.013228266 | | 0.01 | 14.66 | 14.67 |
| 601 | EZR | P15311 | 0.012 | 1.920818754 | | -0.17 | 16.09 | 15.92 |
| 602 | F11R | Q9Y624 | 0.12 | 0.920818754 | | 0.08 | 16.14 | 16.22 |
| 603 | FABP4 | P15090 | 0.84 | 0.075720714 | | -0.03 | 16.78 | 16.75 |
| 604 | FABP5 | Q01469 | 0.0001 | 4 | 0.28 | 17.91 | 18.18 | |
| 605 | FAF2 | Q96CS3 | 0.76 | 0.119186408 | | -0.02 | 16.78 | 16.75 |
| 606 | FAH | P16930 | 0.47 | 0.327902142 | | -0.03 | 13.47 | 13.44 |
| 607 | FAHD1 | Q6P587 | 0.27 | 0.568636236 | | 0.11 | 16.87 | 16.98 |
| 608 | FAHD2A | Q96GK7 | 0.13 | 0.886056648 | | 0.08 | 17.12 | 17.21 |
| 609 | FAM114A1 | | Q8IWE2 | 0.066 | 1.180456064 | 0.16 | 15.35 | 15.5 |
| 610 | FAM114A2 | | Q9NRY5 | 0.2 | 0.698970004 | -0.17 | 14.04 | 13.87 |
| 611 | FAM120A | Q9NZB2 | 0.11 | 0.958607315 | | 0.15 | 15.23 | 15.38 |
| 612 | FAM160B1 | | Q5W0V3 | 0.23 | 0.638272164 | 0.19 | 15.78 | 15.97 |
| 613 | FAM186A | A6NE01 | 0.16 | 0.795880017 | | -0.21 | 16.24 | 16.04 |
| 614 | FAM3C | Q92520 | 0.023 | 1.638272164 | | 0.1 | 16.56 | 16.65 |
| 615 | FAM49B | Q9NUQ9 | 0.78 | 0.107905397 | | 0.03 | 15.33 | 15.35 |
| 616 | FAM98A | Q8NCA5 | 0.26 | 0.585026652 | | 0.13 | 14.53 | 14.66 |
| 617 | FAM98B | Q52LJ0 | 0.39 | 0.408935393 | | -0.07 | 16.07 | 16 |
| 618 | FARSA | Q9Y285 | 0.068 | 1.167491087 | | -0.09 | 17.64 | 17.55 |
| 619 | FARSB | Q9NSD9 | 0.76 | 0.119186408 | | 0.02 | 17.17 | 17.19 |
| 620 | FASN | P49327 | 0.0001 | 4 | 0.16 | 16.15 | 16.31 | |
| 621 | FAU | P62861 | 0.28 | 0.552841969 | | 0.14 | 19.54 | 19.68 |
| 622 | FBL | P22087 | 0.18 | 0.744727495 | | 0.14 | 16.59 | 16.74 |
| 623 | FBLIM1 | Q8WUP2 | 0.26 | 0.585026652 | | 0.09 | 17.72 | 17.81 |
| 624 | FDX1 | P10109 | 0.34 | 0.468521083 | | 0.19 | 14.74 | 14.93 |
| 625 | FDXR | P22570 | 0.036 | 1.443697499 | | 0.29 | 15.72 | 16.01 |
| 626 | FEN1 | P39748 | 0.71 | 0.148741651 | | 0.05 | 13.8 | 13.85 |
| 627 | FERMT2 | Q96AC1 | 0.0037 | 2.431798276 | | 0.18 | 15.39 | 15.57 |
| 628 | FERMT3 | Q86UX7 | 0.12 | 0.920818754 | | 0.18 | 15.2 | 15.38 |
| 629 | FH | P07954 | 0.059 | 1.229147988 | | 0.08 | 16.6 | 16.68 |
| 630 | FHL1 | Q13642 | 0.22 | 0.657577319 | | 0.18 | 17.41 | 17.6 |
| 631 | FHL2 | Q14192 | 0.0039 | 2.408935393 | | 0.28 | 17.32 | 17.59 |
| 632 | FHL3 | Q13643 | 0.34 | 0.468521083 | | -0.27 | 17.01 | 16.74 |
| 633 | FHOD1 | Q9Y613 | 0.63 | 0.200659451 | | 0.04 | 12.89 | 12.93 |
| 634 | FIP1L1 | Q6UN15 | 0.35 | 0.455931956 | | -0.14 | 15.15 | 15.01 |
| 635 | FIS1 | Q9Y3D6 | 0.4 | 0.397940009 | | -0.11 | 14.98 | 14.87 |
| 636 | FKBP10 | Q96AY3 | 0.014 | 1.853871964 | | -0.16 | 15.79 | 15.63 |
| 637 | FKBP11 | Q9NYL4 | 0.15 | 0.823908741 | | -0.07 | 17.22 | 17.15 |
| 638 | FKBP15 | Q5T1M5 | 0.25 | 0.602059991 | | 0.1 | 14.77 | 14.86 |
| 639 | FKBP1A | P62942 | 0.17 | 0.769551079 | | -0.15 | 18.13 | 17.98 |
| 640 | FKBP3 | Q00688 | 0.00045 | 3.346787486 | | -0.12 | 17.03 | 16.92 |
| 641 | FKBP4 | Q02790 | 0.051 | 1.292429824 | | -0.07 | 16.41 | 16.34 |
| 642 | FKBP5 | Q13451 | 0.84 | 0.075720714 | | -0.01 | 17.13 | 17.12 |
| 643 | FKBP8 | Q14318 | 0.22 | 0.657577319 | | -0.07 | 15.65 | 15.58 |
| 644 | FKBP9 | O95302 | 0.41 | 0.387216143 | | 0.04 | 16.44 | 16.47 |
| 645 | FLII | Q13045 | 0.98 | 0.008773924 | | -0.01 | 15.22 | 15.21 |
| 646 | FLNA | P21333 | 0.0001 | 4 | 0.1 | 17.1 | 17.2 | |
| 647 | FLNB | O75369 | 0.0001 | 4 | 0.07 | 17.12 | 17.19 | |

| | | | | | | | |
|---|---|---|---|---|---|---|---|
| 648 | FLNC | Q14315 | 0.0001 | 4 | 0.22 | 17.19 | 17.41 |
| 649 | FLOT1 | O75955 | 0.05 | 1.301029996 | -0.16 | 15.14 | 14.98 |
| 650 | FLOT2 | Q14254 | 0.53 | 0.27572413 | 0.07 | 15.28 | 15.34 |
| 651 | FN1 | P02751 | 0.00048 | 3.318758763 | 0.36 | 15.38 | 15.74 |
| 652 | FNBP1L | Q5T0N5 | 0.39 | 0.408935393 | -0.07 | 16.95 | 16.88 |
| 653 | FNDC3B | Q53EP0 | 0.95 | 0.022276395 | 0 | 16.75 | 16.74 |
| 654 | FOCAD | Q5VW36 | 0.032 | 1.494850022 | 0.26 | 14.64 | 14.9 |
| 655 | FRYL | O94915 | 0.7 | 0.15490196 | 0.08 | 16.78 | 16.85 |
| 656 | FSCN1 | Q16658 | 0.74 | 0.13076828 | -0.01 | 16.4 | 16.39 |
| 657 | FSTL1 | Q12841 | 0.88 | 0.055517328 | 0.03 | 17.73 | 17.76 |
| 658 | FTO | Q9C0B1 | 0.68 | 0.167491087 | 0.03 | 15.57 | 15.6 |
| 659 | FTSJ3 | Q8IY81 | 0.35 | 0.455931956 | -0.04 | 14.41 | 14.37 |
| 660 | FUBP1 | Q96AE4 | 0.44 | 0.356547324 | -0.03 | 18.19 | 18.16 |
| 661 | FUS | P35637 | 1 | 0 | -0.02 | 16.87 | 16.85 |
| 662 | FXR1 | P51114 | 0.24 | 0.619788758 | -0.1 | 17.14 | 17.04 |
| 663 | FYTTD1 | Q96QD9 | 0.21 | 0.677780705 | -0.18 | 16.89 | 16.71 |
| 664 | G3BP1 | Q13283 | 0.75 | 0.124938737 | -0.02 | 16.28 | 16.26 |
| 665 | G3BP2 | Q9UN86 | 0.76 | 0.119186408 | 0.03 | 15.38 | 15.41 |
| 666 | G6PD | P11413 | 0.0001 | 4 | -0.22 | 16.36 | 16.14 |
| 667 | GAK | O14976 | 0.38 | 0.420216403 | 0.12 | 16.23 | 16.35 |
| 668 | GALE | Q14376 | 0.58 | 0.236572006 | -0.08 | 14.39 | 14.31 |
| 669 | GALK1 | P51570 | 0.058 | 1.236572006 | -0.19 | 16.25 | 16.06 |
| 670 | GALNT1 | Q10472 | 0.95 | 0.022276395 | -0.01 | 16.63 | 16.62 |
| 671 | GALNT2 | Q10471 | 0.9 | 0.045757491 | -0.03 | 14.72 | 14.7 |
| 672 | GANAB | Q14697 | 0.0009 | 3.045757491 | 0.11 | 16.42 | 16.53 |
| 673 | GAPDH | P04406 | 0.83 | 0.080921908 | -0.01 | 16.72 | 16.71 |
| 674 | GARS | P41250 | 0.8 | 0.096910013 | 0.02 | 16 | 16.02 |
| 675 | GART | P22102 | 0.00022 | 3.657577319 | 0.19 | 15.53 | 15.72 |
| 676 | GATD3B | A0A0B4J2D5 (+1) | 0.85 | 0.070581074 | -0.04 | 14.46 | 14.42 |
| 677 | GBA | P04062 | 0.91 | 0.040958608 | -0.03 | 14.84 | 14.81 |
| 678 | GBE1 | Q04446 | 0.5 | 0.301029996 | -0.03 | 16.55 | 16.53 |
| 679 | GBF1 | Q92538 | 0.97 | 0.013228266 | 0 | 15.74 | 15.74 |
| 680 | GBP1 | P32455 | 0.2 | 0.698970004 | -0.09 | 16.72 | 16.64 |
| 681 | GBP2 | P32456 | 0.35 | 0.455931956 | -0.07 | 16.38 | 16.31 |
| 682 | GCLM | P48507 | 0.87 | 0.060480747 | -0.04 | 17.23 | 17.19 |
| 683 | GCN1 | Q92616 | 0.23 | 0.638272164 | 0.04 | 15.45 | 15.48 |
| 684 | GCSH | P23434 | 0.92 | 0.036212173 | -0.02 | 15.37 | 15.35 |
| 685 | GDI1 | P31150 | 0.91 | 0.040958608 | 0 | 14.6 | 14.6 |
| 686 | GDI2 | P50395 | 0.0091 | 2.040958608 | 0.14 | 16.19 | 16.33 |
| 687 | GEMIN5 | Q8TEQ6 | 0.33 | 0.48148606 | 0.28 | 11.77 | 12.05 |
| 688 | GFM1 | Q96RP9 | 0.44 | 0.356547324 | 0.09 | 15.5 | 15.59 |
| 689 | GFM2 | Q969S9 | 0.47 | 0.327902142 | 0.15 | 15.62 | 15.77 |
| 690 | GFPT1 | Q06210 | 0.42 | 0.37675071 | -0.08 | 16.17 | 16.09 |
| 691 | GGA1 | Q9UJY5 | 0.18 | 0.744727495 | 0.07 | 14.09 | 14.16 |
| 692 | GGA3 | Q9NZ52 | 0.32 | 0.494850022 | -0.21 | 14.39 | 14.18 |
| 693 | GGCT | O75223 | 0.13 | 0.886056648 | -0.06 | 16.39 | 16.33 |
| 694 | GGH | Q92820 | 0.058 | 1.236572006 | 0.44 | 16.27 | 16.72 |
| 695 | GIGYF2 | Q6Y7W6 | 0.96 | 0.017728767 | 0.02 | 14.42 | 14.43 |
| 696 | GIMAP1 | Q8WWP7 | 0.25 | 0.602059991 | -0.09 | 16.1 | 16.01 |
| 697 | GIMAP4 | Q9NUV9 | 0.33 | 0.48148606 | -0.07 | 15.26 | 15.19 |

| | | | | | | | |
|---|---|---|---|---|---|---|---|
| 698 | GIMAP8 | Q8ND71 | 0.79 | 0.102372909 | -0.01 | 14.97 | 14.96 |
| 699 | GLB1 | P16278 | 0.24 | 0.619788758 | 0.1 | 15.73 | 15.83 |
| 700 | GLCE | O94923 | 0.96 | 0.017728767 | -0.01 | 14.18 | 14.17 |
| 701 | GLG1 | Q92896 | 0.00079 | 3.102372909 | -0.18 | 16.54 | 16.36 |
| 702 | GLO1 | Q04760 | 0.011 | 1.958607315 | 0.28 | 16.51 | 16.79 |
| 703 | GLOD4 | Q9HC38 | 0.67 | 0.173925197 | 0.02 | 16.38 | 16.4 |
| 704 | GLRX3 | O76003 | 0.44 | 0.356547324 | -0.04 | 16.77 | 16.72 |
| 705 | GLS | O94925 | 0.12 | 0.920818754 | 0.14 | 16.75 | 16.9 |
| 706 | GLUD1 | P00367 | 0.013 | 1.886056648 | 0.09 | 16.8 | 16.89 |
| 707 | GMFG | O60234 | 0.07 | 1.15490196 | -0.11 | 15.37 | 15.26 |
| 708 | GMPPA | Q96IJ6 | 0.035 | 1.455931956 | -0.18 | 15.99 | 15.81 |
| 709 | GMPS | P49915 | 0.045 | 1.346787486 | 0.13 | 15.51 | 15.64 |
| 710 | GNAI2 | P04899 | 0.08 | 1.096910013 | 0.07 | 16.13 | 16.2 |
| 711 | GNAQ | P50148 | 0.26 | 0.585026652 | 0.07 | 15.43 | 15.5 |
| 712 | GNAS | P63092 | 0.42 | 0.37675071 | -0.06 | 15.89 | 15.83 |
| 713 | GNB1 | P62873 | 0.99 | 0.004364805 | 0 | 16.92 | 16.92 |
| 714 | GNB2 | P62879 | 0.28 | 0.552841969 | 0.07 | 16.22 | 16.29 |
| 715 | GNB4 | Q9HAV0 | 0.92 | 0.036212173 | 0 | 15.92 | 15.93 |
| 716 | GNG12 | Q9UBI6 | 0.61 | 0.214670165 | 0.07 | 17.16 | 17.23 |
| 717 | GNL1 | P36915 | 0.77 | 0.113509275 | 0.04 | 14.82 | 14.86 |
| 718 | GNL2 | Q13823 | 0.4 | 0.397940009 | 0.2 | 13.95 | 14.15 |
| 719 | GNL3 | Q9BVP2 | 0.3 | 0.522878745 | 0.18 | 15.79 | 15.97 |
| 720 | GNPDA1 | P46926 | 0.98 | 0.008773924 | 0 | 16.56 | 16.56 |
| 721 | GNPNAT1 | Q96EK6 | 0.43 | 0.366531544 | 0.11 | 16.05 | 16.15 |
| 722 | GNS | P15586 | 0.0001 | 4 0.2 | 16.63 | 16.83 | |
| 723 | GOLGA3 | Q08378 | 0.55 | 0.259637311 | 0.05 | 14.14 | 14.19 |
| 724 | GORASP2 | Q9H8Y8 | 0.19 | 0.721246399 | 0.07 | 16.81 | 16.88 |
| 725 | GOT1 | P17174 | 0.055 | 1.259637311 | 0.14 | 15.98 | 16.12 |
| 726 | GOT2 | P00505 | 0.0001 | 4 0.18 | 17.27 | 17.45 | |
| 727 | GPD2 | P43304 | 0.012 | 1.920818754 | 0.16 | 15.09 | 15.24 |
| 728 | GPI | P06744 | 0.17 | 0.769551079 | 0.05 | 17.06 | 17.1 |
| 729 | GPS1 | Q13098 | 0.38 | 0.420216403 | 0.24 | 17.73 | 17.97 |
| 730 | GPX1 | P07203 | 0.0097 | 2.013228266 | -0.14 | 16.1 | 15.96 |
| 731 | GPX4 | P36969 | 0.006 | 2.22184875 | -0.27 | 17.31 | 17.03 |
| 732 | GRHPR | Q9UBQ7 | 0.11 | 0.958607315 | -0.14 | 16.93 | 16.79 |
| 733 | GRN | P28799 | 0.94 | 0.026872146 | -0.01 | 18.04 | 18.04 |
| 734 | GRPEL1 | Q9HAV7 | 0.53 | 0.27572413 | -0.05 | 17.27 | 17.22 |
| 735 | GRSF1 | Q12849 | 0.33 | 0.48148606 | 0.12 | 15.32 | 15.44 |
| 736 | GSDMD | P57764 | 0.13 | 0.886056648 | -0.12 | 16.8 | 16.68 |
| 737 | GSK3B | P49841 | 0.92 | 0.036212173 | 0.02 | 13.78 | 13.79 |
| 738 | GSN | P06396 | 0.00057 | 3.244125144 | 0.19 | 17.83 | 18.02 |
| 739 | GSPT1 | P15170 | 0.045 | 1.346787486 | 0.11 | 16.35 | 16.46 |
| 740 | GSR | P00390 | 0.2 | 0.698970004 | 0.09 | 17.05 | 17.13 |
| 741 | GSS | P48637 | 0.78 | 0.107905397 | 0.03 | 16.03 | 16.06 |
| 742 | GSTM3 | P21266 | 0.068 | 1.167491087 | 0.12 | 14.48 | 14.59 |
| 743 | GSTO1 | P78417 | 0.48 | 0.318758763 | -0.03 | 17.07 | 17.03 |
| 744 | GSTP1 | P09211 | 0.066 | 1.180456064 | 0.05 | 16.27 | 16.33 |
| 745 | GTF2E1 | P29083 | 0.88 | 0.055517328 | -0.01 | 13.97 | 13.96 |
| 746 | GTF2E2 | P29084 | 0.44 | 0.356547324 | -0.12 | 13.88 | 13.77 |
| 747 | GTF2F1 | P35269 | 0.84 | 0.075720714 | 0.01 | 17.14 | 17.15 |

| | | | | | | | |
|---|---|---|---|---|---|---|---|
| 748 | GTF2I | P78347 | 0.88 | 0.055517328 | 0 | 16.45 | 16.46 |
| 749 | GTPBP4 | Q9BZE4 | 0.12 | 0.920818754 | 0.22 | 14.84 | 15.07 |
| 750 | GUK1 | Q16774 | 0.23 | 0.638272164 | 0.19 | 16.88 | 17.07 |
| 751 | GYG1 | P46976 | 0.075 | 1.124938737 | -0.17 | 16.39 | 16.21 |
| 752 | H1F0 | P07305 | 0.013 | 1.886056648 | -0.49 | 17.49 | 16.99 |
| 753 | H1FX | Q92522 | 0.0048 | 2.318758763 | -0.37 | 17.22 | 16.86 |
| 754 | H2AFV | Q71UI9 | 0.21 | 0.677780705 | -0.4 | 17.98 | 17.58 |
| 755 | H2AFY | O75367 | 0.64 | 0.193820026 | 0.02 | 17.15 | 17.17 |
| 756 | HACD3 | Q9P035 | 0.12 | 0.920818754 | 0.17 | 16.67 | 16.84 |
| 757 | HADH | Q16836 | 0.31 | 0.508638306 | -0.12 | 17.29 | 17.16 |
| 758 | HADHA | P40939 | 0.0013 | 2.886056648 | 0.1 | 16.49 | 16.6 |
| 759 | HADHB | P55084 | 0.013 | 1.886056648 | 0.1 | 16.5 | 16.6 |
| 760 | HAGH | Q16775 | 0.32 | 0.494850022 | -0.06 | 15.02 | 14.96 |
| 761 | HARS | P12081 | 0.24 | 0.619788758 | 0.08 | 16.1 | 16.18 |
| 762 | HBS1L | Q9Y450 | 0.43 | 0.366531544 | -0.08 | 16.48 | 16.4 |
| 763 | HCFC1 | P51610 | 0.00024 | 3.619788758 | 0.19 | 17.18 | 17.37 |
| 764 | HCLS1 | P14317 | 0.0026 | 2.585026652 | -0.24 | 18.34 | 18.09 |
| 765 | HDAC1 | Q13547 | 0.3 | 0.522878745 | 0.17 | 16.22 | 16.39 |
| 766 | HDAC2 | Q92769 | 0.027 | 1.568636236 | 0.32 | 15.17 | 15.48 |
| 767 | HDGF | P51858 | 0.0001 | 4 | -0.2 | 16.46 | 16.26 |
| 768 | HDGFL2 | Q7Z4V5 | 0.69 | 0.161150909 | 0.03 | 16.38 | 16.41 |
| 769 | HDGFL3 | Q9Y3E1 | 0.22 | 0.657577319 | -0.07 | 19.34 | 19.26 |
| 770 | HDHD5 | Q9BXW7 | 0.57 | 0.244125144 | -0.09 | 15.25 | 15.16 |
| 771 | HDLBP | Q00341 | 0.023 | 1.638272164 | -0.09 | 15.76 | 15.67 |
| 772 | HEBP1 | Q9NRV9 | 0.26 | 0.585026652 | -0.11 | 14.79 | 14.68 |
| 773 | HEBP2 | Q9Y5Z4 | 0.7 | 0.15490196 | -0.1 | 15.84 | 15.74 |
| 774 | HERC4 | Q5GLZ8 | 0.28 | 0.552841969 | 0.15 | 16.11 | 16.26 |
| 775 | HEXA | P06865 | 0.12 | 0.920818754 | 0.07 | 15.71 | 15.78 |
| 776 | HEXB | P07686 | 0.94 | 0.026872146 | 0 | 16.07 | 16.07 |
| 777 | HGS | O14964 | 0.031 | 1.508638306 | 0.15 | 16.62 | 16.77 |
| 778 | HIBADH | P31937 | 0.34 | 0.468521083 | 0.11 | 15.07 | 15.18 |
| 779 | HIBCH | Q6NVY1 | 0.17 | 0.769551079 | -0.2 | 17.14 | 16.94 |
| 780 | HINT1 | P49773 | 0.018 | 1.744727495 | 0.15 | 16.44 | 16.58 |
| 781 | HINT2 | Q9BX68 | 0.84 | 0.075720714 | 0.02 | 16.55 | 16.57 |
| 782 | HIP1R | O75146 | 0.012 | 1.920818754 | -0.25 | 16.95 | 16.7 |
| 783 | HIST1H1B | P16401 | 0.0001 | 4 | -0.36 | 19.16 | 18.8 |
| 784 | HIST1H1C | P16403 | 0.0001 | 4 | -0.75 | 18.59 | 17.84 |
| 785 | HIST1H1D | P16402 | 0.0001 | 4 | -0.73 | 19.01 | 18.28 |
| 786 | HIST1H1E | P10412 | 0.0011 | 2.958607315 | -0.47 | 19.5 | 19.03 |
| 787 | HIST1H2AJ | Q99878 | 0.48 | 0.318758763 | -0.12 | 16.06 | 15.95 |
| 788 | HIST1H4A | P62805 | 0.77 | 0.113509275 | 0.04 | 18.35 | 18.39 |
| 789 | HIST2H2BF | Q5QNW6 (+1) | 0.0022 | 2.657577319 | -0.33 | 17.36 | 17.03 |
| 790 | HIST2H3A | Q71DI3 | 0.72 | 0.142667504 | -0.04 | 19.26 | 19.22 |
| 791 | HK1 | P19367 | 0.024 | 1.619788758 | 0.08 | 16.86 | 16.95 |
| 792 | HLA-A(1) | P01892 | 0.047 | 1.327902142 | -0.15 | 16.2 | 16.06 |
| 793 | HLA-A(2) | P30512 | 0.59 | 0.229147988 | -0.11 | 16.85 | 16.74 |
| 794 | HLA-B | Q04826 | 0.42 | 0.37675071 | -0.15 | 16.13 | 15.98 |
| 795 | HM13 | Q8TCT9 | 0.74 | 0.13076828 | 0.03 | 18.54 | 18.57 |
| 796 | HMGA1 | P17096 | 0.84 | 0.075720714 | -0.01 | 18.68 | 18.67 |

| | | | | | | | |
|---|---|---|---|---|---|---|---|
| 797 | HMGA2 | P52926 | 0.45 | 0.346787486 | 0.04 | 17.27 | 17.31 |
| 798 | HMGB1 | P09429 | 0.69 | 0.161150909 | -0.01 | 18.79 | 18.78 |
| 799 | HMGB2 | P26583 | 0.0087 | 2.060480747 | -0.2 | 18.51 | 18.31 |
| 800 | HMGB3 | O15347 | NA | NA | -0.1 | 19.45 | 19.35 |
| 801 | HMGCL | P35914 | 0.63 | 0.200659451 | -0.05 | 14.66 | 14.61 |
| 802 | HMGN1 | P05114 | 0.34 | 0.468521083 | 0.07 | 17.09 | 17.16 |
| 803 | HMGN2 | P05204 | 0.2 | 0.698970004 | -0.22 | 17.58 | 17.37 |
| 804 | HMOX2 | P30519 | 0.29 | 0.537602002 | 0.11 | 15.18 | 15.29 |
| 805 | HNRNPA0 | Q13151 | 0.11 | 0.958607315 | -0.23 | 16.89 | 16.66 |
| 806 | HNRNPA1 | P09651 | 0.89 | 0.050609993 | 0 | 17.19 | 17.18 |
| 807 | HNRNPA2B1 | P22626 | 0.48 | 0.318758763 | 0.03 | 17.01 | 17.04 |
| 808 | HNRNPA3 | P51991 | 0.56 | 0.251811973 | 0.03 | 16.46 | 16.5 |
| 809 | HNRNPAB | Q99729 | 0.46 | 0.337242168 | -0.07 | 16.61 | 16.54 |
| 810 | HNRNPC | P07910 | 0.0014 | 2.853871964 | -0.13 | 17.84 | 17.71 |
| 811 | HNRNPD | Q14103 | 0.0085 | 2.070581074 | 0.23 | 13.53 | 13.76 |
| 812 | HNRNPDL | O14979 | 0.27 | 0.568636236 | 0.04 | 17.02 | 17.06 |
| 813 | HNRNPF | P52597 | 0.57 | 0.244125144 | 0.02 | 16.79 | 16.81 |
| 814 | HNRNPH1 | P31943 | 0.53 | 0.27572413 | -0.05 | 17.28 | 17.23 |
| 815 | HNRNPH2 | P55795 | 0.67 | 0.173925197 | -0.06 | 15.33 | 15.27 |
| 816 | HNRNPH3 | P31942 | 0.11 | 0.958607315 | -0.06 | 16.89 | 16.83 |
| 817 | HNRNPK | P61978 | 0.34 | 0.468521083 | -0.04 | 16.78 | 16.75 |
| 818 | HNRNPL | P14866 | 0.032 | 1.494850022 | 0.1 | 17.01 | 17.11 |
| 819 | HNRNPLL | Q8WVV9 | 0.14 | 0.853871964 | 0.37 | 14.1 | 14.47 |
| 820 | HNRNPM | P52272 | 0.0079 | 2.102372909 | 0.13 | 16.18 | 16.31 |
| 821 | HNRNPR | O43390 | 0.89 | 0.050609993 | 0 | 15.77 | 15.76 |
| 822 | HNRNPU | Q00839 | 0.71 | 0.148741651 | -0.03 | 16.77 | 16.74 |
| 823 | HNRNPUL1 | Q9BUJ2 | 0.48 | 0.318758763 | 0.09 | 17.6 | 17.69 |
| 824 | HNRNPUL2 | Q1KMD3 | 0.094 | 1.026872146 | 0.1 | 16.07 | 16.16 |
| 825 | HOMER3 | Q9NSC5 | 0.58 | 0.236572006 | -0.04 | 15.79 | 15.75 |
| 826 | HP1BP3 | Q5SSJ5 | 0.71 | 0.148741651 | -0.03 | 16.23 | 16.2 |
| 827 | HRNR | Q86YZ3 | 0.39 | 0.408935393 | 0.06 | 12.25 | 12.31 |
| 828 | HSD17B10 | Q99714 | 0.013 | 1.886056648 | 0.15 | 15.7 | 15.85 |
| 829 | HSD17B11 | Q8NBQ5 | 0.018 | 1.744727495 | 0.2 | 16.09 | 16.28 |
| 830 | HSD17B12 | Q53GQ0 | 0.13 | 0.886056648 | -0.1 | 15.55 | 15.44 |
| 831 | HSD17B4 | P51659 | 0.4 | 0.397940009 | 0.04 | 16.35 | 16.39 |
| 832 | HSDL2 | Q6YN16 | 0.25 | 0.602059991 | 0.23 | 15.6 | 15.83 |
| 833 | HSP90AA1 | P07900 | 0.082 | 1.086186148 | 0.07 | 17 | 17.07 |
| 834 | HSP90AB1 | P08238 | 0.78 | 0.107905397 | -0.03 | 16.85 | 16.82 |
| 835 | HSP90B1 | P14625 | 0.00011 | 3.958607315 | -0.1 | 17.25 | 17.15 |
| 836 | HSPA1A | P0DMV8 (+1) | 0.00074 | 3.13076828 | -0.13 | 16.41 | 16.28 |
| 837 | HSPA4 | P34932 | 0.0074 | 2.13076828 | 0.07 | 16.4 | 16.48 |
| 838 | HSPA5 | P11021 | 0.0082 | 2.086186148 | -0.1 | 17.26 | 17.16 |
| 839 | HSPA8 | P11142 | 0.77 | 0.113509275 | 0.01 | 17.74 | 17.75 |
| 840 | HSPA9 | P38646 | 0.52 | 0.283996656 | -0.03 | 17.58 | 17.55 |
| 841 | HSPB1 | P04792 | 0.22 | 0.657577319 | -0.05 | 17.28 | 17.23 |
| 842 | HSPB11 | Q9Y547 | 0.33 | 0.48148606 | -0.19 | 13.69 | 13.5 |
| 843 | HSPD1 | P10809 | 0.0024 | 2.619788758 | 0.09 | 16.14 | 16.23 |
| 844 | HSPE1 | P61604 | 0.6 | 0.22184875 | -0.02 | 18.39 | 18.37 |
| 845 | HSPG2 | P98160 | 0.28 | 0.552841969 | 0.04 | 15.46 | 15.5 |
| 846 | HSPH1 | Q92598 | 0.037 | 1.431798276 | 0.08 | 16.14 | 16.22 |

| | | | | | | | |
|---|---|---|---|---|---|---|---|
| 847 | HUWE1 | Q7Z6Z7 | 0.025 | 1.602059991 | 0.14 | 14.78 | 14.92 |
| 848 | HYOU1 | Q9Y4L1 | 0.2 | 0.698970004 | 0.04 | 15.91 | 15.95 |
| 849 | IARS | P41252 | 0.99 | 0.004364805 | 0 | 16.61 | 16.61 |
| 850 | IARS2 | Q9NSE4 | 0.32 | 0.494850022 | 0.04 | 16.83 | 16.87 |
| 851 | ICAM1 | P05362 | 0.0001 | 4 | -0.28 | 17.56 | 17.28 |
| 852 | ICAM2 | P13598 | 0.018 | 1.744727495 | -0.12 | 16.5 | 16.37 |
| 853 | IDE | P14735 | 0.18 | 0.744727495 | 0.1 | 15.45 | 15.54 |
| 854 | IDH1 | O75874 | 0.43 | 0.366531544 | 0.02 | 16.63 | 16.65 |
| 855 | IDH2 | P48735 | 0.93 | 0.031517051 | 0 | 16.78 | 16.78 |
| 856 | IDH3A | P50213 | 0.25 | 0.602059991 | 0.2 | 16.47 | 16.67 |
| 857 | IDH3B | O43837 | 0.0025 | 2.602059991 | 0.4 | 16.02 | 16.42 |
| 858 | IFI16 | Q16666 | 0.81 | 0.091514981 | -0.01 | 17.24 | 17.24 |
| 859 | IFI35 | P80217 | NA | NA | 0.12 | 16.05 | 16.17 |
| 860 | IFIT1 | P09914 | 0.076 | 1.119186408 | -0.42 | 16.83 | 16.41 |
| 861 | IGF2BP2 | Q9Y6M1 | 0.016 | 1.795880017 | -0.11 | 17.18 | 17.06 |
| 862 | IGF2BP3 | O00425 | 0.42 | 0.37675071 | -0.04 | 17.43 | 17.39 |
| 863 | IGF2R | P11717 | 0.72 | 0.142667504 | 0.01 | 17.15 | 17.16 |
| 864 | IGFBP7 | Q16270 | 0.21 | 0.677780705 | -0.24 | 16.76 | 16.52 |
| 865 | IKBIP | Q70UQ0 | 0.023 | 1.638272164 | 0.15 | 16.66 | 16.81 |
| 866 | ILF2 | Q12905 | 0.0015 | 2.823908741 | 0.28 | 15.44 | 15.72 |
| 867 | ILF3 | Q12906 | 0.00057 | 3.244125144 | 0.15 | 16.67 | 16.82 |
| 868 | ILK | Q13418 | 0.14 | 0.853871964 | 0.14 | 16.41 | 16.56 |
| 869 | ILKAP | Q9H0C8 | 0.51 | 0.292429824 | -0.19 | 17.19 | 17 |
| 870 | ILVBL | A1L0T0 | 0.39 | 0.408935393 | 0.07 | 14.64 | 14.71 |
| 871 | IMMT | Q16891 | 0.48 | 0.318758763 | 0.04 | 16.54 | 16.58 |
| 872 | IMPA1 | P29218 | 0.031 | 1.508638306 | 0.06 | 16.95 | 17.01 |
| 873 | IMPDH1 | P20839 | 0.77 | 0.113509275 | 0.03 | 14.68 | 14.71 |
| 874 | IMPDH2 | P12268 | 0.94 | 0.026872146 | 0.01 | 16.95 | 16.96 |
| 875 | INF2 | Q27J81 | 0.62 | 0.207608311 | 0.05 | 15.01 | 15.06 |
| 876 | INPP1 | P49441 | 0.08 | 1.096910013 | -0.07 | 15.59 | 15.52 |
| 877 | IPO4 | Q8TEX9 | 0.77 | 0.113509275 | 0.02 | 14.26 | 14.28 |
| 878 | IPO5 | O00410 | 0.073 | 1.13667714 | 0.06 | 15.97 | 16.03 |
| 879 | IPO7 | O95373 | 0.005 | 2.301029996 | 0.17 | 16.3 | 16.47 |
| 880 | IPO9 | Q96P70 | 0.7 | 0.15490196 | 0.02 | 15.29 | 15.31 |
| 881 | IQGAP1 | P46940 | 0.0001 | 4 | 0.14 | 15.82 | 15.96 |
| 882 | ISG15 | P05161 | 0.049 | 1.30980392 | 0.18 | 17.09 | 17.28 |
| 883 | ITGA2 | P17301 | 0.0001 | 4 | 0.23 | 16.24 | 16.47 |
| 884 | ITGA5 | P08648 | 0.0052 | 2.283996656 | 0.11 | 16.47 | 16.58 |
| 885 | ITGA6 | P23229 | 0.95 | 0.022276395 | 0.03 | 17.06 | 17.09 |
| 886 | ITGAV | P06756 | 0.00043 | 3.366531544 | 0.2 | 16.2 | 16.4 |
| 887 | ITGB1 | P05556 | 0.34 | 0.468521083 | 0.03 | 17.17 | 17.2 |
| 888 | ITGB3 | P05106 | 0.096 | 1.017728767 | 0.24 | 16.7 | 16.94 |
| 889 | ITPA | Q9BY32 | 0.19 | 0.721246399 | 0.07 | 15.3 | 15.37 |
| 890 | ITPR3 | Q14573 | 0.84 | 0.075720714 | 0.07 | 15.7 | 15.77 |
| 891 | ITPRID2 | P28290 | 0.15 | 0.823908741 | 0.15 | 17.26 | 17.41 |
| 892 | ITSN1 | Q15811 | 0.058 | 1.236572006 | -0.14 | 15.18 | 15.04 |
| 893 | IVD | P26440 | 0.34 | 0.468521083 | -0.13 | 18.64 | 18.51 |
| 894 | JAK1 | P23458 | 0.58 | 0.236572006 | 0.04 | 14.25 | 14.28 |
| 895 | JPT1 | Q9UK76 | 0.4 | 0.397940009 | -0.09 | 17.93 | 17.84 |
| 896 | JUP | P14923 | 0.0015 | 2.823908741 | 0.25 | 15.69 | 15.94 |

| | | | | | | | |
|---|---|---|---|---|---|---|---|
| 897 | KANK2 | Q63ZY3 | 0.026 | 1.585026652 | 0.62 | 13.99 | 14.61 |
| 898 | KANK3 | Q6NY19 | 0.58 | 0.236572006 | -0.04 | 15.73 | 15.69 |
| 899 | KARS | Q15046 | 0.034 | 1.468521083 | 0.12 | 16.34 | 16.46 |
| 900 | KCTD12 | Q96CX2 | 0.35 | 0.455931956 | 0.08 | 15.77 | 15.85 |
| 901 | KDR | P35968 | 0.25 | 0.602059991 | -0.1 | 15.77 | 15.67 |
| 902 | KHDRBS1 | Q07666 | 0.12 | 0.920818754 | -0.11 | 16.38 | 16.28 |
| 903 | KHSRP | Q92945 | 0.71 | 0.148741651 | -0.03 | 16.64 | 16.62 |
| 904 | KIF13B | Q9NQT8 | 0.09 | 1.045757491 | 0.59 | 14.23 | 14.82 |
| 905 | KIF5B | P33176 | 0.13 | 0.886056648 | -0.08 | 15.57 | 15.49 |
| 906 | KLC1 | Q07866 | 0.36 | 0.443697499 | -0.04 | 17.09 | 17.06 |
| 907 | KPNA1 | P52294 | 0.097 | 1.013228266 | 0.44 | 15.57 | 16.01 |
| 908 | KPNA2 | P52292 | 0.69 | 0.161150909 | -0.03 | 17.12 | 17.08 |
| 909 | KPNA3 | O00505 | 0.026 | 1.585026652 | 0.17 | 15.07 | 15.24 |
| 910 | KPNA4 | O00629 | 0.0029 | 2.537602002 | 0.34 | 14.14 | 14.48 |
| 911 | KPNA6 | O60684 | 0.4 | 0.397940009 | 0.22 | 13.27 | 13.49 |
| 912 | KPNB1 | Q14974 | 0.00019 | 3.721246399 | 0.18 | 15.99 | 16.17 |
| 913 | KRAS | P01116 | 0.45 | 0.346787486 | -0.14 | 17.11 | 16.97 |
| 914 | KRT1 | P04264 | 0.0049 | 2.30980392 | -0.14 | 17.3 | 17.16 |
| 915 | KRT10 | P13645 | 0.0049 | 2.30980392 | -0.12 | 16.2 | 16.08 |
| 916 | KRT18 | P05783 | 0.0001 | 4 0.21 | 17.31 | 17.52 | |
| 917 | KRT19 | P08727 | 0.048 | 1.318758763 | 0.12 | 16.32 | 16.44 |
| 918 | KRT2 | P35908 | 0.25 | 0.602059991 | -0.13 | 15.53 | 15.4 |
| 919 | KRT7 | P08729 | 0.0001 | 4 0.21 | 17.67 | 17.88 | |
| 920 | KRT9 | P35527 | 0.02 | 1.698970004 | -0.21 | 15.83 | 15.61 |
| 921 | KTN1 | Q86UP2 | 0.86 | 0.065501549 | 0 | 16.08 | 16.08 |
| 922 | KYAT3 | Q6YP21 | 0.063 | 1.200659451 | 0.33 | 17.02 | 17.34 |
| 923 | LACTB | P83111 | 0.11 | 0.958607315 | 0.38 | 15.82 | 16.21 |
| 924 | LAMA4 | Q16363 | 0.98 | 0.008773924 | 0.01 | 14.62 | 14.63 |
| 925 | LAMB1 | P07942 | 0.14 | 0.853871964 | -0.09 | 16.12 | 16.03 |
| 926 | LAMC1 | P11047 | 0.57 | 0.244125144 | 0.02 | 16.68 | 16.7 |
| 927 | LAMP1 | P11279 | 0.46 | 0.337242168 | -0.1 | 16.43 | 16.32 |
| 928 | LAMTOR1 | Q6IAA8 | 0.23 | 0.638272164 | 0.13 | 14.93 | 15.06 |
| 929 | LAMTOR5 | O43504 | 0.52 | 0.283996656 | -0.05 | 16.18 | 16.12 |
| 930 | LAP3 | P28838 | 0.0001 | 4 -0.23 | 16.57 | 16.34 | |
| 931 | LARP1 | Q6PKG0 | 0.41 | 0.387216143 | 0.1 | 16.29 | 16.39 |
| 932 | LARP4B | Q92615 | 0.79 | 0.102372909 | -0.04 | 14.82 | 14.77 |
| 933 | LARS | Q9P2J5 | 0.0086 | 2.065501549 | 0.09 | 15.57 | 15.66 |
| 934 | LASP1 | Q14847 | 0.011 | 1.958607315 | -0.11 | 18.67 | 18.56 |
| 935 | LDHA | P00338 | 0.003 | 2.522878745 | 0.1 | 16.52 | 16.61 |
| 936 | LDHB | P07195 | 0.18 | 0.744727495 | 0.04 | 16.73 | 16.77 |
| 937 | LETM1 | O95202 | 0.51 | 0.292429824 | 0.07 | 16.17 | 16.24 |
| 938 | LGALS1 | P09382 | 0.046 | 1.337242168 | -0.19 | 17.52 | 17.33 |
| 939 | LGALS3 | P17931 | 0.8 | 0.096910013 | 0.03 | 18.05 | 18.08 |
| 940 | LIMA1 | Q9UHB6 | 0.032 | 1.494850022 | 0.24 | 17.37 | 17.61 |
| 941 | LIMS1 | P48059 | 0.045 | 1.346787486 | 0.17 | 16.14 | 16.31 |
| 942 | LMAN1 | P49257 | 0.0032 | 2.494850022 | -0.11 | 17.17 | 17.07 |
| 943 | LMAN2 | Q12907 | 0.79 | 0.102372909 | -0.01 | 17.88 | 17.87 |
| 944 | LMNA | P02545 | 0.0035 | 2.455931956 | -0.1 | 17.69 | 17.59 |
| 945 | LMNB1 | P20700 | 0.0001 | 4 -0.19 | 16.95 | 16.76 | |
| 946 | LMNB2 | Q03252 | 0.061 | 1.214670165 | -0.08 | 17.21 | 17.13 |

| | | | | | | | | |
|---|---|---|---|---|---|---|---|---|
| 947 | LNPEP | Q9UIQ6 | 0.58 | 0.236572006 | -0.03 | 14.97 | 14.94 | |
| 948 | LONP1 | P36776 | 0.022 | 1.657577319 | -0.13 | 15.16 | 15.03 | |
| 949 | LPCAT2 | Q7L5N7 | 0.045 | 1.346787486 | 0.34 | 14.4 | 14.74 | |
| 950 | LPP | Q93052 | 0.062 | 1.207608311 | -0.1 | 16.41 | 16.31 | |
| 951 | LPXN | O60711 | 0.027 | 1.568636236 | 0.29 | 16.19 | 16.48 | |
| 952 | LRP1 | Q07954 | NA | NA | -0.38 | 20.1 | 19.72 | |
| 953 | LRPPRC | P42704 | 0.35 | 0.455931956 | 0.03 | 15.08 | 15.11 | |
| 954 | LRRC40 | Q9H9A6 | 0.34 | 0.468521083 | -0.17 | 15.15 | 14.98 | |
| 955 | LRRC47 | Q8N1G4 | 0.97 | 0.013228266 | 0 | 16.97 | 16.97 | |
| 956 | LRRC57 | Q8N9N7 | 0.24 | 0.619788758 | 0.75 | 15.7 | 16.45 | |
| 957 | LRRC59 | Q96AG4 | 0.0012 | 2.920818754 | -0.14 | 16.65 | 16.51 | |
| 958 | LRRFIP1 | Q32MZ4 | 0.13 | 0.886056648 | 0.09 | 14.46 | 14.54 | |
| 959 | LSM2 | Q9Y333 | 0.68 | 0.167491087 | -0.12 | 14.74 | 14.62 | |
| 960 | LSM4 | Q9Y4Z0 | 0.35 | 0.455931956 | -0.21 | 16.19 | 15.98 | |
| 961 | LSM8 | O95777 | 0.39 | 0.408935393 | -0.09 | 14.75 | 14.66 | |
| 962 | LSS | P48449 | 0.069 | 1.161150909 | 0.23 | 16.73 | 16.96 | |
| 963 | LTA4H | P09960 | 0.99 | 0.004364805 | 0 | 17.36 | 17.36 | |
| 964 | LTBP2 | Q14767 | 0.41 | 0.387216143 | 0.14 | 16.62 | 16.76 | |
| 965 | LTV1 | Q96GA3 | 0.43 | 0.366531544 | 0.26 | 14.35 | 14.6 | |
| 966 | LUC7L2 | Q9Y383 | 0.42 | 0.37675071 | 0.07 | 16.83 | 16.9 | |
| 967 | LUC7L3 | O95232 | 0.34 | 0.468521083 | -0.13 | 15.92 | 15.79 | |
| 968 | LXN | Q9BS40 | 0.03 | 1.522878745 | 0.09 | 17.11 | 17.2 | |
| 969 | LYAR | Q9NX58 | 0.25 | 0.602059991 | -0.15 | 15.17 | 15.02 | |
| 970 | LYPLA1 | O75608 | 0.057 | 1.244125144 | 0.27 | 15.6 | 15.88 | |
| 971 | LYPLA2 | O95372 | 0.063 | 1.200659451 | 0.17 | 15.78 | 15.95 | |
| 972 | MACF1 | Q9UPN3 | 0.16 | 0.795880017 | 0.08 | 14.78 | 14.86 | |
| 973 | MAGED2 | Q9UNF1 | 0.0056 | 2.251811973 | -0.12 | 17.58 | 17.46 | |
| 974 | MAGOH | P61326 | (+1) | 0.13 | 0.886056648 | 0.12 | 15.69 | 15.8 |
| 975 | MAN2A1 | Q16706 | 0.0037 | 2.431798276 | 0.31 | 15.59 | 15.9 | |
| 976 | MANF | P55145 | 0.99 | 0.004364805 | 0 | 16.51 | 16.52 | |
| 977 | MAOA | P21397 | 0.29 | 0.537602002 | -0.13 | 16.25 | 16.12 | |
| 978 | MAP1B | P46821 | 0.0001 | 4 | 0.15 | 16.89 | 17.04 | |
| 979 | MAP1S | Q66K74 | 0.0031 | 2.508638306 | 0.17 | 15.05 | 15.22 | |
| 980 | MAP2K1 | Q02750 | 0.0073 | 2.13667714 | 0.17 | 15.89 | 16.06 | |
| 981 | MAP4 | P27816 | 0.52 | 0.283996656 | 0.02 | 17.3 | 17.32 | |
| 982 | MAP7D1 | Q3KQU3 | 0.0095 | 2.022276395 | -0.12 | 15.18 | 15.06 | |
| 983 | MAPK1 | P28482 | 0.037 | 1.431798276 | 0.12 | 15.57 | 15.69 | |
| 984 | MAPK12 | P53778 | 0.45 | 0.346787486 | 0.2 | 14.62 | 14.82 | |
| 985 | MAPRE1 | Q15691 | 0.71 | 0.148741651 | -0.04 | 15.67 | 15.63 | |
| 986 | MAPRE2 | Q15555 | 0.35 | 0.455931956 | -0.07 | 16.78 | 16.71 | |
| 987 | MARCKS | P29966 | 0.0001 | 4 | -0.24 | 16.94 | 16.69 | |
| 988 | MARCKSL1 | | P49006 | 0.42 | 0.37675071 | -0.06 | 16.01 | 15.95 |
| 989 | MARS | P56192 | 0.052 | 1.283996656 | 0.1 | 15.39 | 15.49 | |
| 990 | MAT2A | P31153 | 0.033 | 1.48148606 | -0.08 | 18.22 | 18.14 | |
| 991 | MAT2B | Q9NZL9 | 0.014 | 1.853871964 | 0.24 | 16.29 | 16.53 | |
| 992 | MATR3 | P43243 | 0.41 | 0.387216143 | -0.04 | 15.31 | 15.27 | |
| 993 | MAVS | Q7Z434 | 0.17 | 0.769551079 | 0.07 | 16.25 | 16.32 | |
| 994 | MBNL1 | Q9NR56 | 0.096 | 1.017728767 | -0.07 | 17.19 | 17.12 | |
| 995 | MBOAT7 | Q96N66 | 0.85 | 0.070581074 | -0.02 | 17.48 | 17.46 | |
| 996 | MCAM | P43121 | 0.96 | 0.017728767 | 0 | 16.5 | 16.5 | |

| | | | | | | | |
|---|---|---|---|---|---|---|---|
| 997 | MCCC2 | Q9HCC0 | 0.063 | 1.200659451 | 0.11 | 16.3 | 16.41 |
| 998 | MCFD2 | Q8NI22 | 0.072 | 1.142667504 | -0.21 | 15.45 | 15.24 |
| 999 | MCM2 | P49736 | 0.092 | 1.036212173 | -0.17 | 14.28 | 14.11 |
| 1000 | MCM3 | P25205 | 0.71 | 0.148741651 | -0.04 | 16.06 | 16.02 |
| 1001 | MCMBP | Q9BTE3 | 0.15 | 0.823908741 | -0.12 | 15.9 | 15.78 |
| 1002 | MCTS1 | Q9ULC4 | 0.25 | 0.602059991 | 0.14 | 16.02 | 16.16 |
| 1003 | MDH1 | P40925 | 0.94 | 0.026872146 | 0 | 17.76 | 17.76 |
| 1004 | MDH2 | P40926 | 0.0041 | 2.387216143 | 0.13 | 17.19 | 17.32 |
| 1005 | MDN1 | Q9NU22 | 0.73 | 0.13667714 | 0.08 | 18.84 | 18.92 |
| 1006 | ME1 | P48163 | 0.12 | 0.920818754 | -0.15 | 16.06 | 15.91 |
| 1007 | ME2 | P23368 | 0.2 | 0.698970004 | 0.17 | 14.59 | 14.75 |
| 1008 | ME3 | Q16798 | 0.045 | 1.346787486 | 0.17 | 14.57 | 14.75 |
| 1009 | MECP2 | P51608 | 0.26 | 0.585026652 | 0.15 | 16.48 | 16.63 |
| 1010 | MEMO1 | Q9Y316 | 0.083 | 1.080921908 | 0.22 | 14.15 | 14.37 |
| 1011 | MESD | Q14696 | 0.026 | 1.585026652 | -0.24 | 18.01 | 17.78 |
| 1012 | METAP1 | P53582 | 0.24 | 0.619788758 | 0.21 | 14.48 | 14.68 |
| 1013 | MFN1 | Q8IWA4 | 0.72 | 0.142667504 | -0.07 | 14.78 | 14.71 |
| 1014 | MFN2 | O95140 | 0.39 | 0.408935393 | -0.11 | 15.06 | 14.96 |
| 1015 | MGARP | Q8TDB4 | 0.66 | 0.180456064 | 0.06 | 15.76 | 15.82 |
| 1016 | MGST3 | O14880 | 0.74 | 0.13076828 | -0.05 | 18.71 | 18.67 |
| 1017 | MIA3 | Q5JRA6 | 0.49 | 0.30980392 | 0.36 | 14.01 | 14.37 |
| 1018 | MICALL2 | Q8IY33 | 0.024 | 1.619788758 | 0.28 | 16.5 | 16.78 |
| 1019 | MICOS13 | Q5XKP0 | 0.097 | 1.013228266 | 0.23 | 13.31 | 13.54 |
| 1020 | MIF | P14174 | 0.41 | 0.387216143 | 0.24 | 17.13 | 17.37 |
| 1021 | MLEC | Q14165 | 0.37 | 0.431798276 | 0.05 | 16.57 | 16.62 |
| 1022 | MMP1 | P03956 | 0.96 | 0.017728767 | -0.02 | 16.16 | 16.14 |
| 1023 | MMRN1 | Q13201 | 0.23 | 0.638272164 | -0.07 | 15.89 | 15.82 |
| 1024 | MMRN2 | Q9H8L6 | 0.16 | 0.795880017 | 0.13 | 15.06 | 15.19 |
| 1025 | MMS19 | Q96T76 | 0.15 | 0.823908741 | 0.14 | 14.29 | 14.43 |
| 1026 | MOGS | Q13724 | 0.0079 | 2.102372909 | 0.41 | 14.96 | 15.37 |
| 1027 | MON2 | Q7Z3U7 | 0.077 | 1.113509275 | 0.27 | 14.71 | 14.98 |
| 1028 | MORC2 | Q9Y6X9 | NA | NA | 0.24 | 16.62 | 16.86 |
| 1029 | MPDU1 | O75352 | 0.76 | 0.119186408 | 0.02 | 16.16 | 16.18 |
| 1030 | MPHOSPH10 | O00566 | 0.045 | 1.346787486 | 0.25 | 15.06 | 15.31 |
| 1031 | MPI | P34949 | 0.89 | 0.050609993 | 0 | 15.2 | 15.2 |
| 1032 | MPST | P25325 | 0.88 | 0.055517328 | -0.01 | 15.84 | 15.82 |
| 1033 | MRE11 | P49959 | 0.9 | 0.045757491 | 0.02 | 16.41 | 16.44 |
| 1034 | MRPL1 | Q9BYD6 | 0.71 | 0.148741651 | -0.02 | 14.34 | 14.32 |
| 1035 | MRPL12 | P52815 | 0.43 | 0.366531544 | -0.11 | 14.12 | 14.01 |
| 1036 | MRPL3 | P09001 | 0.097 | 1.013228266 | 0.2 | 17.85 | 18.05 |
| 1037 | MRPL39 | Q9NYK5 | 0.39 | 0.408935393 | 0.08 | 14.69 | 14.77 |
| 1038 | MRPL4 | Q9BYD3 | 0.75 | 0.124938737 | -0.08 | 15.21 | 15.13 |
| 1039 | MRPL44 | Q9H9J2 | 0.89 | 0.050609993 | 0.02 | 14.5 | 14.52 |
| 1040 | MRPL9 | Q9BYD2 | 0.98 | 0.008773924 | 0.05 | 15.73 | 15.78 |
| 1041 | MRPS22 | P82650 | 0.32 | 0.494850022 | 0.12 | 16.96 | 17.08 |
| 1042 | MRPS27 | Q92552 | 0.68 | 0.167491087 | -0.04 | 16.05 | 16.01 |
| 1043 | MRTO4 | Q9UKD2 | 0.41 | 0.387216143 | 0.07 | 16.52 | 16.59 |
| 1044 | MSH2 | P43246 | 0.042 | 1.37675071 | 0.2 | 15.36 | 15.56 |
| 1045 | MSN | P26038 | 0.0001 | 4 | -0.19 | 16.7 | 16.51 |
| 1046 | MT1E | P04732 | 0.091 | 1.040958608 | -0.27 | 15.61 | 15.34 |

| | | | | | | | |
|---|---|---|---|---|---|---|---|
| 1047 | MT2A | P02795 | 0.054 | 1.26760624 | -0.67 | 15.7 | 15.03 |
| 1048 | MTA2 | O94776 | 0.2 | 0.698970004 | -0.26 | 17.53 | 17.26 |
| 1049 | MTAP | Q13126 | 0.12 | 0.920818754 | 0.09 | 14.07 | 14.16 |
| 1050 | MTCH2 | Q9Y6C9 | 0.11 | 0.958607315 | 0.2 | 16.98 | 17.18 |
| 1051 | MT-CO2 | P00403 | 0.4 | 0.397940009 | 0.03 | 14.97 | 15 |
| 1052 | MTDH | Q86UE4 | 0.0026 | 2.585026652 | -0.15 | 17.52 | 17.37 |
| 1053 | MTHFD1 | P11586 | 0.59 | 0.229147988 | 0.01 | 16.46 | 16.47 |
| 1054 | MTHFD1L | Q6UB35 | 0.43 | 0.366531544 | 0.04 | 15.05 | 15.09 |
| 1055 | MTMR2 | Q13614 | 0.52 | 0.283996656 | 0.07 | 16.23 | 16.3 |
| 1056 | MT-ND1 | P03886 | 0.37 | 0.431798276 | 0.13 | 14.71 | 14.84 |
| 1057 | MTOR | P42345 | 0.041 | 1.387216143 | 0.4 | 14.39 | 14.79 |
| 1058 | MTPN | P58546 | 0.0078 | 2.107905397 | 0.32 | 15.21 | 15.53 |
| 1059 | MTREX | P42285 | 0.73 | 0.13667714 | -0.03 | 13.15 | 13.12 |
| 1060 | MTX2 | O75431 | 0.18 | 0.744727495 | -0.22 | 14.58 | 14.37 |
| 1061 | MVP | Q14764 | 0.0077 | 2.113509275 | 0.08 | 16.12 | 16.2 |
| 1062 | MX1 | P20591 | 0.00012 | 3.920818754 | -0.47 | 16.57 | 16.1 |
| 1063 | MYADM | Q96S97 | 0.065 | 1.187086643 | 0.24 | 16.51 | 16.75 |
| 1064 | MYBBP1A | Q9BQG0 | 0.0024 | 2.619788758 | 0.24 | 15.39 | 15.63 |
| 1065 | MYDGF | Q969H8 | 0.19 | 0.721246399 | 0.15 | 17.45 | 17.6 |
| 1066 | MYH10 | P35580 | 0.056 | 1.251811973 | -0.1 | 14.61 | 14.51 |
| 1067 | MYH16 | Q9H6N6 | 0.052 | 1.283996656 | -0.32 | 19.35 | 19.04 |
| 1068 | MYH9 | P35579 | 0.12 | 0.920818754 | 0.03 | 15.97 | 16.01 |
| 1069 | MYL12B | O14950 | 0.00043 | 3.366531544 | 0.17 | 15.35 | 15.52 |
| 1070 | MYL6 | P60660 | 0.18 | 0.744727495 | 0.06 | 16.99 | 17.05 |
| 1071 | MYO18A | Q92614 | 0.013 | 1.886056648 | 0.59 | 12.62 | 13.21 |
| 1072 | MYO1B | O43795 | 0.45 | 0.346787486 | 0.21 | 15.96 | 16.17 |
| 1073 | MYO1C | O00159 | 0.15 | 0.823908741 | 0.05 | 16.12 | 16.17 |
| 1074 | MYO1D | O94832 | 0.39 | 0.408935393 | 0.07 | 15.47 | 15.54 |
| 1075 | MYO1E | Q12965 | 0.02 | 1.698970004 | 0.31 | 15.2 | 15.51 |
| 1076 | MYO5A | Q9Y4I1 | 0.18 | 0.744727495 | 0.12 | 15.1 | 15.22 |
| 1077 | MYO6 | Q9UM54 | 0.0038 | 2.420216403 | 0.2 | 14.82 | 15.02 |
| 1078 | MYO9B | Q13459 | 0.14 | 0.853871964 | 0.17 | 15.1 | 15.27 |
| 1079 | MYOF | Q9NZM1 | 0.0001 | 4 | 0.13 | 16.33 | 16.46 |
| 1080 | NAA10 | P41227 | 0.71 | 0.148741651 | 0.04 | 17.95 | 17.99 |
| 1081 | NAA15 | Q9BXJ9 | 0.032 | 1.494850022 | 0.16 | 16.03 | 16.2 |
| 1082 | NACA | E9PAV3 | 0.01 | 2 | -0.16 | 17.37 | 17.21 |
| 1083 | NAGK | Q9UJ70 | 0.28 | 0.552841969 | -0.1 | 15.51 | 15.41 |
| 1084 | NAMPT | P43490 | 0.87 | 0.060480747 | -0.01 | 16.16 | 16.15 |
| 1085 | NANS | Q9NR45 | 0.058 | 1.236572006 | -0.08 | 15.09 | 15.02 |
| 1086 | NAP1L1 | P55209 | 0.64 | 0.193820026 | -0.02 | 15.59 | 15.57 |
| 1087 | NAP1L4 | Q99733 | 0.14 | 0.853871964 | -0.12 | 15.79 | 15.67 |
| 1088 | NAPA | P54920 | 0.86 | 0.065501549 | -0.03 | 15.7 | 15.67 |
| 1089 | NARS | O43776 | 0.023 | 1.638272164 | -0.08 | 16.22 | 16.14 |
| 1090 | NASP | P49321 | 0.2 | 0.698970004 | -0.04 | 16.33 | 16.3 |
| 1091 | NAT10 | Q9H0A0 | 0.42 | 0.37675071 | -0.06 | 15.06 | 15 |
| 1092 | NAXD | Q8IW45 | 0.73 | 0.13667714 | -0.03 | 16.19 | 16.16 |
| 1093 | NCBP1 | Q09161 | 0.32 | 0.494850022 | 0.25 | 14.44 | 14.68 |
| 1094 | NCEH1 | Q6PIU2 | 1 | 0 | -0.01 | 15.44 | 15.44 |
| 1095 | NCK1 | P16333 | 0.79 | 0.102372909 | -0.04 | 15.84 | 15.79 |
| 1096 | NCKAP1 | Q9Y2A7 | 0.78 | 0.107905397 | -0.01 | 15.6 | 15.6 |

| | | | | | | | |
|---|---|---|---|---|---|---|---|
| 1097 | NCL | P19338 | 0.0001 | 4 | 0.14 | 17.36 | 17.5 |
| 1098 | NDRG1 | Q92597 | 0.053 | 1.27572413 | 0.41 | 16.09 | 16.51 |
| 1099 | NDRG3 | Q9UGV2 | 0.031 | 1.508638306 | 0.35 | 16.98 | 17.33 |
| 1100 | NDUFA10 | O95299 | 0.22 | 0.657577319 | -0.08 | 14.64 | 14.56 |
| 1101 | NDUFA4 | O00483 | 0.39 | 0.408935393 | -0.17 | 19.43 | 19.26 |
| 1102 | NDUFA5 | Q16718 | 0.68 | 0.167491087 | -0.02 | 14.72 | 14.7 |
| 1103 | NDUFA7 | O95182 | 0.71 | 0.148741651 | -0.02 | 17.82 | 17.8 |
| 1104 | NDUFA9 | Q16795 | 0.061 | 1.214670165 | 0.42 | 15.82 | 16.24 |
| 1105 | NDUFS1 | P28331 | 0.002 | 2.698970004 | 0.17 | 15.79 | 15.96 |
| 1106 | NDUFS2 | O75306 | 0.28 | 0.552841969 | 0.08 | 16.84 | 16.92 |
| 1107 | NDUFS4 | O43181 | 0.13 | 0.886056648 | -0.22 | 17.07 | 16.85 |
| 1108 | NDUFV1 | P49821 | 0.22 | 0.657577319 | 0.16 | 15.1 | 15.26 |
| 1109 | NDUFV2 | P19404 | 0.79 | 0.102372909 | -0.01 | 15.35 | 15.34 |
| 1110 | NEB | P20929 | 0.47 | 0.327902142 | -0.05 | 17.37 | 17.32 |
| 1111 | NECAP2 | Q9NVZ3 | 0.94 | 0.026872146 | 0.01 | 16.68 | 16.69 |
| 1112 | NECTIN2 | Q92692 | 0.23 | 0.638272164 | 0.14 | 18.66 | 18.8 |
| 1113 | NEDD4 | P46934 | 0.016 | 1.795880017 | 0.33 | 15.69 | 16.02 |
| 1114 | NEMF | O60524 | 0.79 | 0.102372909 | 0 | 16.25 | 16.25 |
| 1115 | NES | P48681 | 0.0001 | 4 | 0.19 | 16.55 | 16.74 |
| 1116 | NFKB1 | P19838 | 0.13 | 0.886056648 | 0.1 | 15.68 | 15.78 |
| 1117 | NFKB2 | Q00653 | 0.098 | 1.008773924 | 0.16 | 13.69 | 13.84 |
| 1118 | NFU1 | Q9UMS0 | 0.14 | 0.853871964 | 0.21 | 16.09 | 16.3 |
| 1119 | NIBAN2 | Q96TA1 | 0.0033 | 2.48148606 | 0.13 | 15.93 | 16.06 |
| 1120 | NIFK | Q9BYG3 | 0.14 | 0.853871964 | 0.12 | 15.6 | 15.72 |
| 1121 | NLN | Q9BYT8 | 0.059 | 1.229147988 | 0.37 | 14.11 | 14.48 |
| 1122 | NMD3 | Q96D46 | 0.35 | 0.455931956 | 0.06 | 15.04 | 15.1 |
| 1123 | NME1 | P15531 | 0.27 | 0.568636236 | 0.07 | 17.19 | 17.26 |
| 1124 | NME2 | P22392 | 0.38 | 0.420216403 | 0.06 | 16.51 | 16.57 |
| 1125 | NMT1 | P30419 | 0.065 | 1.187086643 | -0.12 | 15.71 | 15.59 |
| 1126 | NNMT | P40261 | 0.17 | 0.769551079 | -0.17 | 16.58 | 16.41 |
| 1127 | NNT | Q13423 | 0.0023 | 2.638272164 | 0.16 | 16.51 | 16.67 |
| 1128 | NOLC1 | Q14978 | 0.084 | 1.075720714 | 0.08 | 17.75 | 17.83 |
| 1129 | NOMO1 | Q15155 (+1) | 0.038 | 1.420216403 | 0.11 | 16.1 | 16.21 |
| 1130 | NONO | Q15233 | 0.16 | 0.795880017 | 0.09 | 16.81 | 16.9 |
| 1131 | NOP53 | Q9NZM5 | 0.62 | 0.207608311 | 0.03 | 13.39 | 13.41 |
| 1132 | NOP56 | O00567 | 0.0081 | 2.091514981 | 0.1 | 16.06 | 16.16 |
| 1133 | NOP58 | Q9Y2X3 | 0.56 | 0.251811973 | 0.03 | 15.46 | 15.49 |
| 1134 | NOVA2 | Q9UNW9 | 0.0025 | 2.602059991 | 0.26 | 16.14 | 16.4 |
| 1135 | NPC2 | P61916 | 0.64 | 0.193820026 | -0.05 | 16.07 | 16.02 |
| 1136 | NPEPPS | P55786 | 0.28 | 0.552841969 | 0.05 | 16.01 | 16.05 |
| 1137 | NPLOC4 | Q8TAT6 | 0.64 | 0.193820026 | -0.03 | 15.73 | 15.7 |
| 1138 | NPM1 | P06748 | 0.37 | 0.431798276 | 0.04 | 16.87 | 16.9 |
| 1139 | NPM3 | O75607 | 0.05 | 1.301029996 | 0.24 | 15.02 | 15.26 |
| 1140 | NQO1 | P15559 | 0.00053 | 3.27572413 | -0.22 | 17.5 | 17.29 |
| 1141 | NRBP1 | Q9UHY1 | 0.081 | 1.091514981 | -0.11 | 15.41 | 15.3 |
| 1142 | NRCAM | Q92823 | 0.42 | 0.37675071 | 0.04 | 16.29 | 16.34 |
| 1143 | NRDC | O43847 | 0.62 | 0.207608311 | -0.05 | 15.39 | 15.34 |
| 1144 | NSA2 | O95478 | 0.11 | 0.958607315 | 0.23 | 16.82 | 17.05 |
| 1145 | NSDHL | Q15738 | 0.2 | 0.698970004 | 0.16 | 17.22 | 17.39 |
| 1146 | NSF | P46459 | 0.71 | 0.148741651 | 0.01 | 16.33 | 16.34 |

| | | | | | | | |
|---|---|---|---|---|---|---|---|
| 1147 | NSFL1C | Q9UNZ2 | 0.003 | 2.522878745 | -0.16 | 16.7 | 16.53 |
| 1148 | NSUN2 | Q08J23 | 0.43 | 0.366531544 | 0.08 | 15.73 | 15.81 |
| 1149 | NT5E | P21589 | 0.0001 | 4 | -0.14 | 16.25 | 16.11 |
| 1150 | NTPCR | Q9BSD7 | 0.022 | 1.657577319 | 0.32 | 16.55 | 16.86 |
| 1151 | NUCB1 | Q02818 | 0.023 | 1.638272164 | -0.12 | 15.93 | 15.81 |
| 1152 | NUCKS1 | Q9H1E3 | 0.01 | 2 | -0.35 | 18.59 | 18.24 |
| 1153 | NUDC | Q9Y266 | 0.42 | 0.37675071 | -0.05 | 18.7 | 18.65 |
| 1154 | NUDCD1 | Q96RS6 | 0.054 | 1.26760624 | 0.38 | 15.01 | 15.39 |
| 1155 | NUDT21 | O43809 | 0.0068 | 2.167491087 | 0.34 | 15.94 | 16.28 |
| 1156 | NUDT4B | A0A024RBG1 (+1) | 0.91 | 0.040958608 | -0.01 | 17.3 | 17.29 |
| 1157 | NUDT5 | Q9UKK9 | 0.34 | 0.468521083 | 0.11 | 16.09 | 16.2 |
| 1158 | NUDT9 | Q9BW91 | 0.082 | 1.086186148 | 0.35 | 15.46 | 15.82 |
| 1159 | NUFIP2 | Q7Z417 | 0.0046 | 2.337242168 | 0.41 | 16.92 | 17.33 |
| 1160 | NUMA1 | Q14980 | 0.88 | 0.055517328 | -0.01 | 15.29 | 15.29 |
| 1161 | NUP133 | Q8WUM0 | 0.47 | 0.327902142 | 0.03 | 16.06 | 16.1 |
| 1162 | NUP153 | P49790 | 0.16 | 0.795880017 | 0.11 | 15.57 | 15.68 |
| 1163 | NUP155 | O75694 | 0.59 | 0.229147988 | 0.06 | 14.65 | 14.71 |
| 1164 | NUP160 | Q12769 | 0.041 | 1.387216143 | 0.27 | 14.23 | 14.5 |
| 1165 | NUP205 | Q92621 | 0.011 | 1.958607315 | 0.48 | 14.27 | 14.75 |
| 1166 | NUP214 | P35658 | 0.94 | 0.026872146 | 0.05 | 14.96 | 15.01 |
| 1167 | NUP37 | Q8NFH4 | 0.18 | 0.744727495 | 0.12 | 14.52 | 14.64 |
| 1168 | NUP50 | Q9UKX7 | 0.00061 | 3.214670165 | 0.27 | 14.05 | 14.32 |
| 1169 | NUP54 | Q7Z3B4 | 0.21 | 0.677780705 | 0.14 | 16.05 | 16.19 |
| 1170 | NUP62 | P37198 | 0.73 | 0.13667714 | 0.03 | 15.63 | 15.66 |
| 1171 | NUP93 | Q8N1F7 | 0.42 | 0.37675071 | 0.09 | 17.52 | 17.61 |
| 1172 | NUP98 | P52948 | 0.63 | 0.200659451 | -0.07 | 16.43 | 16.36 |
| 1173 | NUTF2 | P61970 | 0.14 | 0.853871964 | 0.23 | 17.41 | 17.64 |
| 1174 | NVL | O15381 | 0.82 | 0.086186148 | 0.06 | 14.65 | 14.71 |
| 1175 | NXN | Q6DKJ4 | 0.63 | 0.200659451 | -0.05 | 14.48 | 14.43 |
| 1176 | OAS2 | P29728 | 0.063 | 1.200659451 | -0.26 | 17.38 | 17.11 |
| 1177 | OAS3 | Q9Y6K5 | 0.042 | 1.37675071 | -0.45 | 15.67 | 15.22 |
| 1178 | OAT | P04181 | 0.094 | 1.026872146 | 0.19 | 14.1 | 14.29 |
| 1179 | OCIAD1 | Q9NX40 | 0.49 | 0.30980392 | 0.05 | 15.79 | 15.84 |
| 1180 | OCIAD2 | Q56VL3 | 0.0007 | 3.15490196 | 0.56 | 16.06 | 16.63 |
| 1181 | OGA | O60502 | 0.025 | 1.602059991 | 0.15 | 14.8 | 14.95 |
| 1182 | OGDH | Q02218 | 0.04 | 1.397940009 | 0.13 | 15.75 | 15.87 |
| 1183 | OGFR | Q9NZT2 | 0.17 | 0.769551079 | -0.16 | 16.95 | 16.8 |
| 1184 | OLA1 | Q9NTK5 | 0.75 | 0.124938737 | 0.02 | 16.2 | 16.22 |
| 1185 | OPA1 | O60313 | 0.087 | 1.060480747 | 0.11 | 15.72 | 15.82 |
| 1186 | OPTN | Q96CV9 | 0.76 | 0.119186408 | -0.01 | 16.28 | 16.27 |
| 1187 | OSBP | P22059 | 0.58 | 0.236572006 | 0.05 | 15.16 | 15.22 |
| 1188 | OSTF1 | Q92882 | 0.04 | 1.397940009 | -0.4 | 15.77 | 15.37 |
| 1189 | OTUB1 | Q96FW1 | 0.09 | 1.045757491 | -0.11 | 15.41 | 15.3 |
| 1190 | OTUD6B | Q8N6M0 | NA | NA | 0 | NA | NA |
| 1191 | OXCT1 | P55809 | 0.19 | 0.721246399 | 0.07 | 15.22 | 15.29 |
| 1192 | OXSR1 | O95747 | 0.61 | 0.214670165 | -0.15 | 15.2 | 15.05 |
| 1193 | P3H1 | Q32P28 | 0.51 | 0.292429824 | 0.08 | 15.71 | 15.79 |
| 1194 | P3H3 | Q8IVL6 | 0.46 | 0.337242168 | 0.05 | 15.31 | 15.37 |
| 1195 | P3H4 | Q92791 | 0.82 | 0.086186148 | -0.05 | 15.45 | 15.4 |
| 1196 | P4HA1 | P13674 | 0.023 | 1.638272164 | -0.19 | 17.26 | 17.07 |

| | | | | | | | | | |
|---|---|---|---|---|---|---|---|---|---|
| 1197 | P4HA2 | O15460 | 0.13 | 0.886056648 | | -0.09 | 16.38 | 16.29 | |
| 1198 | P4HB | P07237 | 0.00036 | 3.443697499 | | -0.1 | 17.48 | 17.39 | |
| 1199 | PA2G4 | Q9UQ80 | 0.026 | 1.585026652 | | 0.08 | 16.89 | 16.97 | |
| 1200 | PABPC1 | P11940 | 0.34 | 0.468521083 | | -0.04 | 17.85 | 17.81 | |
| 1201 | PABPC4 | Q13310 | 0.0001 | 4 | 0.42 | 17.18 | 17.6 | | |
| 1202 | PABPN1 | Q86U42 | 0.03 | 1.522878745 | | -0.24 | 15.25 | 15.01 | |
| 1203 | PACS1 | Q6VY07 | 0.0025 | 2.602059991 | | 0.16 | 15.9 | 16.06 | |
| 1204 | PACSIN2 | Q9UNF0 | 0.092 | 1.036212173 | | -0.07 | 15.46 | 15.38 | |
| 1205 | PAFAH1B1 | | P43034 | 0.68 | 0.167491087 | | -0.04 | 18.45 | 18.41 |
| 1206 | PAFAH1B2 | | P68402 | 0.57 | 0.244125144 | | 0.32 | 14.54 | 14.86 |
| 1207 | PAFAH1B3 | | Q15102 | 0.61 | 0.214670165 | | 0.02 | 15.48 | 15.5 |
| 1208 | PAICS | P22234 | 0.82 | 0.086186148 | | -0.01 | 16.74 | 16.73 | |
| 1209 | PAK2 | Q13177 | 0.034 | 1.468521083 | | 0.11 | 16.82 | 16.93 | |
| 1210 | PALLD | Q8WX93 | 0.0059 | 2.229147988 | | 0.75 | 15.13 | 15.87 | |
| 1211 | PALMD | Q9NP74 | 0.012 | 1.920818754 | | -0.32 | 16.68 | 16.35 | |
| 1212 | PAPSS1 | O43252 | 0.22 | 0.657577319 | | 0.44 | 13.94 | 14.38 | |
| 1213 | PAPSS2 | O95340 | 0.052 | 1.283996656 | | 0.11 | 15.96 | 16.08 | |
| 1214 | PARK7 | Q99497 | 0.057 | 1.244125144 | | -0.09 | 17.42 | 17.33 | |
| 1215 | PARP1 | P09874 | 0.79 | 0.102372909 | | 0.02 | 17.08 | 17.1 | |
| 1216 | PARP14 | Q460N5 | 0.47 | 0.327902142 | | 0.04 | 14.6 | 14.64 | |
| 1217 | PARP4 | Q9UKK3 | 0.0095 | 2.022276395 | | 0.22 | 15.09 | 15.31 | |
| 1218 | PARP9 | Q8IXQ6 | 0.44 | 0.356547324 | | 0.07 | 15.14 | 15.21 | |
| 1219 | PARVA | Q9NVD7 | 0.051 | 1.292429824 | | 0.13 | 16.99 | 17.12 | |
| 1220 | PARVB | Q9HBI1 | 0.34 | 0.468521083 | | 0.11 | 16.94 | 17.04 | |
| 1221 | PBXIP1 | Q96AQ6 | 0.99 | 0.004364805 | | 0 | 15.83 | 15.83 | |
| 1222 | PCBP1 | Q15365 | 0.63 | 0.200659451 | | -0.02 | 17.05 | 17.03 | |
| 1223 | PCBP2 | Q15366 | 0.073 | 1.13667714 | | 0.07 | 16.34 | 16.4 | |
| 1224 | PCMT1 | P22061 | 0.35 | 0.455931956 | | 0.05 | 16.34 | 16.39 | |
| 1225 | PCNA | P12004 | 0.0083 | 2.080921908 | | -0.11 | 16.2 | 16.09 | |
| 1226 | PCNP | Q8WW12 | 0.98 | 0.008773924 | | 0 | 17.51 | 17.52 | |
| 1227 | PCYOX1 | Q9UHG3 | 0.0046 | 2.337242168 | | 0.24 | 14.65 | 14.88 | |
| 1228 | PDCD11 | Q14690 | 0.002 | 2.698970004 | | 0.21 | 15.27 | 15.48 | |
| 1229 | PDCD5 | O14737 | 0.22 | 0.657577319 | | -0.14 | 18 | 17.86 | |
| 1230 | PDCD6IP | Q8WUM4 | 0.0042 | 2.37675071 | | 0.09 | 17.22 | 17.31 | |
| 1231 | PDE12 | Q6L8Q7 | 0.31 | 0.508638306 | | 0.28 | 15.85 | 16.13 | |
| 1232 | PDHA1 | P08559 | 0.86 | 0.065501549 | | 0.01 | 16.25 | 16.26 | |
| 1233 | PDHB | P11177 | 0.9 | 0.045757491 | | 0 | 16.21 | 16.21 | |
| 1234 | PDIA3 | P30101 | 0.0001 | 4 | -0.15 | 17.61 | 17.46 | | |
| 1235 | PDIA4 | P13667 | 0.18 | 0.744727495 | | -0.04 | 16.87 | 16.83 | |
| 1236 | PDIA5 | Q14554 | 0.97 | 0.013228266 | | -0.02 | 16.63 | 16.61 | |
| 1237 | PDIA6 | Q15084 | 0.0052 | 2.283996656 | | -0.14 | 16.91 | 16.77 | |
| 1238 | PDLIM1 | O00151 | 0.045 | 1.346787486 | | -0.09 | 16.71 | 16.62 | |
| 1239 | PDLIM4 | P50479 | 0.91 | 0.040958608 | | -0.01 | 17.43 | 17.42 | |
| 1240 | PDLIM5 | Q96HC4 | 0.00064 | 3.193820026 | | 0.13 | 17.09 | 17.23 | |
| 1241 | PDLIM7 | Q9NR12 | 0.08 | 1.096910013 | | 0.16 | 17.07 | 17.22 | |
| 1242 | PDS5A | Q29RF7 | 0.04 | 1.397940009 | | 0.27 | 15.35 | 15.63 | |
| 1243 | PDS5B | Q9NTI5 | 0.094 | 1.026872146 | | 0.26 | 16.73 | 16.99 | |
| 1244 | PDXDC1 | Q6P996 | 0.76 | 0.119186408 | | -0.06 | 15.34 | 15.28 | |
| 1245 | PEA15 | Q15121 | 0.03 | 1.522878745 | | 0.11 | 16.06 | 16.17 | |
| 1246 | PEBP1 | P30086 | 0.75 | 0.124938737 | | -0.02 | 17.72 | 17.7 | |

| | | | | | | | |
|---|---|---|---|---|---|---|---|
| 1247 | PECAM1 | P16284 | 0.14 | 0.853871964 | -0.05 | 16.77 | 16.72 |
| 1248 | PELP1 | Q8IZL8 | 0.026 | 1.585026652 | 0.43 | 15.96 | 16.39 |
| 1249 | PEPD | P12955 | 0.0085 | 2.070581074 | -0.17 | 17.84 | 17.67 |
| 1250 | PES1 | O00541 | 0.75 | 0.124938737 | 0.04 | 16.07 | 16.11 |
| 1251 | PFAS | O15067 | 0.47 | 0.327902142 | -0.05 | 15.38 | 15.34 |
| 1252 | PFDN2 | Q9UHV9 | 0.48 | 0.318758763 | 0.1 | 15.32 | 15.42 |
| 1253 | PFDN6 | O15212 | 0.26 | 0.585026652 | 0.25 | 16.23 | 16.47 |
| 1254 | PFKL | P17858 | 0.33 | 0.48148606 | 0.04 | 15.35 | 15.39 |
| 1255 | PFKM | P08237 | 0.035 | 1.455931956 | 0.09 | 17.04 | 17.13 |
| 1256 | PFKP | Q01813 | 0.35 | 0.455931956 | 0.03 | 16.33 | 16.37 |
| 1257 | PFN1 | P07737 | 0.12 | 0.920818754 | 0.06 | 16.95 | 17.01 |
| 1258 | PFN2 | P35080 | 0.98 | 0.008773924 | 0.01 | 16.19 | 16.2 |
| 1259 | PGAM1 | P18669 | 0.2 | 0.698970004 | -0.16 | 17.15 | 17 |
| 1260 | PGD | P52209 | 0.043 | 1.366531544 | 0.07 | 16.24 | 16.3 |
| 1261 | PGK1 | P00558 | 0.12 | 0.920818754 | 0.04 | 17.39 | 17.43 |
| 1262 | PGLS | O95336 | 0.34 | 0.468521083 | 0.11 | 16.33 | 16.43 |
| 1263 | PGM1 | P36871 | 0.25 | 0.602059991 | 0.24 | 17.49 | 17.73 |
| 1264 | PGM2 | Q96G03 | 0.4 | 0.397940009 | -0.07 | 17.26 | 17.19 |
| 1265 | PGM2L1 | Q6PCE3 | 0.039 | 1.408935393 | 0.2 | 16.66 | 16.86 |
| 1266 | PGM3 | O95394 | 0.58 | 0.236572006 | 0.02 | 17.44 | 17.46 |
| 1267 | PGP | A6NDG6 | 0.42 | 0.37675071 | 0.09 | 15.97 | 16.06 |
| 1268 | PHB | P35232 | 0.032 | 1.494850022 | 0.08 | 16.25 | 16.34 |
| 1269 | PHB2 | Q99623 | 0.0015 | 2.823908741 | 0.11 | 17.06 | 17.17 |
| 1270 | PHF5A | Q7RTV0 | 0.5 | 0.301029996 | 0.11 | 18.23 | 18.34 |
| 1271 | PHGDH | O43175 | 0.46 | 0.337242168 | 0.09 | 16.63 | 16.73 |
| 1272 | PHLDB2 | Q86SQ0 | 0.68 | 0.167491087 | 0.03 | 16.62 | 16.65 |
| 1273 | PHPT1 | Q9NRX4 | 0.28 | 0.552841969 | 0.09 | 15.52 | 15.61 |
| 1274 | PI4K2A | Q9BTU6 | 0.15 | 0.823908741 | 0.22 | 15.17 | 15.39 |
| 1275 | PICALM | Q13492 | 0.018 | 1.744727495 | 0.24 | 15.94 | 16.18 |
| 1276 | PIGT | Q969N2 | 0.13 | 0.886056648 | 0.13 | 15.5 | 15.62 |
| 1277 | PIK3C3 | Q8NEB9 | 0.12 | 0.920818754 | 0.17 | 17.22 | 17.39 |
| 1278 | PIN1 | Q13526 | 0.48 | 0.318758763 | -0.08 | 16.55 | 16.48 |
| 1279 | PIR | O00625 | 0.0047 | 2.327902142 | -0.17 | 16.17 | 16 |
| 1280 | PITPNA | Q00169 | 0.19 | 0.721246399 | 0.05 | 17.71 | 17.76 |
| 1281 | PITPNB | P48739 | 0.22 | 0.657577319 | -0.04 | 16.38 | 16.34 |
| 1282 | PITRM1 | Q5JRX3 | 0.43 | 0.366531544 | 0.05 | 15.89 | 15.94 |
| 1283 | PKM | P14618 | 0.08 | 1.096910013 | -0.05 | 16.8 | 16.74 |
| 1284 | PLAA | Q9Y263 | 0.17 | 0.769551079 | -0.11 | 15.84 | 15.73 |
| 1285 | PLBD2 | Q8NHP8 | 0.051 | 1.292429824 | 0.13 | 17.1 | 17.23 |
| 1286 | PLCB3 | Q01970 | 0.65 | 0.187086643 | 0.08 | 15.41 | 15.48 |
| 1287 | PLD1 | Q13393 | 0.27 | 0.568636236 | 0.29 | 15.58 | 15.87 |
| 1288 | PLD3 | Q8IV08 | 0.0085 | 2.070581074 | 0.53 | 14.7 | 15.23 |
| 1289 | PLEC | Q15149 | 0.17 | 0.769551079 | 0.02 | 16.47 | 16.49 |
| 1290 | PLIN2 | Q99541 | NA | NA 0.39 | 16.51 | 16.9 | |
| 1291 | PLIN3 | O60664 | 0.028 | 1.552841969 | 0.08 | 16.18 | 16.26 |
| 1292 | PLOD1 | Q02809 | 0.00024 | 3.619788758 | 0.17 | 16.66 | 16.83 |
| 1293 | PLOD2 | O00469 | 0.14 | 0.853871964 | -0.07 | 16.4 | 16.33 |
| 1294 | PLOD3 | O60568 | 0.013 | 1.886056648 | 0.15 | 16.34 | 16.49 |
| 1295 | PLPBP | O94903 | 0.049 | 1.30980392 | -0.4 | 15.35 | 14.94 |
| 1296 | PLS3 | P13797 | 0.00089 | 3.050609993 | 0.1 | 16.95 | 17.05 |

| | | | | | | | |
|---|---|---|---|---|---|---|---|
| 1297 | PLXNA2 | O75051 | 0.15 | 0.823908741 | 0.45 | 13.66 | 14.12 |
| 1298 | PLXND1 | Q9Y4D7 | 0.022 | 1.657577319 | 0.15 | 14.35 | 14.5 |
| 1299 | PML | P29590 | 0.77 | 0.113509275 | 0 | 16.67 | 16.66 |
| 1300 | PMM2 | O15305 | 0.01 | 2 | -0.21 | 16.56 | 16.35 |
| 1301 | PMPCA | Q10713 | 0.7 | 0.15490196 | 0.04 | 15.31 | 15.35 |
| 1302 | PMPCB | O75439 | 0.34 | 0.468521083 | 0.08 | 16.87 | 16.95 |
| 1303 | PNN | Q9H307 | 0.15 | 0.823908741 | 0.09 | 14.29 | 14.38 |
| 1304 | PNP | P00491 | 0.001 | 3 | 0.13 | 16.43 | 16.55 |
| 1305 | PNPT1 | Q8TCS8 | 0.001 | 3 | 0.19 | 15.67 | 15.87 |
| 1306 | PODXL | O00592 | 0.00011 | 3.958607315 | 0.32 | 16.93 | 17.25 |
| 1307 | POFUT1 | Q9H488 | 0.74 | 0.13076828 | 0.02 | 15.58 | 15.6 |
| 1308 | POGLUT3 | Q7Z4H8 | 0.0077 | 2.113509275 | 0.43 | 15.82 | 16.25 |
| 1309 | POGZ | Q7Z3K3 | 0.74 | 0.13076828 | -0.04 | 13.53 | 13.49 |
| 1310 | POLDIP3 | Q9BY77 | 0.38 | 0.420216403 | 0.26 | 17.19 | 17.45 |
| 1311 | POLR1C | O15160 | 0.5 | 0.301029996 | 0.18 | 13.2 | 13.38 |
| 1312 | PON2 | Q15165 | 0.48 | 0.318758763 | -0.06 | 15.75 | 15.69 |
| 1313 | POR | P16435 | 0.12 | 0.920818754 | 0.14 | 15.2 | 15.34 |
| 1314 | PPA1 | Q15181 | 0.064 | 1.193820026 | -0.06 | 16.37 | 16.31 |
| 1315 | PPA2 | Q9H2U2 | 0.12 | 0.920818754 | -0.08 | 18.44 | 18.36 |
| 1316 | PPFIBP1 | Q86W92 | 0.29 | 0.537602002 | 0.21 | 15.56 | 15.77 |
| 1317 | PPIA | P62937 | 0.034 | 1.468521083 | -0.12 | 17.36 | 17.24 |
| 1318 | PPIB | P23284 | 0.063 | 1.200659451 | -0.07 | 18.28 | 18.21 |
| 1319 | PPIC | P45877 | 0.11 | 0.958607315 | -0.19 | 15.77 | 15.58 |
| 1320 | PPID | Q08752 | 0.54 | 0.26760624 | 0.04 | 16.55 | 16.59 |
| 1321 | PPIF | P30405 | 0.023 | 1.638272164 | 0.47 | 16.7 | 17.17 |
| 1322 | PPIH | O43447 | 0.27 | 0.568636236 | 0.05 | 16.97 | 17.02 |
| 1323 | PPIL1 | Q9Y3C6 | 1 | 0 | 0.01 | 17.67 | 17.67 |
| 1324 | PPM1F | P49593 | 0.038 | 1.420216403 | 0.15 | 15.46 | 15.61 |
| 1325 | PPM1G | O15355 | 0.1 | 1 | -0.18 | 16.38 | 16.2 |
| 1326 | PPME1 | Q9Y570 | 0.23 | 0.638272164 | -0.5 | 14.76 | 14.26 |
| 1327 | PPP1CA | P62136 | 0.74 | 0.13076828 | 0.01 | 15.82 | 15.83 |
| 1328 | PPP1R10 | Q96QC0 | 0.35 | 0.455931956 | 0.33 | 13.59 | 13.91 |
| 1329 | PPP1R12A | O14974 | 0.77 | 0.113509275 | 0.02 | 16.07 | 16.1 |
| 1330 | PPP1R18 | Q6NYC8 | 0.54 | 0.26760624 | -0.09 | 16.34 | 16.25 |
| 1331 | PPP1R7 | Q15435 | 0.46 | 0.337242168 | -0.12 | 15.81 | 15.69 |
| 1332 | PPP2CA | P67775 | 0.44 | 0.356547324 | 0.05 | 15.93 | 15.98 |
| 1333 | PPP2R1A | P30153 | 0.097 | 1.013228266 | 0.08 | 15.96 | 16.04 |
| 1334 | PPP2R1B | P30154 | 0.0074 | 2.13076828 | 0.41 | 15.68 | 16.09 |
| 1335 | PPP2R2A | P63151 | 0.51 | 0.292429824 | -0.05 | 16.19 | 16.14 |
| 1336 | PPP4R1 | Q8TF05 | 0.93 | 0.031517051 | 0.02 | 13.99 | 14.01 |
| 1337 | PPP5C | P53041 | 0.063 | 1.200659451 | -0.27 | 19.63 | 19.36 |
| 1338 | PPP6R3 | Q5H9R7 | 0.32 | 0.494850022 | 0.13 | 14.48 | 14.61 |
| 1339 | PPT1 | P50897 | 0.92 | 0.036212173 | 0 | 15.61 | 15.61 |
| 1340 | PRCC | Q92733 | 0.16 | 0.795880017 | 0.14 | 13.71 | 13.84 |
| 1341 | PRCP | P42785 | 0.43 | 0.366531544 | -0.06 | 15.92 | 15.87 |
| 1342 | PRDX1 | Q06830 | 0.76 | 0.119186408 | -0.02 | 18.46 | 18.44 |
| 1343 | PRDX2 | P32119 | 0.022 | 1.657577319 | 0.1 | 17.19 | 17.28 |
| 1344 | PRDX3 | P30048 | 0.45 | 0.346787486 | -0.03 | 17.36 | 17.33 |
| 1345 | PRDX4 | Q13162 | 0.28 | 0.552841969 | 0.05 | 16.27 | 16.32 |
| 1346 | PRDX5 | P30044 | 0.17 | 0.769551079 | 0.06 | 16.15 | 16.21 |

| | | | | | | | |
|---|---|---|---|---|---|---|---|
| 1347 | PRDX6 | P30041 | 0.78 | 0.107905397 | -0.01 | 16.67 | 16.66 |
| 1348 | PREP | P48147 | 0.24 | 0.619788758 | 0.07 | 16.24 | 16.31 |
| 1349 | PRKAA1 | Q13131 | 0.55 | 0.259637311 | 0.08 | 15.39 | 15.47 |
| 1350 | PRKACB | P22694 | 0.97 | 0.013228266 | -0.01 | 17.41 | 17.4 |
| 1351 | PRKAR1A | P10644 | 0.28 | 0.552841969 | -0.05 | 17.18 | 17.13 |
| 1352 | PRKAR2A | P13861 | 0.94 | 0.026872146 | 0 | 16.8 | 16.79 |
| 1353 | PRKAR2B | P31323 | 0.13 | 0.886056648 | 0.2 | 15.88 | 16.07 |
| 1354 | PRKCSH | P14314 | 0.68 | 0.167491087 | 0.01 | 17.23 | 17.24 |
| 1355 | PRKDC | P78527 | 0.0001 | 4 0.15 | 15.96 | 16.11 | |
| 1356 | PRMT1 | Q99873 | 0.09 | 1.045757491 | 0.1 | 16.6 | 16.7 |
| 1357 | PRPF19 | Q9UMS4 | 0.24 | 0.619788758 | 0.09 | 15.44 | 15.54 |
| 1358 | PRPF40A | O75400 | 0.53 | 0.27572413 | 0.06 | 16.14 | 16.2 |
| 1359 | PRPF6 | O94906 | 0.052 | 1.283996656 | -0.12 | 14.73 | 14.61 |
| 1360 | PRPF8 | Q6P2Q9 | 0.79 | 0.102372909 | 0.01 | 15.89 | 15.9 |
| 1361 | PRPS1 | P60891 | 0.049 | 1.30980392 | 0.16 | 15.32 | 15.47 |
| 1362 | PRPSAP1 | Q14558 | 0.96 | 0.017728767 | 0.04 | 14.45 | 14.49 |
| 1363 | PRRC1 | Q96M27 | 0.28 | 0.552841969 | 0.09 | 16.12 | 16.21 |
| 1364 | PRRC2C | Q9Y520 | 0.75 | 0.124938737 | -0.03 | 15.81 | 15.79 |
| 1365 | PRXL2A | Q9BRX8 | 0.7 | 0.15490196 | -0.01 | 16.71 | 16.7 |
| 1366 | PRXL2B | Q8TBF2 | 0.47 | 0.327902142 | 0.18 | 15.5 | 15.68 |
| 1367 | PSAP | P07602 | 0.51 | 0.292429824 | 0.04 | 17.12 | 17.16 |
| 1368 | PSAT1 | Q9Y617 | 0.24 | 0.619788758 | 0.12 | 15.53 | 15.65 |
| 1369 | PSIP1 | O75475 | 0.23 | 0.638272164 | -0.2 | 18.7 | 18.5 |
| 1370 | PSMA1 | P25786 | 0.53 | 0.27572413 | 0.04 | 17.35 | 17.39 |
| 1371 | PSMA2 | P25787 | 0.7 | 0.15490196 | 0.01 | 15.93 | 15.94 |
| 1372 | PSMA3 | P25788 | 0.87 | 0.060480747 | 0 | 17.66 | 17.66 |
| 1373 | PSMA4 | P25789 | 0.96 | 0.017728767 | 0.01 | 16.01 | 16.02 |
| 1374 | PSMA5 | P28066 | 0.16 | 0.795880017 | -0.06 | 16.92 | 16.86 |
| 1375 | PSMA6 | P60900 | 0.00098 | 3.008773924 | -0.11 | 16.69 | 16.57 |
| 1376 | PSMA7 | O14818 | 0.0043 | 2.366531544 | -0.13 | 16.83 | 16.7 |
| 1377 | PSMB1 | P20618 | 0.49 | 0.30980392 | 0.02 | 17.1 | 17.12 |
| 1378 | PSMB2 | P49721 | 0.12 | 0.920818754 | 0.21 | 16.47 | 16.67 |
| 1379 | PSMB3 | P49720 | 0.24 | 0.619788758 | -0.12 | 15.24 | 15.12 |
| 1380 | PSMB4 | P28070 | 0.28 | 0.552841969 | 0.06 | 16.4 | 16.46 |
| 1381 | PSMB5 | P28074 | 0.038 | 1.420216403 | 0.13 | 16.2 | 16.33 |
| 1382 | PSMB6 | P28072 | 0.25 | 0.602059991 | 0.15 | 14.82 | 14.97 |
| 1383 | PSMB7 | Q99436 | 0.00014 | 3.853871964 | -0.15 | 15.62 | 15.47 |
| 1384 | PSMC1 | P62191 | 0.2 | 0.698970004 | 0.04 | 16.18 | 16.23 |
| 1385 | PSMC2 | P35998 | 0.26 | 0.585026652 | -0.06 | 16.03 | 15.97 |
| 1386 | PSMC3 | P17980 | 0.32 | 0.494850022 | 0.03 | 15.6 | 15.63 |
| 1387 | PSMC4 | P43686 | 0.27 | 0.568636236 | 0.11 | 15.75 | 15.86 |
| 1388 | PSMC5 | P62195 | 0.26 | 0.585026652 | -0.04 | 15.92 | 15.88 |
| 1389 | PSMC6 | P62333 | 0.071 | 1.148741651 | -0.08 | 16.46 | 16.38 |
| 1390 | PSMD1 | Q99460 | 0.032 | 1.494850022 | 0.1 | 15.92 | 16.02 |
| 1391 | PSMD10 | O75832 | 0.037 | 1.431798276 | -0.14 | 17.41 | 17.27 |
| 1392 | PSMD11 | O00231 | 0.014 | 1.853871964 | 0.18 | 16.06 | 16.24 |
| 1393 | PSMD12 | O00232 | 0.15 | 0.823908741 | 0.14 | 16.74 | 16.88 |
| 1394 | PSMD13 | Q9UNM6 | 0.3 | 0.522878745 | -0.04 | 18.24 | 18.2 |
| 1395 | PSMD14 | O00487 | 1 | 0 0 | 16.79 | 16.79 | |
| 1396 | PSMD2 | Q13200 | 0.0032 | 2.494850022 | 0.12 | 16.17 | 16.29 |

| | | | | | | | |
|---|---|---|---|---|---|---|---|
| 1397 | PSMD3 | O43242 | 0.62 | 0.207608311 | 0.03 | 16.53 | 16.56 |
| 1398 | PSMD4 | P55036 | 0.77 | 0.113509275 | 0.02 | 16.39 | 16.41 |
| 1399 | PSMD5 | Q16401 | 0.3 | 0.522878745 | 0.08 | 15.08 | 15.16 |
| 1400 | PSMD7 | P51665 | 0.27 | 0.568636236 | -0.06 | 15.74 | 15.68 |
| 1401 | PSMD8 | P48556 | 0.61 | 0.214670165 | -0.03 | 15.77 | 15.75 |
| 1402 | PSME1 | Q06323 | 0.14 | 0.853871964 | -0.1 | 16.79 | 16.69 |
| 1403 | PSME2 | Q9UL46 | 0.35 | 0.455931956 | -0.04 | 16.78 | 16.74 |
| 1404 | PSME3 | P61289 | 0.0039 | 2.408935393 | -0.21 | 13.93 | 13.73 |
| 1405 | PSPC1 | Q8WXF1 | 0.33 | 0.48148606 | 0.1 | 14.93 | 15.03 |
| 1406 | PTBP1 | P26599 | 0.0036 | 2.443697499 | 0.15 | 15.5 | 15.65 |
| 1407 | PTGES3 | Q15185 | 0.0015 | 2.823908741 | -0.31 | 17.43 | 17.12 |
| 1408 | PTGR1 | Q14914 | 0.0001 | 4 | -0.25 | 17.29 | 17.04 |
| 1409 | PTGS1 | P23219 | 0.045 | 1.346787486 | 0.1 | 17.23 | 17.33 |
| 1410 | PTK2 | Q05397 | 0.25 | 0.602059991 | 0.08 | 16.03 | 16.11 |
| 1411 | PTMA | P06454 | 0.0066 | 2.180456064 | -0.21 | 17.49 | 17.28 |
| 1412 | PTMS | P20962 | 0.13 | 0.886056648 | -0.15 | 17.8 | 17.65 |
| 1413 | PTPA | Q15257 | 0.21 | 0.677780705 | 0.13 | 15.48 | 15.6 |
| 1414 | PTPN1 | P18031 | 0.32 | 0.494850022 | 0.14 | 17.01 | 17.15 |
| 1415 | PTPN11 | Q06124 | 0.73 | 0.13667714 | -0.04 | 16.37 | 16.34 |
| 1416 | PTPN12 | Q05209 | 0.65 | 0.187086643 | 0.09 | 14.65 | 14.73 |
| 1417 | PTPN23 | Q9H3S7 | 0.3 | 0.522878745 | 0.08 | 15.33 | 15.42 |
| 1418 | PTPRB | P23467 | 0.25 | 0.602059991 | 0.17 | 17.08 | 17.25 |
| 1419 | PTRH2 | Q9Y3E5 | 0.25 | 0.602059991 | 0.17 | 16.28 | 16.45 |
| 1420 | PUF60 | Q9UHX1 | 0.23 | 0.638272164 | 0.06 | 16.03 | 16.09 |
| 1421 | PUM1 | Q14671 | 0.69 | 0.161150909 | -0.04 | 16.89 | 16.85 |
| 1422 | PUM3 | Q15397 | 0.28 | 0.552841969 | 0.11 | 15 | 15.11 |
| 1423 | PWP2 | Q15269 | 0.31 | 0.508638306 | 0.71 | 13.63 | 14.34 |
| 1424 | PXDN | Q92626 | 0.38 | 0.420216403 | 0.03 | 16 | 16.03 |
| 1425 | PXN | P49023 | 0.031 | 1.508638306 | -0.1 | 16.9 | 16.8 |
| 1426 | PYCR1 | P32322 | 0.11 | 0.958607315 | 0.22 | 16.51 | 16.73 |
| 1427 | PYGB | P11216 | 0.038 | 1.420216403 | 0.21 | 16.88 | 17.08 |
| 1428 | PYGL | P06737 | 0.71 | 0.148741651 | 0.03 | 16.77 | 16.8 |
| 1429 | PYM1 | Q9BRP8 | 0.32 | 0.494850022 | 0.22 | 14.21 | 14.43 |
| 1430 | QARS | P47897 | 0.15 | 0.823908741 | 0.07 | 15.58 | 15.65 |
| 1431 | QRICH1 | Q2TAL8 | 0.87 | 0.060480747 | 0.02 | 17.19 | 17.21 |
| 1432 | RAB10 | P61026 | 0.33 | 0.48148606 | 0.03 | 17.5 | 17.53 |
| 1433 | RAB11B | Q15907 | 0.088 | 1.055517328 | -0.16 | 18.65 | 18.49 |
| 1434 | RAB13 | P51153 | 0.97 | 0.013228266 | 0 | 17.13 | 17.13 |
| 1435 | RAB14 | P61106 | 0.17 | 0.769551079 | 0.18 | 15.38 | 15.56 |
| 1436 | RAB18 | Q9NP72 | 0.66 | 0.180456064 | 0.04 | 15.9 | 15.94 |
| 1437 | RAB1A | P62820 | 0.01 | 2 | -0.11 | 15.99 | 15.87 |
| 1438 | RAB1B | Q9H0U4 | 0.22 | 0.657577319 | -0.11 | 15.09 | 14.98 |
| 1439 | RAB21 | Q9UL25 | 0.21 | 0.677780705 | -0.07 | 17.94 | 17.87 |
| 1440 | RAB2A | P61019 | 0.12 | 0.920818754 | 0.14 | 17.14 | 17.28 |
| 1441 | RAB32 | Q13637 | 0.022 | 1.657577319 | 0.19 | 16.09 | 16.27 |
| 1442 | RAB35 | Q15286 | 0.42 | 0.37675071 | -0.23 | 16.94 | 16.71 |
| 1443 | RAB3GAP1 | Q15042 | 0.35 | 0.455931956 | 0.06 | 15.42 | 15.48 |
| 1444 | RAB3GAP2 | Q9H2M9 | 0.051 | 1.292429824 | 0.18 | 15.47 | 15.65 |
| 1445 | RAB5A | P20339 | 0.011 | 1.958607315 | 0.24 | 15.58 | 15.82 |
| 1446 | RAB5B | P61020 | 0.19 | 0.721246399 | 0.17 | 15.29 | 15.46 |

| | | | | | | | |
|---|---|---|---|---|---|---|---|
| 1447 | RAB5C | P51148 | 0.55 | 0.259637311 | -0.06 | 15.9 | 15.84 |
| 1448 | RAB6A | P20340 | 0.83 | 0.080921908 | -0.03 | 16.45 | 16.42 |
| 1449 | RAB7A | P51149 | 0.3 | 0.522878745 | -0.06 | 17.36 | 17.3 |
| 1450 | RAB8A | P61006 | 0.47 | 0.327902142 | 0.07 | 15.83 | 15.9 |
| 1451 | RAC1 | P63000 | 0.013 | 1.886056648 | 0.26 | 17.42 | 17.67 |
| 1452 | RAC2 | P15153 | 0.62 | 0.207608311 | 0.06 | 16.68 | 16.74 |
| 1453 | RACK1 | P63244 | 0.33 | 0.48148606 | 0.04 | 17.23 | 17.26 |
| 1454 | RAD23A | P54725 | 0.13 | 0.886056648 | 0.29 | 12.66 | 12.95 |
| 1455 | RAD23B | P54727 | 0.89 | 0.050609993 | -0.02 | 16.34 | 16.33 |
| 1456 | RAD50 | Q92878 | 0.43 | 0.366531544 | 0.13 | 13.75 | 13.88 |
| 1457 | RAE1 | P78406 | 0.079 | 1.102372909 | 0.27 | 18.2 | 18.47 |
| 1458 | RAI14 | Q9P0K7 | 0.062 | 1.207608311 | 0.22 | 16.44 | 16.66 |
| 1459 | RALA | P11233 | 0.11 | 0.958607315 | -0.07 | 18.19 | 18.12 |
| 1460 | RALB | P11234 | 0.2 | 0.698970004 | -0.07 | 18.35 | 18.29 |
| 1461 | RALY | Q9UKM9 | 0.58 | 0.236572006 | -0.03 | 18.19 | 18.17 |
| 1462 | RAN | P62826 | 0.91 | 0.040958608 | 0.01 | 17.98 | 17.98 |
| 1463 | RANBP1 | P43487 | 0.00047 | 3.327902142 | -0.24 | 17.86 | 17.62 |
| 1464 | RANBP2 | P49792 | 0.0001 | 4 | 0.18 | 16.49 | 16.67 |
| 1465 | RANBP3 | Q9H6Z4 | 0.044 | 1.356547324 | 0.17 | 16.89 | 17.06 |
| 1466 | RANBP6 | O60518 | 0.16 | 0.795880017 | 0.58 | 13.81 | 14.39 |
| 1467 | RANGAP1 | P46060 | 0.14 | 0.853871964 | 0.08 | 15.43 | 15.51 |
| 1468 | RAP1B | P61224 | 0.0015 | 2.823908741 | 0.15 | 16.08 | 16.23 |
| 1469 | RAP1GDS1 | P52306 | 0.061 | 1.214670165 | 0.3 | 15.43 | 15.73 |
| 1470 | RAP2C | Q9Y3L5 | 0.73 | 0.13667714 | 0.05 | 15.82 | 15.88 |
| 1471 | RAPH1 | Q70E73 | 0.11 | 0.958607315 | 0.32 | 17.06 | 17.38 |
| 1472 | RARS | P54136 | 0.38 | 0.420216403 | -0.03 | 15.75 | 15.72 |
| 1473 | RASA1 | P20936 | 0.99 | 0.004364805 | 0.03 | 13.6 | 13.63 |
| 1474 | RASIP1 | Q5U651 | 0.22 | 0.657577319 | 0.08 | 15.98 | 16.06 |
| 1475 | RBBP4 | Q09028 | 0.18 | 0.744727495 | 0.11 | 16.25 | 16.36 |
| 1476 | RBBP7 | Q16576 | 0.15 | 0.823908741 | -0.19 | 16.1 | 15.91 |
| 1477 | RBM12 | Q9NTZ6 | 0.36 | 0.443697499 | -0.06 | 14.8 | 14.74 |
| 1478 | RBM14 | Q96PK6 | 0.042 | 1.37675071 | 0.09 | 18.17 | 18.26 |
| 1479 | RBM17 | Q96I25 | 0.26 | 0.585026652 | 0.19 | 16.3 | 16.49 |
| 1480 | RBM25 | P49756 | 0.061 | 1.214670165 | -0.17 | 15.33 | 15.16 |
| 1481 | RBM28 | Q9NW13 | 0.25 | 0.602059991 | 0.22 | 17.29 | 17.51 |
| 1482 | RBM3 | P98179 | 0.11 | 0.958607315 | -0.2 | 16.72 | 16.52 |
| 1483 | RBM34 | P42696 | 0.073 | 1.13667714 | 0.27 | 14.82 | 15.09 |
| 1484 | RBM39 | Q14498 | 0.18 | 0.744727495 | -0.08 | 15.89 | 15.81 |
| 1485 | RBM4 | Q9BWF3 | 0.034 | 1.468521083 | 0.24 | 15.08 | 15.32 |
| 1486 | RBM8A | Q9Y5S9 | 0.0016 | 2.795880017 | -0.18 | 16.44 | 16.26 |
| 1487 | RBMS1 | P29558 | 0.044 | 1.356547324 | -0.14 | 18.22 | 18.08 |
| 1488 | RBMX | P38159 | 0.29 | 0.537602002 | 0.05 | 17.74 | 17.79 |
| 1489 | RCC1 | P18754 | 0.81 | 0.091514981 | 0.01 | 16.71 | 16.72 |
| 1490 | RCC2 | Q9P258 | 0.68 | 0.167491087 | 0.04 | 18.01 | 18.04 |
| 1491 | RCL1 | Q9Y2P8 | 0.54 | 0.26760624 | -0.34 | 17.5 | 17.15 |
| 1492 | RCN1 | Q15293 | 0.0001 | 4 | -0.15 | 16.68 | 16.53 |
| 1493 | RCN2 | Q14257 | 0.056 | 1.251811973 | -0.19 | 14.67 | 14.48 |
| 1494 | RCN3 | Q96D15 | 0.33 | 0.48148606 | 0.07 | 16.1 | 16.17 |
| 1495 | RDX | P35241 | 0.32 | 0.494850022 | 0.04 | 15.76 | 15.8 |
| 1496 | RECK | O95980 | 0.061 | 1.214670165 | 0.13 | 15.36 | 15.49 |

| | | | | | | | |
|---|---|---|---|---|---|---|---|
| 1497 | RECQL | P46063 | 0.67 | 0.173925197 | 0.03 | 17.1 | 17.13 |
| 1498 | REEP5 | Q00765 | 0.29 | 0.537602002 | -0.1 | 19.46 | 19.36 |
| 1499 | RELA | Q04206 | 0.014 | 1.853871964 | -0.22 | 15.47 | 15.25 |
| 1500 | RELCH | Q9P260 | 0.57 | 0.244125144 | 0.05 | 13.66 | 13.71 |
| 1501 | REPS1 | Q96D71 | 0.25 | 0.602059991 | -0.08 | 16.48 | 16.4 |
| 1502 | RER1 | O15258 | 0.31 | 0.508638306 | -0.13 | 18.99 | 18.85 |
| 1503 | REST | Q13127 | 0.71 | 0.148741651 | -0.14 | 19.31 | 19.17 |
| 1504 | REXO2 | Q9Y3B8 | 0.42 | 0.37675071 | -0.09 | 16.18 | 16.09 |
| 1505 | RHOA | P61586 | 0.28 | 0.552841969 | 0.1 | 15.77 | 15.87 |
| 1506 | RHOB | P62745 | NA | NA | 0.29 | 15.01 | 15.3 |
| 1507 | RHOC | P08134 | 0.28 | 0.552841969 | -0.29 | 16.55 | 16.26 |
| 1508 | RHOG | P84095 | 0.62 | 0.207608311 | 0.05 | 17.47 | 17.52 |
| 1509 | RHOT2 | Q8IXI1 | 0.41 | 0.387216143 | -0.09 | 15.08 | 15 |
| 1510 | RIC8A | Q9NPQ8 | 0.87 | 0.060480747 | 0.01 | 12.93 | 12.95 |
| 1511 | RIPK1 | Q13546 | 0.65 | 0.187086643 | 0.05 | 15.82 | 15.87 |
| 1512 | RIPOR1 | Q6ZS17 | 0.95 | 0.022276395 | 0.01 | 14.18 | 14.2 |
| 1513 | RNF213 | Q63HN8 | NA | NA | 0.41 | 13.86 | 14.27 |
| 1514 | RNH1 | P13489 | 0.92 | 0.036212173 | -0.01 | 15.55 | 15.55 |
| 1515 | RNPEP | Q9H4A4 | 0.0093 | 2.031517051 | 0.17 | 15.42 | 15.59 |
| 1516 | RNPS1 | Q15287 | 0.17 | 0.769551079 | -0.11 | 17.62 | 17.51 |
| 1517 | RO60 | P10155 | 0.0053 | 2.27572413 | 0.16 | 16.04 | 16.21 |
| 1518 | RP2 | O75695 | 0.074 | 1.13076828 | 0.12 | 17 | 17.12 |
| 1519 | RPA1 | P27694 | 0.23 | 0.638272164 | 0.06 | 16.26 | 16.32 |
| 1520 | RPA2 | P15927 | 0.46 | 0.337242168 | -0.07 | 14.55 | 14.49 |
| 1521 | RPL10 | P27635 | 0.35 | 0.455931956 | 0.06 | 15.6 | 15.65 |
| 1522 | RPL10A | P62906 | 0.29 | 0.537602002 | 0.05 | 17.45 | 17.5 |
| 1523 | RPL11 | P62913 | 0.31 | 0.508638306 | 0.12 | 17.97 | 18.09 |
| 1524 | RPL12 | P30050 | 0.47 | 0.327902142 | 0.04 | 17.31 | 17.36 |
| 1525 | RPL13 | P26373 | 0.45 | 0.346787486 | -0.07 | 18.46 | 18.38 |
| 1526 | RPL13A | P40429 | 0.3 | 0.522878745 | -0.08 | 19.12 | 19.04 |
| 1527 | RPL14 | P50914 | 0.74 | 0.13076828 | -0.02 | 18.18 | 18.16 |
| 1528 | RPL15 | P61313 | 0.038 | 1.420216403 | 0.15 | 16.83 | 16.98 |
| 1529 | RPL17 | P18621 | 0.5 | 0.301029996 | -0.05 | 17.19 | 17.14 |
| 1530 | RPL18 | Q07020 | 0.14 | 0.853871964 | 0.06 | 17.12 | 17.19 |
| 1531 | RPL18A | Q02543 | 0.047 | 1.327902142 | 0.24 | 17.27 | 17.51 |
| 1532 | RPL19 | P84098 | 0.14 | 0.853871964 | -0.1 | 18.06 | 17.96 |
| 1533 | RPL21 | P46778 | 0.42 | 0.37675071 | -0.05 | 17.11 | 17.06 |
| 1534 | RPL22 | P35268 | 0.13 | 0.886056648 | 0.03 | 15.6 | 15.62 |
| 1535 | RPL23 | P62829 | 0.35 | 0.455931956 | 0.07 | 16.96 | 17.03 |
| 1536 | RPL23A | P62750 | 0.0001 | 4 | -0.18 | 17.78 | 17.6 |
| 1537 | RPL24 | P83731 | 0.92 | 0.036212173 | -0.01 | 18.13 | 18.12 |
| 1538 | RPL26 | P61254 | 0.38 | 0.420216403 | -0.06 | 19.24 | 19.18 |
| 1539 | RPL27 | P61353 | 0.068 | 1.167491087 | 0.12 | 18.91 | 19.03 |
| 1540 | RPL27A | P46776 | 0.75 | 0.124938737 | 0.04 | 18.13 | 18.17 |
| 1541 | RPL28 | P46779 | 0.17 | 0.769551079 | 0.12 | 18.46 | 18.59 |
| 1542 | RPL29 | P47914 | 0.78 | 0.107905397 | 0.02 | 18.04 | 18.06 |
| 1543 | RPL3 | P39023 | 0.51 | 0.292429824 | 0.03 | 17.43 | 17.46 |
| 1544 | RPL30 | P62888 | 0.42 | 0.37675071 | -0.04 | 17.24 | 17.2 |
| 1545 | RPL31 | P62899 | 0.066 | 1.180456064 | 0.11 | 17.23 | 17.34 |
| 1546 | RPL32 | P62910 | 0.83 | 0.080921908 | 0.01 | 17.75 | 17.76 |

| | | | | | | | |
|---|---|---|---|---|---|---|---|
| 1547 | RPL34 | P49207 | 0.063 | 1.200659451 | -0.18 | 18.41 | 18.23 |
| 1548 | RPL35 | P42766 | 0.9 | 0.045757491 | -0.05 | 17.58 | 17.53 |
| 1549 | RPL35A | P18077 | 0.12 | 0.920818754 | -0.25 | 19.57 | 19.32 |
| 1550 | RPL36A | P83881 | 0.11 | 0.958607315 | -0.11 | 19.56 | 19.45 |
| 1551 | RPL37 | P61927 | 0.12 | 0.920818754 | -0.18 | 20.14 | 19.95 |
| 1552 | RPL37A | P61513 | 0.11 | 0.958607315 | -0.08 | 18.3 | 18.21 |
| 1553 | RPL38 | P63173 | 0.15 | 0.823908741 | 0.16 | 18.78 | 18.94 |
| 1554 | RPL4 | P36578 | 0.73 | 0.13667714 | 0.02 | 16.88 | 16.9 |
| 1555 | RPL5 | P46777 | 0.12 | 0.920818754 | 0.07 | 17.85 | 17.92 |
| 1556 | RPL6 | Q02878 | 0.43 | 0.366531544 | -0.06 | 17.7 | 17.64 |
| 1557 | RPL7 | P18124 | 0.2 | 0.698970004 | -0.05 | 17.43 | 17.37 |
| 1558 | RPL7A | P62424 | 0.79 | 0.102372909 | -0.01 | 18.06 | 18.06 |
| 1559 | RPL7L1 | Q6DKI1 | 0.35 | 0.455931956 | 0.19 | 16.42 | 16.61 |
| 1560 | RPL8 | P62917 | 0.59 | 0.229147988 | 0.02 | 18.12 | 18.14 |
| 1561 | RPL9 | P32969 | 0.096 | 1.017728767 | 0.09 | 16.39 | 16.47 |
| 1562 | RPLP0 | P05388 | 0.13 | 0.886056648 | 0.07 | 17.05 | 17.12 |
| 1563 | RPLP1 | P05386 | 0.0055 | 2.259637311 | -0.14 | 16.07 | 15.93 |
| 1564 | RPLP2 | P05387 | 0.51 | 0.292429824 | -0.06 | 16.48 | 16.42 |
| 1565 | RPN1 | P04843 | 0.46 | 0.337242168 | 0.02 | 16.77 | 16.79 |
| 1566 | RPN2 | P04844 | 0.073 | 1.13667714 | 0.07 | 15.59 | 15.67 |
| 1567 | RPRD1B | Q9NQG5 | 0.36 | 0.443697499 | 0.23 | 14.08 | 14.31 |
| 1568 | RPS10 | P46783 | 0.34 | 0.468521083 | -0.08 | 17.87 | 17.79 |
| 1569 | RPS11 | P62280 | 0.39 | 0.408935393 | 0.05 | 19.01 | 19.06 |
| 1570 | RPS12 | P25398 | 0.55 | 0.259637311 | 0.03 | 15.14 | 15.17 |
| 1571 | RPS13 | P62277 | 0.19 | 0.721246399 | -0.05 | 17.49 | 17.44 |
| 1572 | RPS14 | P62263 | 0.17 | 0.769551079 | 0.15 | 16.3 | 16.45 |
| 1573 | RPS15 | P62841 | 0.97 | 0.013228266 | 0 | 15.93 | 15.92 |
| 1574 | RPS15A | P62244 | 0.045 | 1.346787486 | 0.33 | 18.29 | 18.61 |
| 1575 | RPS16 | P62249 | 0.34 | 0.468521083 | 0.03 | 17.89 | 17.92 |
| 1576 | RPS17 | P08708 | 0.088 | 1.055517328 | -0.09 | 15.34 | 15.26 |
| 1577 | RPS18 | P62269 | 0.96 | 0.017728767 | 0.01 | 17.43 | 17.43 |
| 1578 | RPS19 | P39019 | 0.14 | 0.853871964 | -0.07 | 18.07 | 18 |
| 1579 | RPS2 | P15880 | 0.18 | 0.744727495 | 0.04 | 17.65 | 17.69 |
| 1580 | RPS20 | P60866 | 0.42 | 0.37675071 | 0.1 | 17.85 | 17.95 |
| 1581 | RPS21 | P63220 | 0.72 | 0.142667504 | -0.04 | 19.81 | 19.77 |
| 1582 | RPS23 | P62266 | 0.97 | 0.013228266 | 0 | 16.39 | 16.39 |
| 1583 | RPS24 | P62847 | 0.62 | 0.207608311 | 0.06 | 17.13 | 17.19 |
| 1584 | RPS25 | P62851 | 0.29 | 0.537602002 | 0.14 | 18.47 | 18.61 |
| 1585 | RPS26 | P62854 | 0.94 | 0.026872146 | 0 | 17.11 | 17.11 |
| 1586 | RPS27A | P62979 | 0.86 | 0.065501549 | -0.02 | 18.02 | 18 |
| 1587 | RPS27L | Q71UM5 | 0.24 | 0.619788758 | -0.17 | 18.19 | 18.03 |
| 1588 | RPS28 | P62857 | 0.55 | 0.259637311 | -0.01 | 17.61 | 17.6 |
| 1589 | RPS3 | P23396 | 0.21 | 0.677780705 | 0.05 | 18.04 | 18.09 |
| 1590 | RPS3A | P61247 | 0.51 | 0.292429824 | -0.03 | 17.97 | 17.94 |
| 1591 | RPS4X | P62701 | 0.0012 | 2.920818754 | 0.16 | 17.16 | 17.32 |
| 1592 | RPS4Y1 | P22090 | 0.033 | 1.48148606 | 0.38 | 16.3 | 16.67 |
| 1593 | RPS5 | P46782 | 0.65 | 0.187086643 | 0.07 | 14.79 | 14.86 |
| 1594 | RPS6 | P62753 | 0.48 | 0.318758763 | 0.04 | 16.66 | 16.7 |
| 1595 | RPS6KA3 | P51812 | 0.83 | 0.080921908 | -0.03 | 18.02 | 17.99 |
| 1596 | RPS7 | P62081 | 0.26 | 0.585026652 | 0.04 | 15.77 | 15.81 |

| | | | | | | | |
|---|---|---|---|---|---|---|---|
| 1597 | RPS8 | P62241 | 0.32 | 0.494850022 | -0.08 | 17.69 | 17.6 |
| 1598 | RPS9 | P46781 | 0.7 | 0.15490196 | 0.04 | 17.53 | 17.57 |
| 1599 | RPSA | P08865 | 0.13 | 0.886056648 | 0.24 | 16.64 | 16.88 |
| 1600 | RRAGC | Q9HB90 | 0.35 | 0.455931956 | 0.14 | 15.02 | 15.16 |
| 1601 | RRAS | P10301 | 0.13 | 0.886056648 | -0.1 | 16.82 | 16.72 |
| 1602 | RRBP1 | Q9P2E9 | 0.0001 | 4 -0.22 | 17.95 | 17.73 | |
| 1603 | RRM1 | P23921 | 0.28 | 0.552841969 | -0.09 | 16.52 | 16.43 |
| 1604 | RRM2B | Q7LG56 | 0.0056 | 2.251811973 | 0.43 | 16.42 | 16.85 |
| 1605 | RRP1 | P56182 | 0.52 | 0.283996656 | -0.07 | 16.73 | 16.66 |
| 1606 | RRP12 | Q5JTH9 | 0.3 | 0.522878745 | 0.1 | 14.63 | 14.73 |
| 1607 | RRS1 | Q15050 | 0.62 | 0.207608311 | -0.06 | 15.09 | 15.03 |
| 1608 | RSL1D1 | O76021 | 0.0001 | 4 0.29 | 16.34 | 16.63 | |
| 1609 | RSU1 | Q15404 | 0.4 | 0.397940009 | 0.06 | 15.56 | 15.62 |
| 1610 | RTCA | O00442 | 0.26 | 0.585026652 | 0.21 | 16.67 | 16.88 |
| 1611 | RTCB | Q9Y3I0 | 0.018 | 1.744727495 | 0.07 | 17.32 | 17.39 |
| 1612 | RTN4 | Q9NQC3 | 0.0017 | 2.769551079 | -0.2 | 15.71 | 15.51 |
| 1613 | RTRAF | Q9Y224 | 0.04 | 1.397940009 | 0.32 | 13.5 | 13.82 |
| 1614 | RUVBL1 | Q9Y265 | 0.034 | 1.468521083 | 0.08 | 16.23 | 16.31 |
| 1615 | RUVBL2 | Q9Y230 | 0.15 | 0.823908741 | 0.05 | 17.42 | 17.47 |
| 1616 | S100A10 | P60903 | 0.12 | 0.920818754 | -0.09 | 17.48 | 17.39 |
| 1617 | S100A11 | P31949 | 0.54 | 0.26760624 | 0.07 | 16.15 | 16.22 |
| 1618 | S100A13 | Q99584 | 0.0058 | 2.236572006 | 0.19 | 18.86 | 19.05 |
| 1619 | S100A16 | Q96FQ6 | 0.26 | 0.585026652 | -0.12 | 16.89 | 16.77 |
| 1620 | S100A6 | P06703 | 0.2 | 0.698970004 | 0.68 | 16.61 | 17.29 |
| 1621 | SACM1L | Q9NTJ5 | 0.35 | 0.455931956 | 0.13 | 14.92 | 15.05 |
| 1622 | SAE1 | Q9UBE0 | 0.016 | 1.795880017 | 0.11 | 17.36 | 17.47 |
| 1623 | SAFB | Q15424 | 0.029 | 1.537602002 | -0.1 | 17.39 | 17.28 |
| 1624 | SAMHD1 | Q9Y3Z3 | 0.53 | 0.27572413 | 0.04 | 15.8 | 15.84 |
| 1625 | SAMM50 | Q9Y512 | 0.23 | 0.638272164 | 0.14 | 14.92 | 15.06 |
| 1626 | SAP18 | O00422 | 0.41 | 0.387216143 | -0.1 | 15.24 | 15.14 |
| 1627 | SAP30BP | Q9UHR5 | 0.74 | 0.13076828 | 0.05 | 16.41 | 16.45 |
| 1628 | SAR1A | Q9NR31 | 0.036 | 1.443697499 | 0.09 | 15.82 | 15.91 |
| 1629 | SARNP | P82979 | 0.72 | 0.142667504 | 0.03 | 17.89 | 17.91 |
| 1630 | SARS | P49591 | 0.0085 | 2.070581074 | -0.08 | 15.98 | 15.9 |
| 1631 | SART1 | O43290 | 0.016 | 1.795880017 | -0.15 | 14.63 | 14.48 |
| 1632 | SBDS | Q9Y3A5 | 0.3 | 0.522878745 | 0.06 | 17 | 17.05 |
| 1633 | SCARB2 | Q14108 | 0.53 | 0.27572413 | -0.04 | 17.59 | 17.55 |
| 1634 | SCCPDH | Q8NBX0 | 0.015 | 1.823908741 | 0.26 | 15.67 | 15.93 |
| 1635 | SCFD1 | Q8WVM8 | 0.095 | 1.022276395 | 0.16 | 14.57 | 14.73 |
| 1636 | SCN1A | P35498 | 0.92 | 0.036212173 | 0.02 | 19.21 | 19.23 |
| 1637 | SCP2 | P22307 | 0.81 | 0.091514981 | 0 | 16.72 | 16.71 |
| 1638 | SCPEP1 | Q9HB40 | 0.53 | 0.27572413 | -0.06 | 15.45 | 15.39 |
| 1639 | SCRN1 | Q12765 | 0.71 | 0.148741651 | -0.03 | 16.24 | 16.21 |
| 1640 | SDCBP | O00560 | 0.11 | 0.958607315 | -0.11 | 15.62 | 15.52 |
| 1641 | SDHA | P31040 | 0.44 | 0.356547324 | 0.03 | 16.15 | 16.18 |
| 1642 | SDHB | P21912 | 0.25 | 0.602059991 | 0.09 | 16.13 | 16.22 |
| 1643 | SDSL | Q96GA7 | 0.79 | 0.102372909 | 0.04 | 13.87 | 13.91 |
| 1644 | SEC13 | P55735 | 0.38 | 0.420216403 | 0.03 | 17.11 | 17.14 |
| 1645 | SEC16A | O15027 | 0.96 | 0.017728767 | 0 | 16.24 | 16.23 |
| 1646 | SEC22B | O75396 | 0.057 | 1.244125144 | -0.1 | 17.29 | 17.19 |

| | | | | | | | |
|---|---|---|---|---|---|---|---|
| 1647 | SEC23A | Q15436 | 0.31 | 0.508638306 | 0.06 | 15.96 | 16.02 |
| 1648 | SEC23IP | Q9Y6Y8 | 0.72 | 0.142667504 | 0.05 | 16.22 | 16.27 |
| 1649 | SEC24B | O95487 | 0.06 | 1.22184875 | 0.26 | 15.97 | 16.23 |
| 1650 | SEC24C | P53992 | 0.00011 | 3.958607315 | 0.21 | 15.64 | 15.84 |
| 1651 | SEC24D | O94855 | 0.38 | 0.420216403 | -0.05 | 15.53 | 15.49 |
| 1652 | SEC31A | O94979 | 0.0038 | 2.420216403 | 0.18 | 17.27 | 17.45 |
| 1653 | SEC61A1 | P61619 | 0.87 | 0.060480747 | 0.02 | 16.81 | 16.83 |
| 1654 | SEC61B | P60468 | NA | NA | 0.23 | 17.47 | 17.7 |
| 1655 | SEC63 | Q9UGP8 | 0.0054 | 2.26760624 | 0.23 | 16.29 | 16.52 |
| 1656 | SEL1L | Q9UBV2 | 0.68 | 0.167491087 | 0.05 | 13.42 | 13.48 |
| 1657 | SELENOF | O60613 | 0.033 | 1.48148606 | -0.33 | 16.1 | 15.78 |
| 1658 | SEPHS1 | P49903 | 0.3 | 0.522878745 | 0.16 | 15.74 | 15.91 |
| 1659 | SEPTIN11 | Q9NVA2 | 0.072 | 1.142667504 | 0.25 | 16.9 | 17.15 |
| 1660 | SEPTIN2 | Q15019 | 0.61 | 0.214670165 | 0.03 | 16.47 | 16.5 |
| 1661 | SEPTIN6 | Q14141 | 0.054 | 1.26760624 | 0.24 | 15.74 | 15.98 |
| 1662 | SEPTIN7 | Q16181 | 0.099 | 1.004364805 | 0.06 | 16.94 | 17 |
| 1663 | SEPTIN9 | Q9UHD8 | 0.085 | 1.070581074 | 0.27 | 15.94 | 16.2 |
| 1664 | SERBP1 | Q8NC51 | 0.00049 | 3.30980392 | -0.15 | 17.16 | 17.01 |
| 1665 | SERPINB1 | P30740 | 0.76 | 0.119186408 | 0.05 | 15.67 | 15.72 |
| 1666 | SERPINB2 | P05120 | 0.0001 | 4 | -0.19 | 16.14 | 15.95 |
| 1667 | SERPINB6 | P35237 | 0.14 | 0.853871964 | 0.07 | 16.44 | 16.52 |
| 1668 | SERPINB8 | P50452 | 0.77 | 0.113509275 | -0.05 | 16.13 | 16.08 |
| 1669 | SERPINB9 | P50453 | 0.35 | 0.455931956 | 0.06 | 17.18 | 17.23 |
| 1670 | SERPINE1 | P05121 | 0.32 | 0.494850022 | 0.03 | 16.52 | 16.55 |
| 1671 | SERPINH1 | P50454 | 0.97 | 0.013228266 | 0 | 17.38 | 17.38 |
| 1672 | SET | Q01105 | 0.19 | 0.721246399 | -0.07 | 17.09 | 17.02 |
| 1673 | SF1 | Q15637 | 0.64 | 0.193820026 | 0.02 | 15.66 | 15.68 |
| 1674 | SF3A1 | Q15459 | 0.19 | 0.721246399 | -0.1 | 15.97 | 15.87 |
| 1675 | SF3A2 | Q15428 | 0.77 | 0.113509275 | 0.03 | 16.96 | 16.99 |
| 1676 | SF3A3 | Q12874 | 0.53 | 0.27572413 | 0.18 | 15.09 | 15.26 |
| 1677 | SF3B1 | O75533 | 0.34 | 0.468521083 | 0.05 | 15.9 | 15.94 |
| 1678 | SF3B2 | Q13435 | 0.0043 | 2.366531544 | -0.21 | 16.59 | 16.39 |
| 1679 | SF3B3 | Q15393 | 0.0001 | 4 | 0.15 | 16.34 | 16.49 |
| 1680 | SF3B4 | Q15427 | 0.096 | 1.017728767 | -0.2 | 16.09 | 15.89 |
| 1681 | SFPQ | P23246 | 0.18 | 0.744727495 | 0.05 | 17.23 | 17.28 |
| 1682 | SFXN1 | Q9H9B4 | 0.035 | 1.455931956 | 0.17 | 14.95 | 15.12 |
| 1683 | SFXN3 | Q9BWM7 | 0.1 | 1 | 0.24 | 15.63 | 15.87 |
| 1684 | SGTA | O43765 | 0.08 | 1.096910013 | 0.22 | 17.06 | 17.28 |
| 1685 | SH3BGRL | O75368 | 0.0061 | 2.214670165 | -0.33 | 18.71 | 18.39 |
| 1686 | SH3BGRL3 | Q9H299 | 0.42 | 0.37675071 | 0.1 | 18.81 | 18.91 |
| 1687 | SH3GL1 | Q99961 | 0.15 | 0.823908741 | -0.08 | 16.41 | 16.33 |
| 1688 | SH3GLB1 | Q9Y371 | 0.34 | 0.468521083 | 0.04 | 16 | 16.04 |
| 1689 | SH3KBP1 | Q96B97 | 0.53 | 0.27572413 | -0.03 | 17.22 | 17.19 |
| 1690 | SH3PXD2B | A1X283 | 0.0065 | 2.187086643 | 0.3 | 16.89 | 17.19 |
| 1691 | SHMT2 | P34897 | 0.039 | 1.408935393 | 0.07 | 15.57 | 15.65 |
| 1692 | SHTN1 | A0MZ66 | 0.1 | 1 | 0.21 | 16.13 | 16.34 |
| 1693 | SKIV2L | Q15477 | 0.46 | 0.337242168 | 0.06 | 14.2 | 14.26 |
| 1694 | SKP1 | P63208 | 0.83 | 0.080921908 | -0.03 | 15.95 | 15.92 |
| 1695 | SLC12A2 | P55011 | 0.7 | 0.15490196 | -0.04 | 16.5 | 16.46 |
| 1696 | SLC16A1 | P53985 | 0.044 | 1.356547324 | -0.24 | 16.79 | 16.55 |

| | | | | | | | |
|---|---|---|---|---|---|---|---|
| 1697 | SLC16A3 | O15427 | 0.046 | 1.337242168 | -0.24 | 18.44 | 18.2 |
| 1698 | SLC25A1 | P53007 | 0.12 | 0.920818754 | 0.09 | 16.17 | 16.26 |
| 1699 | SLC25A11 | Q02978 | 0.6 | 0.22184875 | 0.06 | 15.83 | 15.89 |
| 1700 | SLC25A12 | O75746 | 0.036 | 1.443697499 | 0.19 | 14.7 | 14.89 |
| 1701 | SLC25A13 | Q9UJS0 | 0.041 | 1.387216143 | 0.14 | 15.34 | 15.49 |
| 1702 | SLC25A20 | O43772 | 0.69 | 0.161150909 | 0.05 | 15.12 | 15.17 |
| 1703 | SLC25A24 | Q6NUK1 | 0.017 | 1.769551079 | 0.14 | 15.01 | 15.15 |
| 1704 | SLC25A3 | Q00325 | 0.87 | 0.060480747 | 0 | 17.24 | 17.24 |
| 1705 | SLC25A5 | P05141 | 0.026 | 1.585026652 | 0.15 | 17.91 | 18.07 |
| 1706 | SLC25A6 | P12236 | 0.18 | 0.744727495 | 0.08 | 15.63 | 15.7 |
| 1707 | SLC35B2 | Q8TB61 | 0.21 | 0.677780705 | 0.29 | 16.79 | 17.08 |
| 1708 | SLC3A2 | P08195 | 0.13 | 0.886056648 | 0.1 | 17.02 | 17.12 |
| 1709 | SLC4A7 | Q9Y6M7 | 0.011 | 1.958607315 | 0.24 | 14.97 | 15.22 |
| 1710 | SLFN5 | Q08AF3 | 0.0098 | 2.008773924 | 0.09 | 16.56 | 16.65 |
| 1711 | SLIRP | Q9GZT3 | 0.4 | 0.397940009 | 0.09 | 17.6 | 17.69 |
| 1712 | SLK | Q9H2G2 | 0.11 | 0.958607315 | 0.07 | 16.49 | 16.55 |
| 1713 | SMAP | O00193 | 0.006 | 2.22184875 | -0.21 | 17.22 | 17 |
| 1714 | SMARCA5 | O60264 | 0.63 | 0.200659451 | -0.05 | 14.52 | 14.47 |
| 1715 | SMARCC2 | Q8TAQ2 | 0.57 | 0.244125144 | -0.04 | 17.21 | 17.17 |
| 1716 | SMC1A | Q14683 | 0.027 | 1.568636236 | 0.25 | 15.54 | 15.79 |
| 1717 | SMN1 | Q16637 | 0.64 | 0.193820026 | 0.11 | 15.29 | 15.4 |
| 1718 | SMS | P52788 | 0.05 | 1.301029996 | -0.08 | 15.35 | 15.27 |
| 1719 | SMU1 | Q2TAY7 | 0.18 | 0.744727495 | 0.14 | 16.46 | 16.59 |
| 1720 | SNCA | P37840 | 0.99 | 0.004364805 | -0.01 | 17.76 | 17.75 |
| 1721 | SNCG | O76070 | 0.059 | 1.229147988 | -0.18 | 16.06 | 15.87 |
| 1722 | SND1 | Q7KZF4 | 0.0034 | 2.468521083 | -0.09 | 16.31 | 16.22 |
| 1723 | SNRNP200 | O75643 | 0.026 | 1.585026652 | 0.06 | 15.17 | 15.23 |
| 1724 | SNRNP70 | P08621 | 0.15 | 0.823908741 | -0.09 | 16.91 | 16.82 |
| 1725 | SNRPA | P09012 | 0.23 | 0.638272164 | -0.04 | 18.29 | 18.25 |
| 1726 | SNRPA1 | P09661 | 0.064 | 1.193820026 | -0.14 | 15.95 | 15.81 |
| 1727 | SNRPB | P14678 (+1) | 0.79 | 0.102372909 | 0.04 | 18.42 | 18.46 |
| 1728 | SNRPC | P09234 | 0.94 | 0.026872146 | -0.01 | 16.78 | 16.77 |
| 1729 | SNRPD2 | P62316 | 0.59 | 0.229147988 | 0.06 | 17.26 | 17.32 |
| 1730 | SNRPD3 | P62318 | 0.44 | 0.356547324 | -0.08 | 16.3 | 16.22 |
| 1731 | SNRPE | P62304 | 0.41 | 0.387216143 | 0.04 | 17.62 | 17.66 |
| 1732 | SNRPF | P62306 | 0.19 | 0.721246399 | -0.15 | 16.12 | 15.97 |
| 1733 | SNRPG | P62308 | 0.031 | 1.508638306 | -0.12 | 19.6 | 19.48 |
| 1734 | SNU13 | P55769 | 0.23 | 0.638272164 | 0.1 | 16.63 | 16.73 |
| 1735 | SNW1 | Q13573 | 0.16 | 0.795880017 | -0.15 | 15.91 | 15.76 |
| 1736 | SNX1 | Q13596 | 0.76 | 0.119186408 | -0.03 | 17.03 | 16.99 |
| 1737 | SNX2 | O60749 | 0.61 | 0.214670165 | 0.04 | 16.69 | 16.73 |
| 1738 | SNX3 | O60493 | 0.33 | 0.48148606 | -0.09 | 17.62 | 17.53 |
| 1739 | SNX6 | Q9UNH7 | 0.039 | 1.408935393 | -0.09 | 17.38 | 17.28 |
| 1740 | SNX9 | Q9Y5X1 | 0.91 | 0.040958608 | -0.02 | 15.18 | 15.16 |
| 1741 | SOD1 | P00441 | 0.0001 | 4 | -0.23 | 17.7 | 17.47 |
| 1742 | SOD2 | P04179 | 0.0001 | 4 | -0.94 | 17.45 | 16.51 |
| 1743 | SON | P18583 | 0.0055 | 2.259637311 | -0.1 | 16.23 | 16.12 |
| 1744 | SORBS2 | O94875 | 0.18 | 0.744727495 | 0.24 | 16.63 | 16.86 |
| 1745 | SORD | Q00796 | 0.05 | 1.301029996 | 0.41 | 13.99 | 14.4 |
| 1746 | SP100 | P23497 | 0.12 | 0.920818754 | -0.08 | 15.88 | 15.8 |

| | | | | | | | |
|---|---|---|---|---|---|---|---|
| 1747 | SPAG9 | O60271 | 0.041 | 1.387216143 | 0.35 | 15.02 | 15.37 |
| 1748 | SPARC | P09486 | 0.06 | 1.22184875 | -0.17 | 14.94 | 14.77 |
| 1749 | SPART | Q8N0X7 | 0.25 | 0.602059991 | -0.18 | 16.32 | 16.14 |
| 1750 | SPECC1L | Q69YQ0 | 0.19 | 0.721246399 | -0.09 | 17.88 | 17.79 |
| 1751 | SPR | P35270 | 0.00016 | 3.795880017 | 0.48 | 14.81 | 15.29 |
| 1752 | SPTAN1 | Q13813 | 0.0001 | 4 -0.09 | 15.83 | 15.74 | |
| 1753 | SPTBN1 | Q01082 | 0.03 | 1.522878745 | -0.04 | 16.5 | 16.46 |
| 1754 | SPTLC1 | O15269 | 0.32 | 0.494850022 | 0.19 | 15.63 | 15.82 |
| 1755 | SQOR | Q9Y6N5 | 0.9 | 0.045757491 | 0.02 | 16.07 | 16.09 |
| 1756 | SQSTM1 | Q13501 | 0.0065 | 2.187086643 | -0.33 | 16.15 | 15.82 |
| 1757 | SRC | P12931 | 0.0092 | 2.036212173 | 0.51 | 14.91 | 15.42 |
| 1758 | SRGAP2 | O75044 | 0.32 | 0.494850022 | -0.08 | 14.73 | 14.65 |
| 1759 | SRI | P30626 | 0.0012 | 2.920818754 | 0.11 | 17.58 | 17.69 |
| 1760 | SRM | P19623 | 0.59 | 0.229147988 | 0.03 | 16.66 | 16.69 |
| 1761 | SRP14 | P37108 | 0.041 | 1.387216143 | 0.25 | 16.94 | 17.19 |
| 1762 | SRP19 | P09132 | 0.78 | 0.107905397 | 0.03 | 15.45 | 15.48 |
| 1763 | SRP54 | P61011 | 0.98 | 0.008773924 | 0 | 16.9 | 16.9 |
| 1764 | SRP68 | Q9UHB9 | 0.72 | 0.142667504 | -0.04 | 16.56 | 16.53 |
| 1765 | SRP72 | O76094 | 0.88 | 0.055517328 | -0.04 | 16.59 | 16.55 |
| 1766 | SRPRA | P08240 | 0.11 | 0.958607315 | 0.11 | 15.06 | 15.17 |
| 1767 | SRPRB | Q9Y5M8 | 0.2 | 0.698970004 | 0.2 | 17.2 | 17.4 |
| 1768 | SRRM1 | Q8IYB3 | 0.76 | 0.119186408 | 0.01 | 15.09 | 15.1 |
| 1769 | SRRM2 | Q9UQ35 | 0.91 | 0.040958608 | 0.01 | 16.57 | 16.58 |
| 1770 | SRRT | Q9BXP5 | 0.6 | 0.22184875 | 0.06 | 14.41 | 14.47 |
| 1771 | SRSF1 | Q07955 | 0.00097 | 3.013228266 | -0.16 | 17.38 | 17.22 |
| 1772 | SRSF2 | Q01130 | 0.15 | 0.823908741 | -0.07 | 16.31 | 16.24 |
| 1773 | SRSF3 | P84103 | 0.77 | 0.113509275 | -0.02 | 18.27 | 18.26 |
| 1774 | SRSF5 | Q13243 | 0.05 | 1.301029996 | -0.16 | 15.27 | 15.11 |
| 1775 | SRSF6 | Q13247 | 0.71 | 0.148741651 | 0.06 | 18.93 | 18.99 |
| 1776 | SRSF7 | Q16629 | 0.58 | 0.236572006 | -0.09 | 18.56 | 18.47 |
| 1777 | SRSF9 | Q13242 | 0.97 | 0.013228266 | -0.01 | 16.04 | 16.03 |
| 1778 | SSB | P05455 | 0.013 | 1.886056648 | 0.08 | 15.89 | 15.97 |
| 1779 | SSBP1 | Q04837 | 0.37 | 0.431798276 | 0.15 | 15.58 | 15.73 |
| 1780 | SSR1 | P43307 | 0.35 | 0.455931956 | -0.06 | 17.52 | 17.45 |
| 1781 | SSR4 | P51571 | 0.093 | 1.031517051 | -0.21 | 18.75 | 18.55 |
| 1782 | SSRP1 | Q08945 | 0.076 | 1.119186408 | 0.17 | 16.17 | 16.34 |
| 1783 | ST13 | P50502 | 0.93 | 0.031517051 | -0.01 | 18.23 | 18.22 |
| 1784 | STAB1 | Q9NY15 | 0.8 | 0.096910013 | 0.06 | 14.93 | 14.99 |
| 1785 | STAT1 | P42224 | 0.00061 | 3.214670165 | -0.19 | 17.12 | 16.93 |
| 1786 | STAT3 | P40763 | 0.22 | 0.657577319 | 0.07 | 15.4 | 15.47 |
| 1787 | STAU1 | O95793 | 0.033 | 1.48148606 | 0.16 | 16.68 | 16.84 |
| 1788 | STIM1 | Q13586 | 0.06 | 1.22184875 | 0.8 | 15.58 | 16.38 |
| 1789 | STIP1 | P31948 | 0.00048 | 3.318758763 | -0.15 | 17.47 | 17.32 |
| 1790 | STK10 | O94804 | 0.76 | 0.119186408 | 0.07 | 15.38 | 15.45 |
| 1791 | STK24 | Q9Y6E0 | 0.12 | 0.920818754 | 0.14 | 14.86 | 15 |
| 1792 | STMN1 | P16949 | 0.0035 | 2.455931956 | -0.23 | 17.4 | 17.17 |
| 1793 | STOM | P27105 | 0.79 | 0.102372909 | 0.02 | 15.98 | 15.99 |
| 1794 | STOML2 | Q9UJZ1 | 0.12 | 0.920818754 | -0.07 | 15.84 | 15.77 |
| 1795 | STRAP | Q9Y3F4 | 0.49 | 0.30980392 | -0.03 | 16.24 | 16.21 |
| 1796 | STT3A | P46977 | 0.041 | 1.387216143 | 0.16 | 16.56 | 16.72 |

| | | | | | | | |
|---|---|---|---|---|---|---|---|
| 1797 | STT3B | Q8TCJ2 | 0.019 | 1.721246399 | 0.29 | 16.09 | 16.38 |
| 1798 | STX12 | Q86Y82 | 0.008 | 2.096910013 | 0.37 | 16.28 | 16.65 |
| 1799 | STXBP1 | P61764 | 0.71 | 0.148741651 | 0.06 | 15.72 | 15.78 |
| 1800 | STXBP3 | O00186 | 0.25 | 0.602059991 | 0.19 | 15.58 | 15.77 |
| 1801 | SUB1 | P53999 | 0.014 | 1.853871964 | -0.15 | 17.41 | 17.26 |
| 1802 | SUCLA2 | Q9P2R7 | 0.14 | 0.853871964 | 0.29 | 16.41 | 16.69 |
| 1803 | SUCLG1 | P53597 | 0.0001 | 4 | 0.3 | 17.05 | 17.35 |
| 1804 | SUCLG2 | Q96I99 | 0.0001 | 4 | 0.17 | 16.79 | 16.96 |
| 1805 | SUGT1 | Q9Y2Z0 | 0.35 | 0.455931956 | -0.08 | 16.63 | 16.55 |
| 1806 | SULT1B1 | O43704 | 0.69 | 0.161150909 | 0.06 | 15.55 | 15.61 |
| 1807 | SUMF2 | Q8NBJ7 | NA | NA | 0.04 | 15.89 | 15.93 |
| 1808 | SUMO2 | P61956 | 0.15 | 0.823908741 | -0.21 | 18.8 | 18.6 |
| 1809 | SUN2 | Q9UH99 | 0.077 | 1.113509275 | -0.1 | 15.22 | 15.12 |
| 1810 | SUPT16H | Q9Y5B9 | 0.048 | 1.318758763 | 0.27 | 15 | 15.27 |
| 1811 | SUPT6H | Q7KZ85 | 0.69 | 0.161150909 | 0.09 | 11.97 | 12.06 |
| 1812 | SURF4 | O15260 | 0.073 | 1.13667714 | 0.34 | 15.24 | 15.58 |
| 1813 | SVIL | O95425 | 0.24 | 0.619788758 | 0.15 | 14.84 | 15 |
| 1814 | SWAP70 | Q9UH65 | 0.59 | 0.229147988 | -0.06 | 14 | 13.93 |
| 1815 | SYMPK | Q92797 | 0.6 | 0.22184875 | -0.07 | 14.87 | 14.8 |
| 1816 | SYNCRIP | O60506 | 0.0001 | 4 | -0.11 | 16.05 | 15.94 |
| 1817 | SYNPO | Q8N3V7 | 0.16 | 0.795880017 | -0.08 | 15.95 | 15.87 |
| 1818 | TACC1 | O75410 | 0.67 | 0.173925197 | 0.08 | 14.94 | 15.02 |
| 1819 | TACC2 | O95359 | 0.12 | 0.920818754 | 0.18 | 15.2 | 15.39 |
| 1820 | TACO1 | Q9BSH4 | 0.8 | 0.096910013 | -0.06 | 14.88 | 14.83 |
| 1821 | TAF15 | Q92804 | 0.7 | 0.15490196 | 0.11 | 15.18 | 15.28 |
| 1822 | TAGLN | Q01995 | 0.0016 | 2.795880017 | 1.03 | 15.22 | 16.25 |
| 1823 | TAGLN2 | P37802 | 0.0014 | 2.853871964 | 0.24 | 16.35 | 16.6 |
| 1824 | TALDO1 | P37837 | 0.12 | 0.920818754 | -0.08 | 17.85 | 17.77 |
| 1825 | TAP1 | Q03518 | 0.36 | 0.443697499 | 0.2 | 15.2 | 15.39 |
| 1826 | TAPBP | O15533 | 0.19 | 0.721246399 | 0.34 | 17.35 | 17.69 |
| 1827 | TARDBP | Q13148 | 0.011 | 1.958607315 | 0.14 | 15.26 | 15.41 |
| 1828 | TARS | P26639 | 0.64 | 0.193820026 | 0.04 | 15.63 | 15.67 |
| 1829 | TBCB | Q99426 | 0.99 | 0.004364805 | 0 | 17.17 | 17.16 |
| 1830 | TBCD | Q9BTW9 | 0.2 | 0.698970004 | 0.13 | 14.14 | 14.27 |
| 1831 | TCOF1 | Q13428 | 0.77 | 0.113509275 | 0.01 | 18.18 | 18.19 |
| 1832 | TCP1 | P17987 | 0.0001 | 4 | 0.15 | 17.05 | 17.19 |
| 1833 | TECR | Q9NZ01 | 0.61 | 0.214670165 | 0.06 | 18.53 | 18.6 |
| 1834 | TES | Q9UGI8 | 0.48 | 0.318758763 | 0.03 | 17.08 | 17.11 |
| 1835 | TFG | Q92734 | 0.51 | 0.292429824 | -0.04 | 16.29 | 16.25 |
| 1836 | TFRC | P02786 | 0.082 | 1.086186148 | -0.12 | 15.96 | 15.83 |
| 1837 | TGFB1I1 | O43294 | 0.18 | 0.744727495 | 0.24 | 16.06 | 16.3 |
| 1838 | TGM2 | P21980 | 0.0001 | 4 | -0.24 | 16.49 | 16.25 |
| 1839 | TGOLN2 | O43493 | 0.95 | 0.022276395 | -0.01 | 18.55 | 18.54 |
| 1840 | THBS1 | P07996 | 0.0001 | 4 | -0.17 | 16.6 | 16.43 |
| 1841 | THOP1 | P52888 | 0.18 | 0.744727495 | 0.16 | 15.71 | 15.86 |
| 1842 | THRAP3 | Q9Y2W1 | 0.12 | 0.920818754 | -0.11 | 17.42 | 17.31 |
| 1843 | THUMPD1 | Q9NXG2 | 0.22 | 0.657577319 | 0.08 | 17.28 | 17.36 |
| 1844 | TIAL1 | Q01085 | 0.86 | 0.065501549 | 0.01 | 17.02 | 17.03 |
| 1845 | TIGAR | Q9NQ88 | 0.074 | 1.13076828 | 0.26 | 16.96 | 17.22 |
| 1846 | TIMM44 | O43615 | 0.97 | 0.013228266 | 0 | 16.3 | 16.3 |

| | | | | | | | |
|---|---|---|---|---|---|---|---|
| 1847 | TIMM50 | Q3ZCQ8 | 0.64 | 0.193820026 | 0.05 | 16.89 | 16.94 |
| 1848 | TIPRL | O75663 | 0.47 | 0.327902142 | -0.07 | 15.61 | 15.54 |
| 1849 | TJP1 | Q07157 | 0.014 | 1.853871964 | 0.15 | 16.38 | 16.52 |
| 1850 | TJP2 | Q9UDY2 | 0.83 | 0.080921908 | 0 | 16.38 | 16.38 |
| 1851 | TKT | P29401 | 0.97 | 0.013228266 | 0 | 16.8 | 16.81 |
| 1852 | TLN1 | Q9Y490 | 0.0001 | 4 | 0.37 | 16.03 | 16.4 |
| 1853 | TLN2 | Q9Y4G6 | NA | NA | 0.52 | 17.89 | 18.42 |
| 1854 | TM9SF2 | Q99805 | 0.15 | 0.823908741 | 0.17 | 15.64 | 15.8 |
| 1855 | TM9SF3 | Q9HD45 | 0.4 | 0.397940009 | 0.15 | 15.85 | 16 |
| 1856 | TMED10 | P49755 | 0.37 | 0.431798276 | -0.04 | 16.22 | 16.18 |
| 1857 | TMED2 | Q15363 | 0.87 | 0.060480747 | 0.01 | 17.15 | 17.16 |
| 1858 | TMED4 | Q7Z7H5 | 0.27 | 0.568636236 | 0.12 | 18.06 | 18.18 |
| 1859 | TMED5 | Q9Y3A6 | 0.51 | 0.292429824 | 0.08 | 14.83 | 14.91 |
| 1860 | TMED7 | Q9Y3B3 | 0.015 | 1.823908741 | 0.11 | 17.51 | 17.61 |
| 1861 | TMED9 | Q9BVK6 | 0.66 | 0.180456064 | 0.08 | 17.18 | 17.26 |
| 1862 | TMEM109 | Q9BVC6 | 0.15 | 0.823908741 | -0.17 | 20.21 | 20.04 |
| 1863 | TMEM165 | Q9HC07 | 0.056 | 1.251811973 | -0.18 | 15.04 | 14.86 |
| 1864 | TMEM214 | Q6NUQ4 | 0.32 | 0.494850022 | 0.13 | 17.34 | 17.47 |
| 1865 | TMEM263 | Q8WUH6 | 0.026 | 1.585026652 | 0.29 | 16.53 | 16.82 |
| 1866 | TMEM33 | P57088 | 0.18 | 0.744727495 | 0.32 | 15.66 | 15.97 |
| 1867 | TMEM43 | Q9BTV4 | 0.2 | 0.698970004 | 0.12 | 16.16 | 16.27 |
| 1868 | TMOD3 | Q9NYL9 | 0.83 | 0.080921908 | 0.01 | 15.68 | 15.69 |
| 1869 | TMPO | P42167 | 0.11 | 0.958607315 | -0.13 | 15.26 | 15.13 |
| 1870 | TMSB10 | P63313 | NA | NA | -0.5 | 18.94 | 18.44 |
| 1871 | TMSB4X | P62328 | NA | NA | -0.46 | 18.53 | 18.07 |
| 1872 | TMX1 | Q9H3N1 | 0.5 | 0.301029996 | 0.04 | 17.24 | 17.28 |
| 1873 | TMX3 | Q96JJ7 | 0.024 | 1.619788758 | 0.28 | 16.07 | 16.35 |
| 1874 | TNKS1BP1 | Q9C0C2 | 0.74 | 0.13076828 | 0.02 | 16.02 | 16.04 |
| 1875 | TNPO1 | Q92973 | 0.73 | 0.13667714 | 0.02 | 16.17 | 16.19 |
| 1876 | TNPO3 | Q9Y5L0 | 0.32 | 0.494850022 | 0.19 | 14.15 | 14.34 |
| 1877 | TOMM22 | Q9NS69 | 0.038 | 1.420216403 | 0.31 | 14.71 | 15.01 |
| 1878 | TOMM40 | O96008 | 0.34 | 0.468521083 | -0.08 | 16.11 | 16.03 |
| 1879 | TOMM70 | O94826 | 0.43 | 0.366531544 | 0.06 | 16.05 | 16.12 |
| 1880 | TOP1 | P11387 | 0.1 | 1 | 0.09 | 17.65 | 17.74 |
| 1881 | TOP2B | Q02880 | 0.21 | 0.677780705 | 0.11 | 15.84 | 15.95 |
| 1882 | TOR1A | O14656 | 0.23 | 0.638272164 | 0.17 | 16 | 16.17 |
| 1883 | TOR1AIP1 | Q5JTV8 | 0.48 | 0.318758763 | 0.04 | 17.1 | 17.14 |
| 1884 | TOR1AIP2 | Q8NFQ8 | 0.15 | 0.823908741 | 0.1 | 17.2 | 17.31 |
| 1885 | TP53BP1 | Q12888 | 0.97 | 0.013228266 | 0 | 16.38 | 16.38 |
| 1886 | TP53I3 | Q53FA7 | 0.43 | 0.366531544 | 0.08 | 17.42 | 17.5 |
| 1887 | TPD52L2 | O43399 | 0.15 | 0.823908741 | -0.16 | 16.82 | 16.67 |
| 1888 | TPI1 | P60174 | 0.69 | 0.161150909 | -0.01 | 17.31 | 17.3 |
| 1889 | TPM3 | P06753 | 0.099 | 1.004364805 | -0.2 | 17.21 | 17 |
| 1890 | TPM4 | P67936 | 0.61 | 0.214670165 | -0.02 | 16.73 | 16.72 |
| 1891 | TPP1 | O14773 | 0.65 | 0.187086643 | -0.05 | 16.6 | 16.55 |
| 1892 | TPP2 | P29144 | 0.67 | 0.173925197 | -0.04 | 16.92 | 16.87 |
| 1893 | TPR | P12270 | 0.67 | 0.173925197 | -0.01 | 16.82 | 16.8 |
| 1894 | TPT1 | P13693 | 0.61 | 0.214670165 | -0.03 | 16.68 | 16.64 |
| 1895 | TRA2B | P62995 | 0.57 | 0.244125144 | -0.03 | 17.31 | 17.28 |
| 1896 | TRAP1 | Q12931 | 0.64 | 0.193820026 | -0.02 | 17.13 | 17.1 |

| | | | | | | | | | |
|---|---|---|---|---|---|---|---|---|---|
| 1897 | TRIM21 | P19474 | 0.46 | 0.337242168 | | 0.11 | 15.67 | 15.78 | |
| 1898 | TRIM25 | Q14258 | 0.14 | 0.853871964 | | 0.11 | 16.68 | 16.79 | |
| 1899 | TRIM28 | Q13263 | 0.0074 | 2.13076828 | | 0.09 | 16.46 | 16.56 | |
| 1900 | TRIM47 | Q96LD4 | 0.056 | 1.251811973 | | 0.19 | 15.19 | 15.38 | |
| 1901 | TRIP10 | Q15642 | 0.082 | 1.086186148 | | -0.14 | 15.74 | 15.6 | |
| 1902 | TRIP12 | Q14669 | 0.47 | 0.327902142 | | -0.19 | 14.14 | 13.96 | |
| 1903 | TRIR | Q9BQ61 | 0.36 | 0.443697499 | | -0.1 | 17.82 | 17.72 | |
| 1904 | TRMT10C | Q7L0Y3 | 0.69 | 0.161150909 | | 0.06 | 16.07 | 16.13 | |
| 1905 | TRPV2 | Q9Y5S1 | 0.64 | 0.193820026 | | 0.04 | 14.74 | 14.77 | |
| 1906 | TSC22D4 | Q9Y3Q8 | 0.42 | 0.37675071 | | 0.11 | 16.99 | 17.1 | |
| 1907 | TSFM | P43897 | 0.51 | 0.292429824 | | 0.04 | 14.92 | 14.96 | |
| 1908 | TSN | Q15631 | 0.4 | 0.397940009 | | 0.12 | 14.84 | 14.96 | |
| 1909 | TSNAX | Q99598 | 0.34 | 0.468521083 | | 0.06 | 14.05 | 14.12 | |
| 1910 | TSPO | P30536 | 0.16 | 0.795880017 | | 0.12 | 15.37 | 15.48 | |
| 1911 | TSR1 | Q2NL82 | 0.75 | 0.124938737 | | 0.01 | 13.88 | 13.89 | |
| 1912 | TST | Q16762 | 0.68 | 0.167491087 | | 0.03 | 15.32 | 15.35 | |
| 1913 | TTC37 | Q6PGP7 | 0.76 | 0.119186408 | | -0.02 | 16.35 | 16.33 | |
| 1914 | TTLL12 | Q14166 | 0.59 | 0.229147988 | | 0.02 | 16.69 | 16.71 | |
| 1915 | TUBA1B | P68363 | 0.17 | 0.769551079 | | 0.04 | 16.07 | 16.1 | |
| 1916 | TUBA1C | Q9BQE3 | 0.059 | 1.229147988 | | 0.19 | 16.34 | 16.53 | |
| 1917 | TUBB | P07437 | 0.33 | 0.48148606 | | 0.07 | 16.77 | 16.84 | |
| 1918 | TUBB2B | Q9BVA1 | 0.68 | 0.167491087 | | -0.03 | 15.8 | 15.78 | |
| 1919 | TUBB3 | Q13509 | 0.012 | 1.920818754 | | 0.66 | 13.55 | 14.21 | |
| 1920 | TUBB4A | P04350 | NA | NA | -0.12 | 15.91 | 15.79 | | |
| 1921 | TUBB4B | P68371 | NA | NA | -0.04 | 17.86 | 17.82 | | |
| 1922 | TUBB6 | Q9BUF5 | 0.0033 | 2.48148606 | | 0.18 | 16.05 | 16.23 | |
| 1923 | TUBG1 | P23258 (+1) | | 0.44 | 0.356547324 | | 0.17 | 14.7 | 14.87 |
| 1924 | TUFM | P49411 | 0.074 | 1.13076828 | | 0.04 | 16.71 | 16.75 | |
| 1925 | TWF1 | Q12792 | 0.28 | 0.552841969 | | -0.03 | 15.75 | 15.72 | |
| 1926 | TWF2 | Q6IBS0 | 0.9 | 0.045757491 | | 0 | 15.26 | 15.26 | |
| 1927 | TXN | P10599 | 0.14 | 0.853871964 | | 0.25 | 17.11 | 17.36 | |
| 1928 | TXNDC12 | O95881 | 0.34 | 0.468521083 | | 0.04 | 16.72 | 16.76 | |
| 1929 | TXNDC17 | Q9BRA2 | 0.24 | 0.619788758 | | 0.12 | 16.91 | 17.03 | |
| 1930 | TXNDC5 | Q8NBS9 | 0.0001 | 4 | -0.21 | 17.68 | 17.47 | | |
| 1931 | TXNDC9 | O14530 | 0.092 | 1.036212173 | | 0.19 | 16.3 | 16.48 | |
| 1932 | TXNL1 | O43396 | 0.32 | 0.494850022 | | -0.03 | 16.21 | 16.17 | |
| 1933 | TXNRD1 | Q16881 | 0.0001 | 4 | -0.18 | 16.04 | 15.86 | | |
| 1934 | TXNRD2 | Q9NNW7 | 0.9 | 0.045757491 | | -0.05 | 16.74 | 16.69 | |
| 1935 | U2AF1L5 | P0DN76 (+1) | | 0.63 | 0.200659451 | | 0.04 | 15.41 | 15.45 |
| 1936 | U2AF2 | P26368 | 0.02 | 1.698970004 | | 0.14 | 16.02 | 16.17 | |
| 1937 | U2SURP | O15042 | 0.45 | 0.346787486 | | -0.07 | 15.36 | 15.3 | |
| 1938 | UAP1 | Q16222 | 0.0016 | 2.795880017 | | 0.26 | 16.2 | 16.46 | |
| 1939 | UAP1L1 | Q3KQV9 | 0.21 | 0.677780705 | | 0.11 | 15.73 | 15.84 | |
| 1940 | UBA1 | P22314 | 0.0014 | 2.853871964 | | 0.09 | 16.36 | 16.45 | |
| 1941 | UBA2 | Q9UBT2 | 0.63 | 0.200659451 | | -0.02 | 16.71 | 16.68 | |
| 1942 | UBA3 | Q8TBC4 | 0.11 | 0.958607315 | | 0.16 | 15.28 | 15.44 | |
| 1943 | UBA6 | A0AVT1 | 0.022 | 1.657577319 | | 0.15 | 15.94 | 16.1 | |
| 1944 | UBAP2 | Q5T6F2 | 0.81 | 0.091514981 | | 0.06 | 14.82 | 14.89 | |
| 1945 | UBAP2L | Q14157 | 0.71 | 0.148741651 | | -0.02 | 16.44 | 16.42 | |
| 1946 | UBE2D2 | P62837 | 0.99 | 0.004364805 | | 0 | 16.92 | 16.92 | |

| | | | | | | | |
|---|---|---|---|---|---|---|---|
| 1947 | UBE2I | P63279 | 0.76 | 0.119186408 | -0.04 | 14.83 | 14.79 |
| 1948 | UBE2K | P61086 | 0.97 | 0.013228266 | 0.01 | 15.49 | 15.5 |
| 1949 | UBE2L3 | P68036 | 0.21 | 0.677780705 | -0.05 | 16.64 | 16.6 |
| 1950 | UBE2M | P61081 | 0.027 | 1.568636236 | -0.15 | 16.45 | 16.3 |
| 1951 | UBE2N | P61088 | 0.14 | 0.853871964 | -0.05 | 15.49 | 15.44 |
| 1952 | UBE2O | sp\|Q9C0C9\|UBE2O_HUMAN | 0.59 | 0.229147988 | -0.07 | 14.92 | 14.85 |
| 1953 | UBE2V1 | Q13404 | 0.53 | 0.27572413 | -0.02 | 15.56 | 15.53 |
| 1954 | UBE4A | Q14139 | 0.047 | 1.327902142 | 0.44 | 14.25 | 14.69 |
| 1955 | UBQLN1 | Q9UMX0 | 0.036 | 1.443697499 | 0.1 | 14.38 | 14.47 |
| 1956 | UBQLN4 | Q9NRR5 | 0.43 | 0.366531544 | 0.07 | 13.65 | 13.71 |
| 1957 | UBR4 | Q5T4S7 | 0.0013 | 2.886056648 | 0.19 | 15 | 15.19 |
| 1958 | UBXN4 | Q92575 | 0.26 | 0.585026652 | 0.12 | 15.35 | 15.47 |
| 1959 | UCHL1 | P09936 | 0.59 | 0.229147988 | 0.02 | 18.08 | 18.11 |
| 1960 | UCHL3 | P15374 | 0.28 | 0.552841969 | 0.12 | 16.17 | 16.29 |
| 1961 | UFL1 | O94874 | 0.6 | 0.22184875 | 0.06 | 15.68 | 15.74 |
| 1962 | UFM1 | P61960 | 0.02 | 1.698970004 | -0.1 | 15.21 | 15.1 |
| 1963 | UFSP2 | Q9NUQ7 | 0.19 | 0.721246399 | 0.25 | 14.4 | 14.65 |
| 1964 | UGDH | O60701 | 0.3 | 0.522878745 | -0.08 | 15.73 | 15.65 |
| 1965 | UGGT1 | Q9NYU2 | 0.72 | 0.142667504 | 0.01 | 16.24 | 16.25 |
| 1966 | UGP2 | Q16851 | 0.0088 | 2.055517328 | 0.25 | 17.03 | 17.28 |
| 1967 | UMPS | P11172 | 0.85 | 0.070581074 | 0.02 | 15.58 | 15.6 |
| 1968 | UNC45A | Q9H3U1 | 0.61 | 0.214670165 | 0.01 | 15.55 | 15.55 |
| 1969 | UPF1 | Q92900 | 0.42 | 0.37675071 | 0.07 | 16.43 | 16.5 |
| 1970 | UPF2 | Q9HAU5 | 0.53 | 0.27572413 | -0.13 | 14.73 | 14.61 |
| 1971 | UPP1 | Q16831 | 0.38 | 0.420216403 | 0.18 | 17.4 | 17.58 |
| 1972 | UQCRC1 | P31930 | 0.093 | 1.031517051 | 0.06 | 16.62 | 16.68 |
| 1973 | UQCRC2 | P22695 | 0.99 | 0.004364805 | 0 | 18.01 | 18.01 |
| 1974 | UQCRFS1 | P47985 | 0.021 | 1.677780705 | -0.15 | 15.82 | 15.67 |
| 1975 | UROD | P06132 | 0.47 | 0.327902142 | -0.06 | 15.7 | 15.64 |
| 1976 | USO1 | O60763 | 0.21 | 0.677780705 | 0.08 | 15.88 | 15.96 |
| 1977 | USP10 | Q14694 | 0.07 | 1.15490196 | -0.39 | 12.71 | 12.32 |
| 1978 | USP14 | P54578 | 0.037 | 1.431798276 | 0.13 | 16.26 | 16.39 |
| 1979 | USP15 | Q9Y4E8 | 0.018 | 1.744727495 | 0.59 | 13.74 | 14.32 |
| 1980 | USP39 | Q53GS9 | 0.93 | 0.031517051 | -0.08 | 13.92 | 13.84 |
| 1981 | USP5 | P45974 | 0.88 | 0.055517328 | -0.01 | 16.44 | 16.44 |
| 1982 | USP9X | Q93008 | 0.92 | 0.036212173 | 0 | 17.13 | 17.12 |
| 1983 | UTP20 | O75691 | 0.55 | 0.259637311 | 0.17 | 13.55 | 13.72 |
| 1984 | UTRN | P46939 | 0.8 | 0.096910013 | 0.04 | 14.49 | 14.52 |
| 1985 | VAC14 | Q08AM6 | 0.54 | 0.26760624 | 0.05 | 15.89 | 15.94 |
| 1986 | VAPA | Q9P0L0 | 0.028 | 1.552841969 | -0.14 | 16.97 | 16.83 |
| 1987 | VAPB | O95292 | 0.28 | 0.552841969 | 0.1 | 18.24 | 18.34 |
| 1988 | VARS | P26640 | 0.9 | 0.045757491 | -0.02 | 16.91 | 16.89 |
| 1989 | VASP | P50552 | 0.24 | 0.619788758 | 0.06 | 15.96 | 16.02 |
| 1990 | VAT1 | Q99536 | 0.12 | 0.920818754 | 0.08 | 16.31 | 16.38 |
| 1991 | VBP1 | P61758 | 0.77 | 0.113509275 | 0.02 | 16.02 | 16.04 |
| 1992 | VCL | P18206 | 0.015 | 1.823908741 | 0.06 | 16.91 | 16.97 |
| 1993 | VCP | P55072 | 0.0014 | 2.853871964 | -0.11 | 16.46 | 16.35 |
| 1994 | VDAC1 | P21796 | 0.7 | 0.15490196 | 0.01 | 17.37 | 17.38 |
| 1995 | VDAC2 | P45880 | 0.016 | 1.795880017 | 0.1 | 17.58 | 17.68 |

| | | | | | | | | | |
|---|---|---|---|---|---|---|---|---|---|
| 1996 | VDAC3 | Q9Y277 | 0.26 | 0.585026652 | | 0.05 | 17.03 | 17.07 | |
| 1997 | VIM | P08670 | 0.0076 | 2.119186408 | | -0.09 | 17.34 | 17.25 | |
| 1998 | VPS16 | Q9H269 | 0.061 | 1.214670165 | | 0.18 | 13.37 | 13.55 | |
| 1999 | VPS26A | O75436 | 0.027 | 1.568636236 | | -0.08 | 18.18 | 18.1 | |
| 2000 | VPS29 | Q9UBQ0 | 0.85 | 0.070581074 | | 0.02 | 17.46 | 17.48 | |
| 2001 | VPS35 | Q96QK1 | 0.00017 | 3.769551079 | | 0.32 | 15.51 | 15.83 | |
| 2002 | VPS35L | Q7Z3J2 | NA | NA | 0.32 | 12.66 | 12.99 | | |
| 2003 | VPS37A | Q8NEZ2 | 0.72 | 0.142667504 | | 0.06 | 13.56 | 13.61 | |
| 2004 | VRK2 | Q86Y07 | 0.24 | 0.619788758 | | 0.31 | 15.03 | 15.33 | |
| 2005 | VTA1 | Q9NP79 | 0.83 | 0.080921908 | | -0.08 | 17.04 | 16.96 | |
| 2006 | VWF | P04275 | 0.13 | 0.886056648 | | 0.03 | 16.06 | 16.09 | |
| 2007 | WARS | P23381 | 0.26 | 0.585026652 | | -0.07 | 16.15 | 16.07 | |
| 2008 | WASHC1 | A8K0Z3 (+4) | | 0.47 | 0.327902142 | | 0.1 | 14.78 | 14.89 |
| 2009 | WASHC2A | Q641Q2 | 0.038 | 1.420216403 | | -0.15 | 14.92 | 14.77 | |
| 2010 | WASHC4 | Q2M389 | 0.24 | 0.619788758 | | 0.14 | 15.51 | 15.65 | |
| 2011 | WASL | O00401 | 0.58 | 0.236572006 | | 0.07 | 13.92 | 13.99 | |
| 2012 | WBP11 | Q9Y2W2 | 0.48 | 0.318758763 | | -0.18 | 14.83 | 14.65 | |
| 2013 | WDR1 | O75083 | 0.0001 | 4 | 0.12 | 17.98 | 18.1 | | |
| 2014 | WDR11 | Q9BZH6 | 0.46 | 0.337242168 | | -0.12 | 18.99 | 18.87 | |
| 2015 | WDR18 | Q9BV38 | 0.16 | 0.795880017 | | -0.06 | 17.88 | 17.82 | |
| 2016 | WDR43 | Q15061 | 0.0016 | 2.795880017 | | 0.24 | 15.67 | 15.91 | |
| 2017 | WDR61 | Q9GZS3 | 0.064 | 1.193820026 | | -0.12 | 16.29 | 16.18 | |
| 2018 | WDR75 | Q8IWA0 | 0.1 | 1 | 0.26 | 16.61 | 16.87 | | |
| 2019 | WFS1 | O76024 | 0.013 | 1.886056648 | | -0.44 | 16.59 | 16.16 | |
| 2020 | WWTR1 | Q9GZV5 | 0.79 | 0.102372909 | | 0.04 | 16.31 | 16.35 | |
| 2021 | XPNPEP1 | Q9NQW7 | 0.94 | 0.026872146 | | 0 | 16.76 | 16.76 | |
| 2022 | XPO1 | O14980 | 0.24 | 0.619788758 | | 0.05 | 15.78 | 15.83 | |
| 2023 | XPO5 | Q9HAV4 | 0.43 | 0.366531544 | | 0.11 | 15.68 | 15.79 | |
| 2024 | XPO7 | Q9UIA9 | 0.22 | 0.657577319 | | 0.23 | 13.41 | 13.64 | |
| 2025 | XPOT | O43592 | 0.23 | 0.638272164 | | 0.12 | 14.58 | 14.7 | |
| 2026 | XRCC5 | P13010 | 0.0001 | 4 | 0.18 | 15.89 | 16.06 | | |
| 2027 | XRCC6 | P12956 | 0.21 | 0.677780705 | | 0.03 | 16.47 | 16.5 | |
| 2028 | XRN2 | Q9H0D6 | 0.86 | 0.065501549 | | 0.01 | 15.93 | 15.94 | |
| 2029 | YARS | P54577 | 0.4 | 0.397940009 | | 0.03 | 16.34 | 16.38 | |
| 2030 | YBX1 | P67809 | 0.0007 | 3.15490196 | | -0.29 | 16.29 | 16 | |
| 2031 | YBX3 | P16989 | 0.061 | 1.214670165 | | -0.05 | 16.31 | 16.26 | |
| 2032 | YES1 | P07947 | 0.0011 | 2.958607315 | | 0.55 | 15.49 | 16.04 | |
| 2033 | YKT6 | O15498 | 0.66 | 0.180456064 | | 0.07 | 15.1 | 15.16 | |
| 2034 | YTHDF2 | Q9Y5A9 | 0.38 | 0.420216403 | | -0.22 | 16.91 | 16.69 | |
| 2035 | YWHAB | P31946 | 0.72 | 0.142667504 | | -0.01 | 16.03 | 16.02 | |
| 2036 | YWHAE | P62258 | 0.017 | 1.769551079 | | -0.07 | 16.49 | 16.42 | |
| 2037 | YWHAG | P61981 | 0.0049 | 2.30980392 | | -0.19 | 16.23 | 16.04 | |
| 2038 | YWHAH | Q04917 | 0.04 | 1.397940009 | | 0.19 | 15.28 | 15.47 | |
| 2039 | YWHAQ | P27348 | 0.81 | 0.091514981 | | -0.02 | 15.62 | 15.61 | |
| 2040 | YWHAZ | P63104 | 0.1 | 1 | -0.1 | 16.95 | 16.84 | | |
| 2041 | ZC3H15 | Q8WU90 | 0.5 | 0.301029996 | | -0.07 | 17.25 | 17.18 | |
| 2042 | ZC3H18 | Q86VM9 | 0.72 | 0.142667504 | | -0.04 | 16.62 | 16.58 | |
| 2043 | ZC3HAV1 | Q7Z2W4 | 0.32 | 0.494850022 | | 0.13 | 15.77 | 15.9 | |
| 2044 | ZC3HAV1L | | Q96H79 | 0.0094 | 2.026872146 | | 0.58 | 14.01 | 14.59 |
| 2045 | ZC3HC1 | Q86WB0 | 0.48 | 0.318758763 | | 0.14 | 16.12 | 16.25 | |

| | | | | | | | | |
|---|---|---|---|---|---|---|---|---|
| 2046 | ZMPSTE24 | | O75844 | 0.95 | 0.022276395 | 0 | 18.7 | 18.7 |
| 2047 | ZNF185 | O15231 | 0.0066 | 2.180456064 | -0.13 | 17.2 | 17.07 | |
| 2048 | ZNF207 | O43670 | 0.96 | 0.017728767 | 0.06 | 14.91 | 14.97 | |
| 2049 | ZRANB2 | O95218 | 0.31 | 0.508638306 | 0.07 | 17.64 | 17.71 | |
| 2050 | ZW10 | O43264 | 0.18 | 0.744727495 | 0.12 | 14.53 | 14.65 | |
| 2051 | ZYX | Q15942 | 0.0001 | 4 | 0.55 | 16.19 | 16.75 | |
